# Cysteine enrichment mediates co-option of uricase in reptilian skin and transition to uricotelism

**DOI:** 10.1101/2023.06.02.543418

**Authors:** Giulia Mori, Anastasia Liuzzi, Luca Ronda, Michele Di Palma, Magda S. Chegkazi, Soi Bui, Mitla Garcia-Maya, Jasmine Ragazzini, Marco Malatesta, Emanuele Della Monica, Claudio Rivetti, Parker Antin, Stefano Bettati, Roberto A. Steiner, Riccardo Percudani

**Affiliations:** Department Of Chemistry, Life Sciences And Environmental Sustainability, University of Parma, Parma, Italy; Department of Medicine and Surgery, University of Parma, Parma, Italy; Department of Biomedical Sciences, University of Padova, Padova, Italy; Randall Centre of Cell and Molecular Biophysics, King’s College London, London, UK; Department of Cellular and Molecular Medicine, University of Arizona, Tucson, US

## Abstract

Uric acid is the main means of nitrogen excretion in uricotelic vertebrates (birds and reptiles) and the end product of purine catabolism in humans and a few other mammals. While uricase is inactivated in mammals unable to degrade urate, the presence of orthologous genes without inactivating mutations in avian and reptilian genomes is unexplained. Here we show that the *Gallus gallus* gene we name cysteine-rich urate oxidase (CRUOX) encodes a functional protein representing a unique case of cysteine enrichment in the evolution of vertebrate orthologous genes. CRUOX retains the ability to catalyze urate oxidation to hydrogen peroxide and 5-hydroxyisourate (HIU), albeit with a 100-fold reduced efficiency. However, differently from all uricases hitherto characterized, it can also facilitate urate regeneration from HIU, a catalytic property which we propose depends on its enrichment in cysteine residues. X-ray structural analysis highlights differences in the active site compared to known orthologs and suggests a mechanism for cysteine-mediated self-aggregation under H_2_O_2_-oxidative conditions. Cysteine enrichment was concurrent with transition to uricotelism and a shift in gene expression from the liver to the skin where CRUOX is co-expressed with β-keratins. Therefore, the loss of urate degradation in amniotes has followed opposite evolutionary trajectories: while uricase has been eliminated by pseudogenization in some mammals, it has been repurposed as a redox-sensitive enzyme in the reptilian skin.

## Introduction

Uricase or urate oxidase (Uox) is an evolutionary conserved enzyme of ancient origin that is found in all three domains of life. In vertebrates, it acts in the liver peroxisomes (Hayashi et al. 2000) by converting the purine catabolic product uric acid to 5-hydroxyisourate (Kahn and Tipton 1998; Bui et al. 2014). Depending on the species, purine degradation proceeds further to give allantoin or urea (Noguchi et al. 1979; Dembech et al. 2023) as nitrogen waste products that are excreted from the body through urine. In various species, however, the end product is the poorly soluble uric acid.

Truncation of purine degradation to uric acid coupled with de novo purine synthesis allows uricotelic animals to eliminate nitrogenous waste through minimal water consumption (Campbell et al. 1987; Wright 1995; Salway 2018). Among vertebrates, uricotelism is a monophyletic trait of sauropsids (birds and reptiles), which excrete both amino acid and purine nitrogen in the form of uric acid. Although uric acid is the end product of purine degradation also in some mammals, including humans, these organisms excrete mainly urea (ureotelism). Fish and amphibians, on the other hand, excrete mainly ammonia (ammonotelism), as do invertebrate deuterostomes (**Fig 1**).

**Fig 1.**
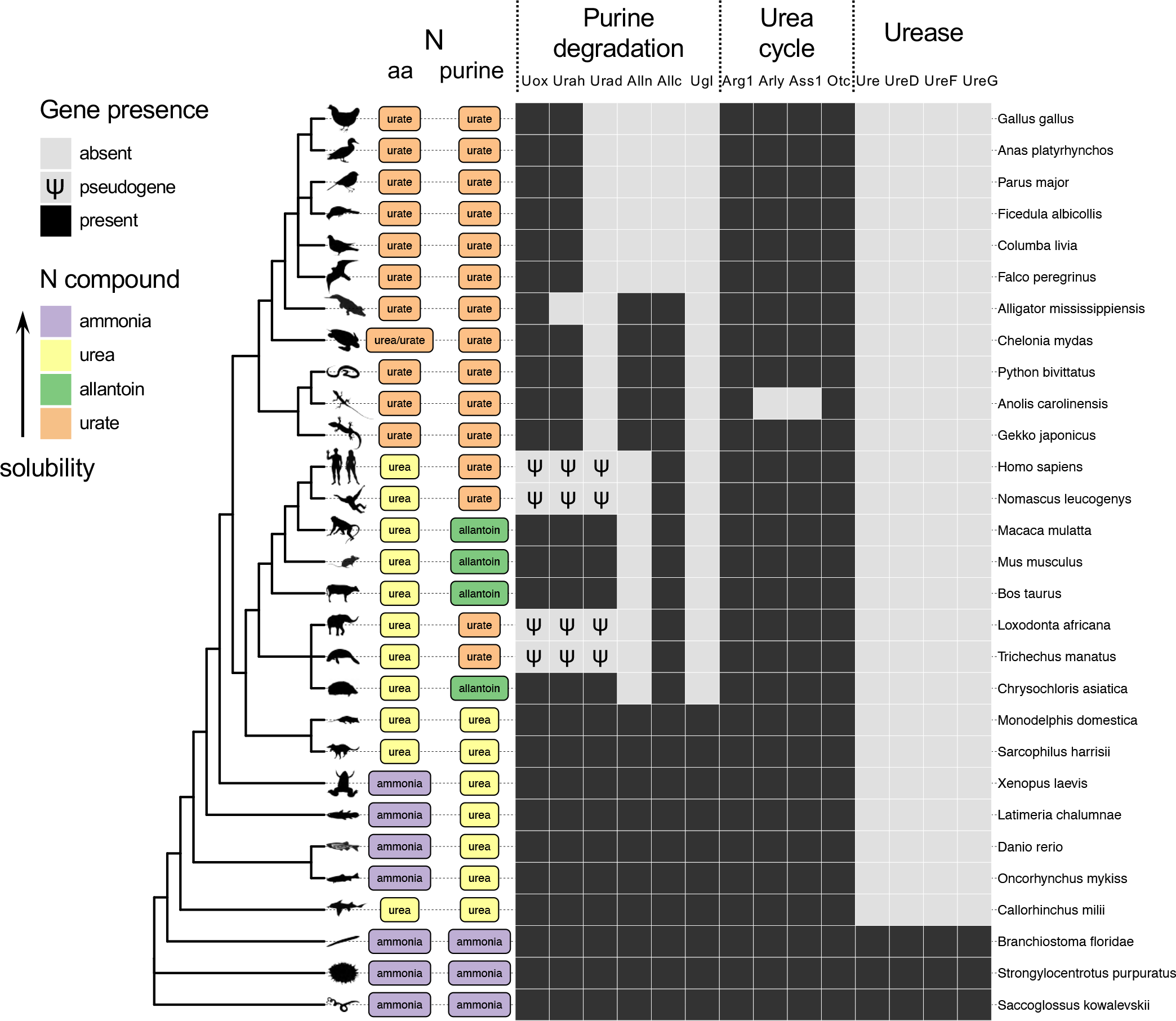
Nitrogen elimination in aquatic and terrestrial deuterostomes. Scheme of the main nitrogenous end product of amino acid and purine metabolism in vertebrate and invertebrate deuterostomes across the ncbi phylogeny of selected species with emphasis on vertebrates. The presence of genes relevant for the metabolism of urea and uric acid is indicated by black boxes. Pseudogenes are indicated by a Ѱ symbol. The main nitrogenous waste product of amino acid and purine metabolism is indicated based on literature evidence and/or genetic content (S1 File). PhyloPic silhouettes (http://phylopic.org) are added to selected branches to aid species identification.

The different strategies of nitrogen elimination in vertebrates exemplify the metabolic adaptations associated with different living environments. Nitrogen disposal in the form of ammonia can be accomplished by aquatic vertebrates as large amounts of water are required to maintain ammonia concentrations below toxic levels. Vertebrates that tolerate a dry environment convert ammonia into compounds (urea or uric acid) that can be more concentrated than ammonia in body fluids. Excretion of urea requires about ten times less water than ammonia, whereas the highly insoluble uric acid requires about 50 times less water (Wright 1995). Indeed, ureotelism and uricotelism have been two routes of adaptation to terrestrial life; in particular, uricotelism contributed to the radiation of reptiles during the Triassic Period providing them with the ability to tolerate the extremely dry Triassic climate (Campbell et al. 1987). It has been noted that this advantage comes at the expense of energy consumption, since about four phosphoanhydride ATP bonds are required for each nitrogen atom to convert ammonia to uric acid, compared with two required for its conversion to urea (Salway 2018). Even though vertebrates are unable to recycle nitrogen contained in urea or uric acid, this capacity can be provided by intestinal microbes that degrade urea and uric acid to generate ammonia as a nitrogen source. This nitrogen conservation mechanism may be relevant in conditions of limited nitrogen intake (Singer 2003).

In addition to water conservation, the truncation of purine degradation to uric acid, that ultimately led to the development of uricotelism in sauropsids, is thought to have provided at least one more selective advantage, due to the antioxidant properties of uric acid (Simic and Jovanovic 1989). The atmosphere at the time of reptiles radiation was more strongly oxidizing than is today and sauropsids, having a blood content of uric acid more than five-fold higher than that of ureotelic vertebrates (Dessauer 1970), may have had a selective advantage under those conditions. It has been suggested that the antioxidant properties of uric acid are the reason why the loss of uricase occurred in apes, especially after the loss in Haplorhini primates of the ability to synthesize the antioxidant ascorbic acid (Ames et al. 1981; Nishikimi et al. 1994). An evolutionary cost of high levels of uric acid is the formation of crystalline precipitate in the kidney and joints and the development of a painful arthritis known as gout. Similarly to humans, birds and reptiles suffer gout (Peterson et al. 1971; Rothschild et al. 2013), a disease which also affected dinosaurs according to the fossil record (Rothschild et al. 1997).

Truncation of purine degradation in apes was caused by *Uox* pseudogenization through nonsense mutations (Wu et al. 1989; Oda et al. 2002), possibly preceded by a missense mutation increasing the enzyme K_M_ for the urate substrate (Kratzer et al. 2014; Marchetti et al. 2016; Li et al. 2022). These events were accompanied by pseudogenization of the other two genes involved in the conversion of uric acid to allantoin, HIU hydrolase (*Urah*) and OHCU decarboxylase (*Urad*), through inactivating mutations of the coding sequence and transcriptional promoter (Ramazzina et al. 2006; Keebaugh and Thomas 2010). A similar pattern of pseudogenization (**Fig 1**), has been recently identified in elephants and manatee (Tethytheria), suggesting convergent loss of urate degradation in these mammals (Sharma and Hiller 2020). By contrast, although liver uricase is absent in uricotelic organisms, no inactivating mutations are found in the *Uox* gene, which appears to be universally conserved in sauropsids (**Fig 1**). This conservation suggests that the gene is still subjected to selection pressure, as also supported by a low Ka/Ks ratio (Keebaugh and Thomas 2010). Overall, these lines of evidence point to the possibility that reptilian uricase has been repurposed for a different function concurrently with transition to uricotelism, more than 300 MYA. Gene co-option, i.e. the establishment of a new use for an existing gene, is known to be an important mechanism in vertebrate evolution (True and Carroll 2002; McLennan 2008), but whether it has a role in the transition to uricotelism and maintenance of uricase in sauropsids is unknown.

To understand the biological significance of uricase maintenance in sauropsids, we studied the gene and protein from the model organism *Gallus gallus*. We found evidence that uricase has been repurposed as a redox sensitive enzyme in the skin of uricotelic vertebrates. By a large-scale analysis, we also found that co-option of uricase in the reptilian ancestor was mediated by an increase in the cysteine content that is unmatched in the evolution of vertebrate orthologous genes.

## Results

### Distinct sequence alterations in reptilian Uox genes

Alignment of Uox sequences (**Fig 2A**) revealed conservation of the active site residues in reptilian proteins with a single exception: a tyrosine substitutes a valine in position 230 of *Gallus gallus* Uox (*Gg*Uox). In contrast, the N and C termini are visibly different between reptilian and non-reptilian Uox. Differences in the N-terminal regions are due to modification in the exon-intron structure of vertebrate *Uox* genes (**Fig 2B**). In spite of the conserved synteny between *Uox* and *Dnase2b* in vertebrates (**Fig 2B**), it was previously noted that the first Uox-coding exon mapped to a distinct location in some birds (Keebaugh and Thomas 2010). Although the amino acid sequence encoded by the first exon is not conserved between avian and non-avian reptiles (**Fig 2A**), the exon-intron structure of the *Uox* locus in reptiles is the same, wherein the first exon lies upstream of the neighbor Dnase2b gene (first two species in **Fig 2B**). Even in cases (e.g. *Anolis carolinensis*) in which the transcription start site of *Uox* is reported downstream of *Dnase2b*, the existence of the upstream exon could be demonstrated by the alignment of the EST sequences (**S1 Fig**). Therefore, the first exon of vertebrate *Uox*, which lies downstream of the bidirectional promoter shared with the divergently oriented *Dnase2b* gene (last two species in **Fig 2B**), was probably lost in the reptile ancestor and a new one was acquired upstream the 3’ end of *Dnase2b*, together with a new promoter. At the C terminus, the peroxisome targeting signal (PTS1) present in eukaryotic Uox is absent owing to the addition of an extra peptide of ∼20 amino acids (**Fig 2A**). This suggests that Uox is not a peroxisomal protein in reptiles.

**Fig 2.**
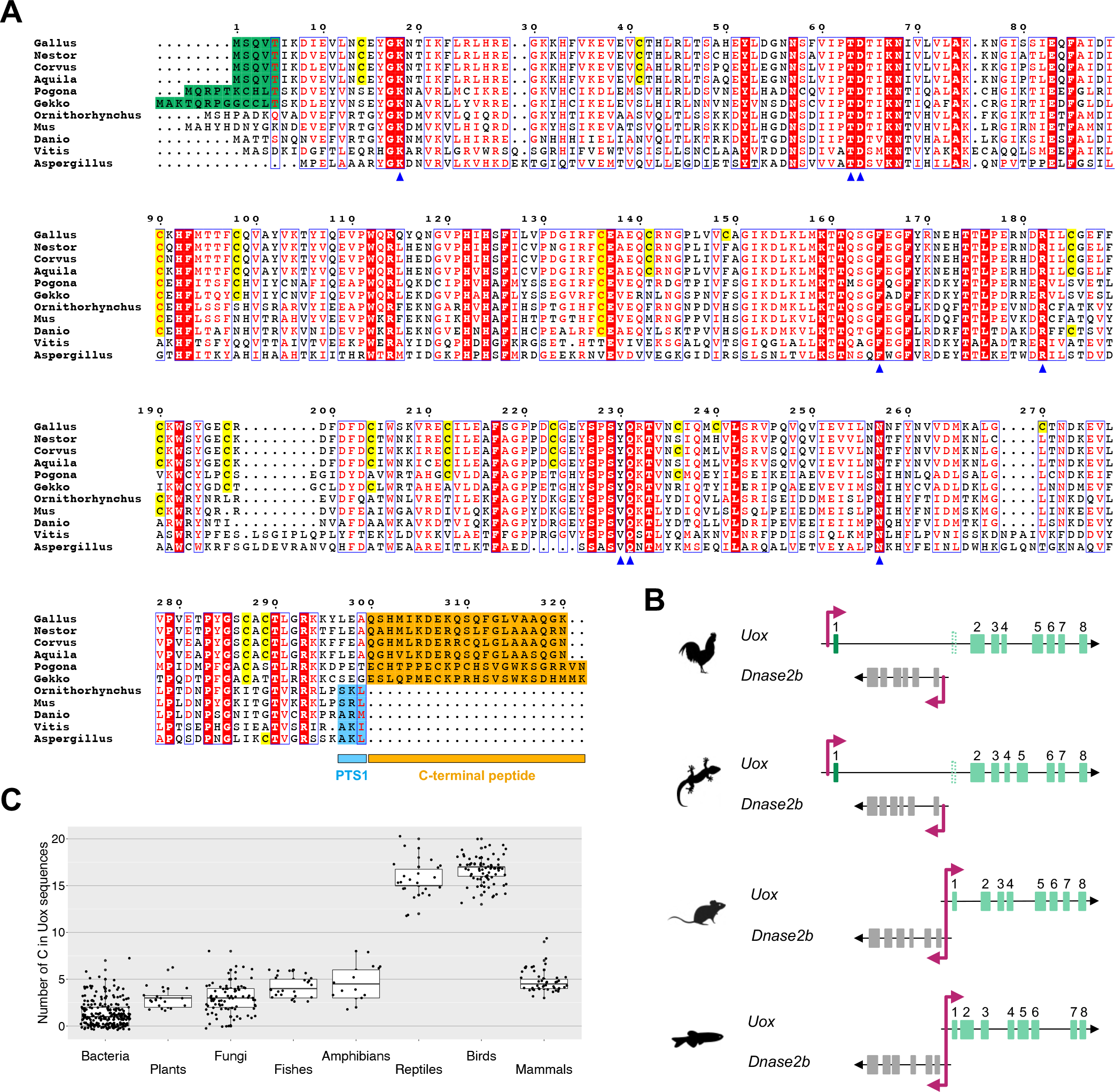
Distinct sequence alterations in reptilian Uox genes. (A) Multiple alignment of Uox sequences from different species, showing conservation of active site residues (blue arrowheads) together with diversification in cysteine content and in the C-terminal region. Cysteines of *Gallus gallus* (*Gg*) Uox and cysteines that are conserved in the same positions in the other sequences are highlighted in yellow. The N-terminal sequence encoded by the first exon in reptilian Uox is highlighted in dark green; the C-terminal peptide in reptilian Uox is highlighted in orange; the tripeptide corresponding to PTS1 in non-reptilian Uox is highlighted in light blue. (B) Gene structure of *Uox* and *Dnase2b* from *Gallus gallus* (*Uox*: LOC101747367), *Gekko japonicus* (*Uox*: LOC107117302), *Mus musculus*, and *Danio rerio*, showing absence of a bidirectional promoter in sauropsids, and presence in the other vertebrates. (C) Boxplots illustrating cysteine abundance in Uox sequences of different taxonomic groups. Median (thick lines), first and third quartile (thin lines) of the number of cysteines in individual Uox sequences (black circles) grouped according to taxonomy. Here, the term “reptiles” refers to non-avian reptile species.

A striking difference is found in the content of cysteine residues, that in reptilian Uox is about five-fold higher than that of non-reptilian Uox (∼5% vs ∼1%), including metazoan, fungal, plant, and bacterial sequences (**Fig 2C**). Cysteines appear randomly distributed along the reptilian Uox sequence and not always conserved across species (**Fig 2A**).

### Unique increase in cysteine content in reptilian Uox

Intrigued by the enrichment of cysteines in reptilian Uox sequences, we analyzed the variation in amino acid composition of orthologous proteins in vertebrates. We used a large set of orthologous groups (orthogroups) retrieved from the database OrthoDB, which was filtered to include only single-copy genes and orthogroups present in at least 90% of genomes. The selected dataset contained 3,393 orthogroups and 745,945 sequences distributed over three vertebrate clades: Sauropsida, Mammalia, and Actinopterygii. For the three taxonomic groups the mean content of each amino acid in each orthogroup was analyzed through pairwise comparisons (Sauropsida-Mammalia; Sauropsida-Actinopterygii; Mammalia-Actinopterygii).

Orthogroups with significant variation in amino acid content were identified by using a p-value cutoff of 1e-16 and a fold-change cutoff of ±1.

Only a few orthogroups were found to be significantly differentiated in cysteine content between the three group pairs: three between Sauropsida and Mammalia (**Fig 3A**), nine between Sauropsida and Actinopterygii (**Fig 3B**), and eight between Mammalia and Actinopterygii (**S2A Fig**, **S1 Table**). Interestingly, the Uox orthogroup stands out as an exceptional case of cysteine enrichment in the evolution of orthologous genes in amniotes (**Fig 3A** and **3B**, arrows).

**Fig 3.**
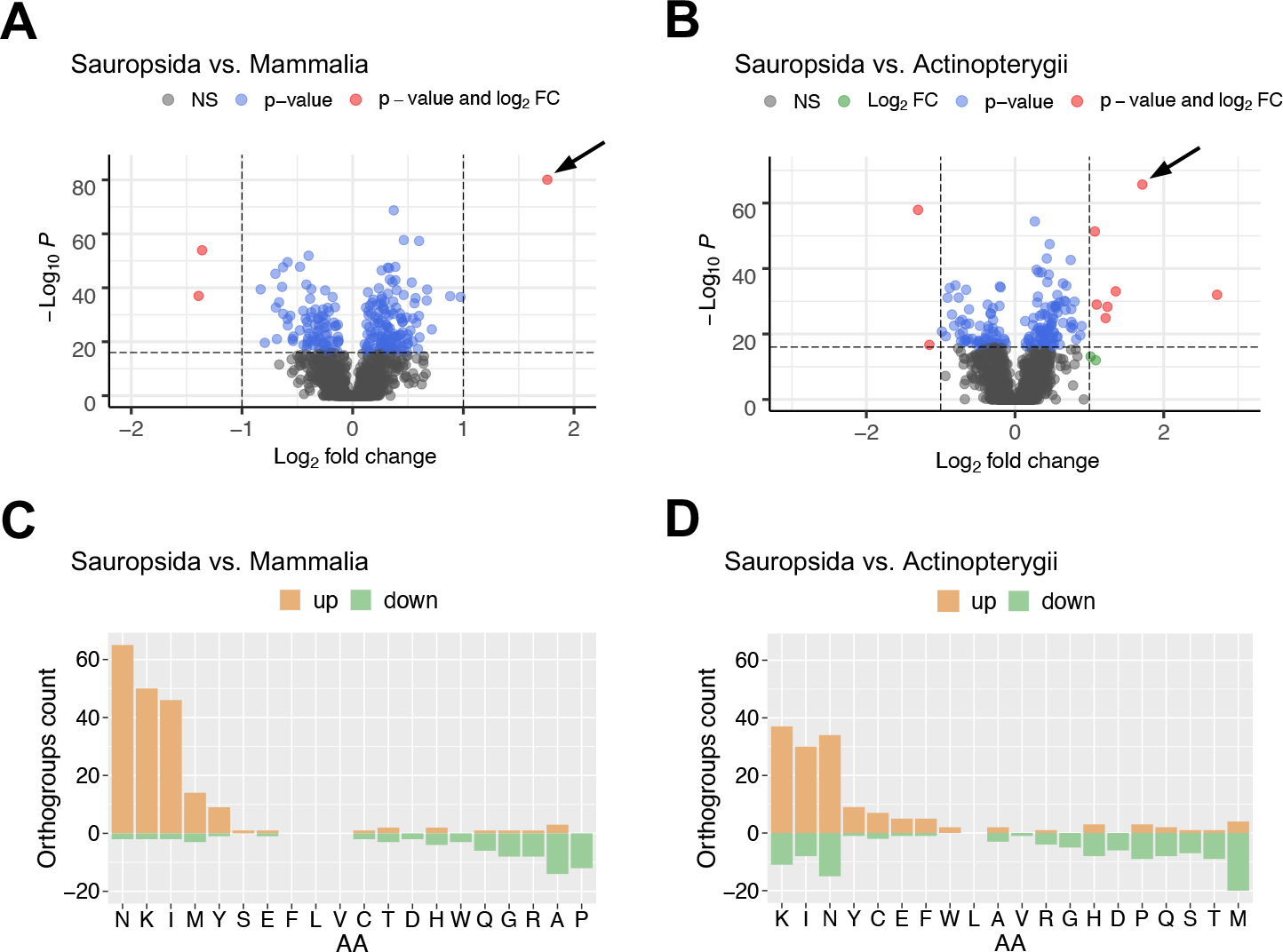
Amino acid content variation in vertebrate orthologous proteins. (A-B) Volcano plot depicting the cysteine content variation of orthologous proteins between groups of vertebrates: (A) sauropsids vs. mammals, and (B) sauropsids vs. fishes. Horizontal dashed line indicates p-value of 10^-16^; vertical dashed lines indicate log_2_ fold change of −1 and 1. The Uox orthogroup in which the cysteine content is significantly increased is indicated by an arrow. Orthogroup names and numeric values are reported in **S1 Table**. (C-D) Bar plot illustrating the number of orthogroups in which the content of the corresponding amino acid is significantly increased (light orange) or decreased (green) according to p-value < 10^-16^ and log_2_ fold change > ±1 in (C) sauropsids vs. mammals, and (D) sauropsids vs. fishes.

In our analysis we observed that glycine, leucine, valine, phenylalanine, tryptophan, glutamine, serine, threonine, tyrosine, aspartate, glutamate, arginine and histidine followed the same trend of cysteine, as they have emerged as significantly varying in a few (<10) or no orthogroups across the three vertebrate clades (**Figs 3C**, **3D** and **S2B**). In contrast, isoleucine, asparagine, and lysine (INK) appeared as significantly varying in a greater number of orthogroups (>50) in each of the three pairwise comparisons. In particular, the three amino acids are increased in reptiles (**Figs 3C**, **3D** and **S3**); asparagine is also decreased in fishes with respect to mammals (**S2B Fig**). Of the remaining amino acids, methionine is increased in some piscine orthogroups (**Figs 3D** and **S2B**); alanine and proline are increased in some mammalian orthogroups (**Figs 3C** and **S2B**).

### *Gg*Uox catalyzes the oxidation of urate and the reduction of 5-hydroxyisourate

To investigate the activity of reptilian uricase, we overexpressed the coding sequence of *Gallus gallus* Uox (*Gg*Uox) in *E. coli* and purified the corresponding protein to apparent homogeneity using either xanthine-agarose (**Fig 4A**) or nickel affinity columns (**S4A Fig**). The binding to a stationary phase with a specific affinity for uricase (Nishimura et al. 1982) provides evidence that *Gg*Uox has a preserved purine-binding ability. The purified protein eluted as a single peak compatible with a tetramer in size-exclusion chromatography (SEC) (**S4B Fig**). The enzyme catalyzed the oxidation of urate to 5-hydroxyisourate (HIU) and hydrogen peroxide and exhibited Michaelis-Menten kinetics at increasing substrate concentrations (**Fig 4B**). However, despite a K_M_ (12.5 μM) comparable to that of *Danio rerio* Uox (*Dr*Uox) (11 μM) (Marchetti et al. 2016), the turnover number (k_cat_) is >100-fold lower (0.03 s^-1^ for *Gg*Uox vs 3.95 s^-1^ for *Dr*Uox). Consistently, the kinetics of hydrogen peroxide formation coupled to the oxidation of urate in single turnover conditions is slower in the presence of *Gg*Uox than in the presence of *Dr*Uox (**Figs 4C** and **S4C**).

**Fig 4.**
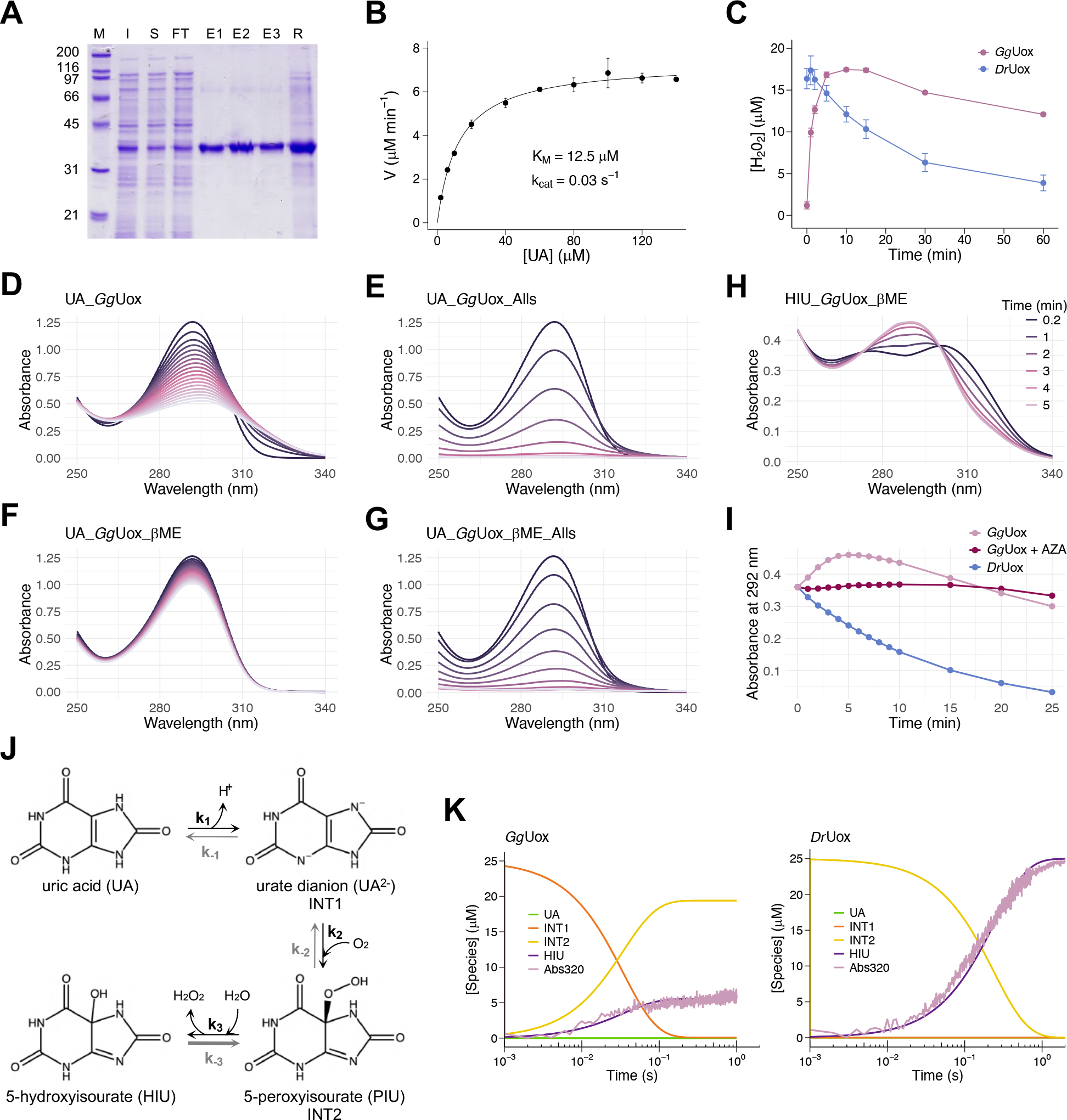
Uricase activity of *Gallus gallus* Uox. (A) SDS-PAGE of the purification of *Gg*Uox by xanthine-agarose affinity chromatography. M: marker; I: post-induction total cell fraction; S: soluble cell fraction; FT: flow-through; E1-E3: elution fractions; R: resin. (B) Michaelis-Menten dependencies of *Gg*Uox uricase activity on uric acid (UA) concentration. Kinetic measurements were carried out in 100 mM potassium phosphate (pH 7.6), at 25 °C, using 4.1 µM *Gg*Uox and 1 µM HIU hydrolase (HIUase). (C) Kinetics of hydrogen peroxide turnover in the urate oxidation reaction catalyzed by *Gg*Uox or *Dr*Uox. Reactions were carried out under single turnover conditions with 100 mM potassium phosphate (pH 7.6), 30 μM enzyme, 25 μM UA. The amount of hydrogen peroxide was quantified by Ferrous Oxidation Xylenol Orange (FOX) assay. (D-G) Time-resolved UV-Vis spectra of urate oxidation by *Gg*Uox. (D) Reaction mixture contained 100 mM potassium phosphate (pH 7.6), 30 μM *Gg*Uox, and 100 μM UA. (E-G) Reaction mixtures were the same as in (D) with the addition of (E) 1.25 μM allantoin synthase (Alls), (F) 300 μM β-mercaptoethanol (βME), (G) 1.25 μM Alls and 300 μM β-ME. Spectra were acquired every 1 min at 25 °C. (H) Time-resolved UV-Vis spectra of of 5-hydroxyisourate (HIU) degradation by *Gg*Uox. Reaction contained 100 mM potassium phosphate (pH 7.6), 60 μM HIU, 30 μM *Gg*Uox, and 300 μM βME. Spectra were acquired every 1 min at 25 °C. (I) Time-dependent change of absorbance at 292 nm during HIU degradation by *Gg*Uox (pink), *Gg*Uox with 8-azaxanthine (AZA; bordeaux), and *Dr*Uox (light blue). Reaction mixtures were the same as in (H); AZA was 50 μM. (J) Three-steps reversible model for UA to HIU conversion. Intermediate species were already attributed to urate dianion (UA^2-^) and 5-peroxyisourate (PIU) (Kahn and Tipton 1998). (K) Experimental stopped flow kinetics (“Abs320”: pink) of *Gg*Uox (left) or *Dr*Uox (right) reaction and fitting to three-steps reversible model for UA to HIU conversion. Reactions were carried out under single turnover conditions at 25 °C with 100 mM potassium phosphate (pH 7.6), 35 μM enzyme, 25 μM UA.

Accumulation of the unstable oxidation product, HIU, can be detected as a shift of the urate peak from 292 to 301 nm during the *Dr*Uox reaction (**S5A Fig**). In the case of *Gg*Uox, HIU accumulation was less noticeable (**Fig 4D**), but when an enzyme with HIU hydrolase activity (allantoin synthase; Alls) was added, the reaction reached completion at an initial rate about three-fold faster (**Fig 4E**). This behavior suggests a possible reconversion of HIU to urate that has been described for uricase catalysis in the presence of a reducing agent, such as DTT or cysteine (Sarma and Tipton 2000). When the *Gg*Uox reaction was conducted in the presence of β-mercaptoethanol (βME), the oxidation of urate to HIU was extremely slow (**Fig 4F**), and, again, the addition of Alls resulted in the completion of the reaction at an initial rate comparable to that in the absence of the reducing agent (**Fig 4G**). In contrast, the *Dr*Uox activity under the same reducing conditions was almost unaffected (**S5B Fig**). This suggests that βME, while unable to reduce HIU, can reduce *Gg*Uox cysteines that are significantly more abundant than in *Dr*Uox. To confirm the reconversion reaction, we isolated enzymatically generated HIU and monitored its reduction back to urate in the presence of either *Gg*Uox and βME (**Fig 4H**) or *Dr*Uox and βME (**S6A Fig**). Only with *Gg*Uox we observed the shift of the HIU peak to 292 nm, indicating the formation of urate; with *Dr*Uox, HIU degraded spontaneously in a time-dependent manner similarly as without enzymes (**S5B Fig**). The recycling of urate during *Gg*Uox (and not *Dr*Uox) reaction may be explained with the increased cysteine content of *Gg*Uox.

To assess the involvement of *Gg*Uox active site residues in the reduction of HIU, we repeated the experiment in the presence of 8-azaxanthine (AZA; **S6C Fig**), a competitive inhibitor of uricase activity (Heitaroh et al. 1973) acting on *Gg*Uox as well (**S7 Fig**). In this case, the reconversion of HIU to urate is slower because, despite the shift of the HIU peak to 292 nm (**S6C Fig**), we did not observe a similar increase of absorbance at 292 nm as in the reaction without the inhibitor (**Fig 4I**). However, the disappearance of HIU, as monitored by absorbance decrease at 315 nm, is faster than that in the presence of *Dr*Uox (**S6D Fig**). These results suggest that the enzymatic regeneration of urate from HIU depends at least in part on the oxidoreductase activity at the *Gg*Uox active site.

*Gg*Uox activity was also investigated following urate degradation under single turnover conditions (25 μM urate in the presence of excess *Gg*Uox - 35 μM) upon rapid mixing with a stopped flow apparatus; the same experiment was carried out with *Dr*Uox for comparison. In agreement with the results described above, the observed kinetics indicated clear differences between *Gg*Uox and *Dr*Uox reactions. We observed that only ∼20% HIU is formed in the presence of *Gg*Uox compared to *Dr*Uox (**S8A Fig**). Singular value decomposition (SVD) on data matrices of kinetic traces for both enzymes showed a first spectral component, accounting for the large majority of the signal, with a negative peak centered at 294 nm and a positive peak at 314 nm (**S8B Fig**, blue and light blue lines). This analysis allowed us to define a wavelength (320 nm) where only this component contributes to the kinetics (**S8B Fig**, orange and yellow lines). Single wavelength kinetics at 320 nm were then evaluated in terms of a three-step sequential model previously proposed for uricase-catalyzed reaction (Kahn and Tipton 1998), considering reversible (**Fig 4J**) or irreversible (**S9A Fig**) processes. In this model, the first intermediate is a urate dianion and the second intermediate is 5-peroxyisourate (PIU).

Noteworthy, based on the previously reported spectra of Uox reaction intermediates (Kahn and Tipton 1998), the first SVD component corresponds to the difference spectrum between PIU and HIU (**S8C Fig**). We confidently attribute the time-evolution of this SVD component to this direct conversion. For *Dr*Uox, irreversible steps were successfully applied to fit the experimental data (**Fig 4K**), indicating that the forward reaction rates are much faster than the reverse reaction rates (**S9B Fig**). We observed that while urate oxidation promptly proceeds to PIU, as the concentration of this second intermediate is maxima at the very beginning of the reaction (**Fig 4K**, yellow line in right panel), the final conversion to HIU is the rate limiting step, with a rate constant (4.11±0.02 s^-1^) similar to the determined k_cat_ (3.95 s^-1^). Because with *Gg*Uox the reaction did not reach completion (**Figs 4K**, left panel and **S8A**), reversible steps were introduced (**Fig 4J**). Indeed, a reaction model with three reversible steps well describes the experimental kinetics (**Fig 4K**, left panel). The last reaction step involves the reconversion of HIU to PIU (*k*_‒3_>*k*_3_) accounting for an equilibrium concentration of HIU never reaching the starting urate levels. Once hydrogen peroxide is formed, its concentration continues to be high in *Gg*Uox reaction compared with *Dr*Uox (**Figs 4C** and **S4**), suggesting that the last step involves the reduction of HIU without H_2_O_2_ consumption.

### X-ray crystallographic analysis of Cys-rich *Gg*Uox

As *Gg*Uox displays unconventional catalytic properties, we pursued its X-ray crystallographic analysis. We crystallized *Gg*Uox in complex with its AZA inhibitor in two space groups (*P*2_1_2_1_2_1_ and *C*222_1_) and solved its structure at resolution values between 1.7 and 2.1 Å (see **S2 Table**). In the crystal (**Fig 5A, B**), *Gg*Uox is a tetramer (dimer of dimers) as expected from SEC (see **S4B Fig**) and observed in other species (Colloc’h et al. 1997; Kratzer et al. 2014; Hibi et al. 2016; Marchetti et al. 2016; Chiu et al. 2021). Electron density is generally well defined, but flexibility prevented modelling of the C-terminal region corresponding to the unique reptilian peptide extension (last 20 amino acids; see **Fig 2A**). The structure of *Gg*Uox is very similar to those of uricases from other organisms with typical rms deviations as assessed using the PDBeFold server (Krissinel and Henrick 2004) around 1.50 Å – the fossil euarchontoglires uricase (Kratzer et al. 2014) is the most similar one (PDB code 4MB8; r.m.s.d. 1.16 Å).

**Fig 5.**
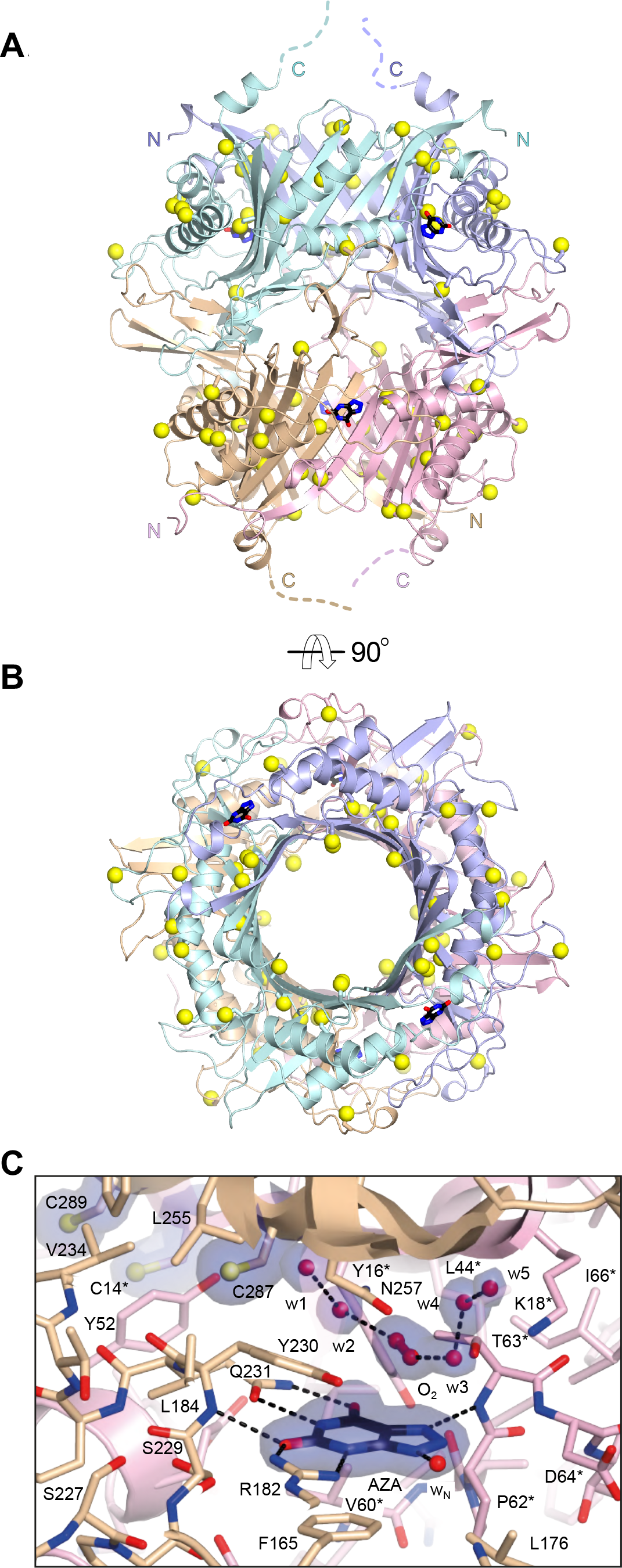
X-ray structure of Cys-rich *Gg*Uox. (A-B) Side (A) and top (B) cartoon representations of the 146.5 kDa *Gg*Uox tetramer with the four protomers highlighted by different colors. The N and C termini of the different chains are indicated by the letters N and C, respectively. The flexible C-terminal regions corresponding to the last 18-20 amino acids are indicated in (A) by broken lines. Bound 8-azaxanthine (AZA) ligands at the dimers’ interface are shown as sticks. Cysteine residues are also shown as sticks with their SG atoms highlighted by yellow spheres. (C) Stick representation of AZA (black) bound in the active site at the interface between two different *Gg*Uox protomers, shown in wheat and pink for chain A and chain B, respectively. O_2_ and water molecules (labelled w1-5) closest to AZA are represented by spheres. 2*mF*_o_-*DF*_c_ electron density map for AZA, cysteines, and the solvent above the inhibitor is shown in blue at the 1.0s level. Residues labelled with an asterisk indicate that they belong to a different Uox protomer. Cysteine residues in proximity are also shown with their SG atoms highlighted by spheres. Nitrogen, oxygen, and sulfur atoms are in blue, red, and yellow, respectively. Hydrogen bonds are shown by broken black lines.

Individual protomers are composed of a tandem of structurally similar tunneling-fold domains of mixed ɑ+β topology that form an antiparallel curved β-sheet with helices arranged on the convex side of the sheet. Dimerization engenders a basket-like structure that hosts two symmetric active sites contributed by residues from both chains. Recruitment of an additional dimer positioned upside-down on top of the first one and rotated by 90° generates the complete tetrameric assembly of D2 symmetry. Reptilian uricase displays a unique enrichment in cysteines (see **Figs 2** and **3**). **Figure 5A** and **5B** shows that cysteine residues mostly decorate the central section of the *Gg*Uox dimer where they are fairly homogeneously distributed.

AZA binding to *Gg*Uox is stabilized by direct hydrogen bonds with main-chain nitrogen atoms of T63* and Y230 as well as with the side chains of R182 and Q231, and a water molecule (wN) (**Fig 5C**, the asterisk indicates residues belonging to a different protomer). Residues Y16*, P62*, D64*, F165, L176, and S229 further line the binding pocket with the aromatic side-chain of F165 engaged in π-stacking with the pyrimidine ring of the inhibitor. At different stages of the uricase reaction O_2_ and water must occupy the ‘peroxo hole’ above the organic substrate in proximity of the Thr-Lys catalytic dyad (T63*-K18* in *Gg*Uox) (Bui et al. 2014; McGregor et al. 2021). In the present structures, electron density at this location is best explained by a mixture of water molecules and O_2_ connected via hydrogen bonds that are generally conserved in the different *Gg*Uox active sites (**Figs 5C** and **S10**). Dioxygen interacts via H-bonds with the nearby T63* and its stabilization is likely also promoted by the conformation of the sauropsid-specific Y230 that lines the ‘peroxo hole’ with the hydrophobic ‘side’ of its phenolic side chain. Of the eighteen cysteine residues of *Gg*Uox, C287, positioned directly above Q231, is in closest proximity of the ‘peroxo hole’ whilst C14* and C289 are further away at ∼8.5 Å from C287. All these cysteines are sauropsid-specific and conserved (see **Fig 2A**).

### *Gg*Uox aggregation upon catalysis

In addition to those decorating the central section of the *Gg*Uox dimer, a number of cysteines are distributed on the protein surface (see **F**ig **5A** and **5B**). These exposed cysteines could be susceptible to H_2_O_2_ oxidation during *Gg*Uox reaction suggesting possible interactions between tetramers. To test this hypothesis, we measured the hydrodynamic diameter of *Gg*Uox in solution during the reaction by dynamic light scattering (DLS). Measurements were done at different time points after the addition of the urate substrate. Initially, we employed reaction conditions similar to those used for the spectroscopic measurements, namely 100 μM urate and a substrate-enzyme ratio of 5:1. Under these conditions, no aggregation was observed over a 2-hour period (**S11A Fig**). However, when the concentration of urate was increased to 2.1 mM, with a substrate-enzyme ratio 100:1, the hydrodynamic diameter of GgUox increased from 12 to ∼700 nm after 5 minutes and stabilized at ∼500 nm within 30 minutes (**Fig 6A**, left panel). When *Gg*Uox was pre-incubated with βME, no change of the hydrodynamic diameter occurred over a 1-hour period (**Fig 6A**, right panel). By repeating the same experiments with *Dr*Uox we observed no changes in the hydrodynamic diameter either in the presence or in the absence of βME (**Fig 6B**). These results strongly suggest that, under our experimental conditions, *Gg*Uox aggregates during the reaction most probably through the formation of disulfide bonds.

**Fig 6.**
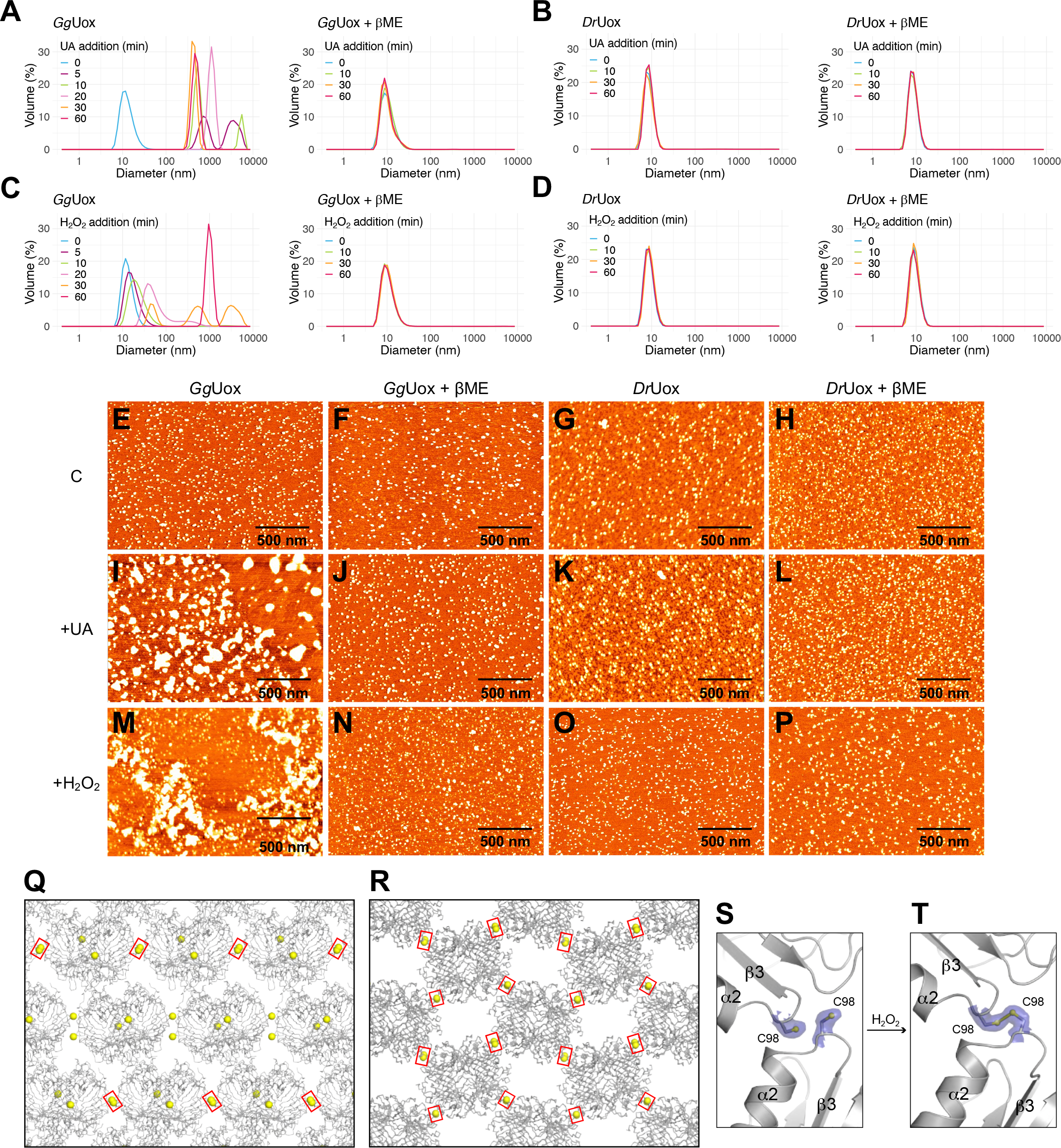
Self-aggregation behavior of *Gg*Uox. (A-B) DLS measurements of (A) *Gg*Uox or (B) *Dr*Uox samples containing 21 μM enzyme, and (left panel) 0 or (right panel) 300 μM β-mercaptoethanol (βME) incubated at RT and analyzed at different time points after the addition of 2.1 mM urate (UA). The protein-substrate solutions were diluted to a protein concentration of 7 μM before measurements. (C-D) DLS measurements of (C) *Gg*Uox or (D) *Dr*Uox samples containing 7 μM enzyme, and (left panel) 0 or (right panel) 300 μM β-mercaptoethanol (βME) incubated at RT and analysed at different time points after the addition of 100 μM H_2_O_2_. (E-H) AFM 2D images of (E) *Gg*Uox not pre-incubated or (F) pre-incubated with βME, and of (G) *Dr*Uox not pre-incubated or (H) pre-incubated with βME. (I-L) AFM 2D images of (I-J) *Gg*Uox or (K-L) *Dr*Uox after incubation at RT with 2.1 mM UA at a substrate/enzyme ratio of 100:1, and (I, K) 0 or (J, L) 300 μM βME. (M-P) AFM 2D images of (M-N) *Gg*Uox or (O-P) *Dr*Uox after incubation at RT with 100 μM H_2_O_2_ at a substrate/enzyme ratio of 70:1, and (M, O) 0 or (N, P) 300 μM βME. The protein-substrate solutions were diluted to a protein concentration of 100 nM before being deposited on the mica surface. Size bar, 500 nm. (Q-R) Packing arrangement for *Gg*Uox in space groups *C*222_1_ (Q) and *P*2_1_2_1_2_1_ (R) with surface-exposed C98 residues represented by yellow spheres. In the crystal, some, or all of them engage in intermolecular disulfide bridges highlighted by red rectangles. In *C*222_1_ these S-S bonds generate cross-linked linear filaments of *Gg*Uox whilst in *P*2_1_2_1_2_1_ they produce two-dimensional layers. (S-T) Representative C98-mediated intermolecular disulfide crosslinking triggered by H_2_O_2_ treatment. 2*mF*_o_-*DF*_c_ electron density map shown in blue at the 1.0s level. Going from the reduced (S) to the oxidized (T) state, the sidechain of C98 in the bottom *Gg*Uox molecule undergoes a ∼120° rotation from the *plus* to the *minus* conformer engendering a disulfide bond with a neighboring molecule.

To assess the oxidizing action of the uricase product hydrogen peroxide on Uox cysteines, we measured the particle diameter at different time points after the two enzymes were exposed to this compound. Aggregation of *Gg*Uox occurred only in the absence of βME (**Fig 6C**), whereas aggregation of *Dr*Uox occurred neither in the absence nor in the presence of βME (**Fig 6D**).

Once formed, *Gg*Uox aggregates were unaffected by the addition of βME (**S11B Fig**).

In addition to DLS, atomic force microscopy (AFM) was employed for the direct visualization of Uox aggregation. Firstly, *Gg*Uox and *Dr*Uox, under native or reducing conditions, were each deposited on a freshly-cleaved mica surface (**Fig 6E-6H**). The average particle volume measured by AFM was consistent with a tetrameric protein of ∼150 kDa. No significant difference in size was observed between samples pre-incubated or not with βME (**S12A-S12E Fig**). Then, the two enzymes were each incubated for 1 hour at room temperature with urate or hydrogen peroxide either in the presence or absence of βME and deposited onto freshly-cleaved mica (**Fig 6I-6P**). Consistently with DLS measurements, we observed the formation of large protein aggregates only in *Gg*Uox samples incubated either with urate or hydrogen peroxide in the absence of βME (**Figs 6I, 6M** and **S12F-S12G**).

The X-ray structure of *Gg*Uox suggests a possible mechanism for disulfide-mediated aggregation. *Gg*Uox was crystallized in two alternative space groups (*C*222_1_ and *P*2_1_2_1_2_1_) that display a different packing arrangement (**Fig 6S** and **6T**). However, in both space groups intermolecular contacts are, at least partly, mediated by the loop containing the sauropsid-specific C98 residue, indicating that this interaction is also favored in solution. Under reducing conditions all C98 residues are seen adopting the *plus* rotamer (chi1 = 55°) with SG atoms from neighboring ones typically ∼5.4 Å apart (**Fig 6U**). Upon treatment with H_2_O_2_ one cysteine rotates to its *minus* rotational conformer (chi1 = -65°) allowing these residues to engage in disulfide bridges (**Fig 6V**) that in space group *C*222_1_ generate linear filaments (**Fig 6S**) whilst in *P*2_1_2_1_2_1_ give rise to two-dimensional cross-linked arrays (Fig 6T). Exposure to H_2_O_2_ also selectively oxidizes other *Gg*Uox cysteine residues. Whilst most are unmodified, residues C41, C141, C197 and C204 that are sauropsid-specific (see **Fig 2A**) are generally seen in their sulfinic (Cys-SO_2_H) form, although in a few chains some of these are present as sulfenic acids (Cys-SOH), probably as intermediates to their higher oxidation state (**S13 Fig**).

### Uricase gene duplication in chelonian reptiles

The *Uox-Dnase2b* locus with a head-to-head arrangement is present in conserved synteny in metazoans (Mori et al. 2022). Among reptiles, chelonians (tortoises and turtles) have a duplicate *Uox*, which we call *Uox2*, and which lies tail-to-tail with *Dnase2b* (**S14A Fig**). The two paralogs are sister to each other in the Uox phylogeny (**S14B Fig**), suggesting that the duplication event occurred within the Testudines order.

Curiously, these paralogs differ at the very active site position that distinguishes reptilian and non-reptilian Uox: chelonian Uox sequences have a histidine at position 230 (relative to the *Gg*Uox sequence), whereas Uox2 maintained the tyrosine residue (**S15 Fig**). In contrast, non reptilian Uox have an aliphatic amino acid (in most cases valine) at that position (see **Fig 2A**). To evaluate the effect of these substitutions on uricase activity, we mutated the tyrosine in the active site of *Gg*Uox to either histidine (Y230H), as observed in chelonian Uox, or valine (Y230V), as in a canonical Uox sequence. For both mutants, the specific activity was decreased by ∼75% compared to that of the wild-type enzyme (**S14C Fig**), although no differences were observed in protein solubility (**S16 Fig**).

### Reptilian *Uox* is expressed in the skin and shows a punctuated pattern in feather epithelium

Available RNA-sequencing (RNA-seq) data from *Gallus gallus* and another bird, *Serinus canaria,* and from the lizard *Anolis carolinensis* were analyzed to explore the expression pattern of Uox in adult tissues (**S3 Table** and **S18 Fig**). In the two birds, we detected *Uox* expression in the skin (**Fig 7A** and **7B**), particularly in the feather follicle (**Fig 7A**), while in the lizard, *Uox* transcripts are abundant in regenerating tail tissues (**Fig 7C**). *Uox* transcripts are absent in the liver at variance with non-reptiles. By contrast, high expression in the liver is maintained by *Dnase2b* and *Urah* (**S18 Fig**). Expression of the two *Uox* paralogs of turtles was evaluated by analyzing RNA-seq data from the *Chelonia mydas* and *Pelodiscus sinensis* (**S4 Table**). *Uox1* and *Uox2* transcripts are both present in the skin, with higher levels for *Uox2*, and not in the liver or kidney at variance with Dnase2b and Urah (**S19 Fig**).

**Fig 7.**
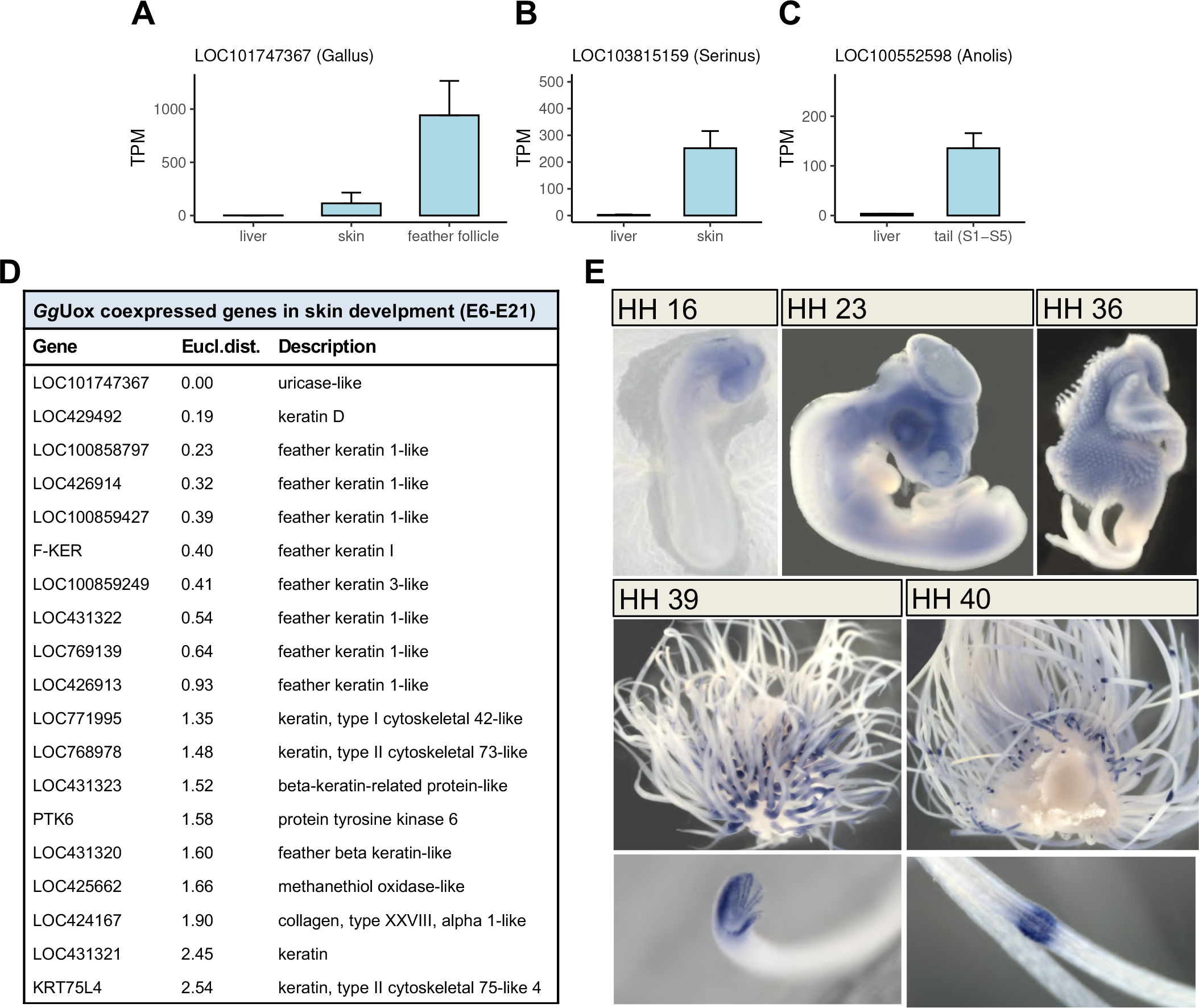
Expression of *Uox* in reptilian skin. (A-C) *Uox* gene expression levels (TPM: Transcripts Per kilobase Million) in the liver and tegumental tissues (skin and feather follicle) of (A) *Gallus gallus*, (B) *Serinus canaria*, and (C) *Anolis carolinensis*, as derived from RNA-seq data analysis. (D) List of genes included in the same cluster as *Gg*Uox gene by similarity in their expression profiles in skin embryo developmental stages E6-E21. (E) In situ hybridization analysis of UOX expression in chick embryos at HH developmental stages 16, 24, 36, 39 and 40.

By analyzing available data from the chicken skin transcriptome from day 6 to day 21 of embryogenesis (**S3 Table**), we could observe the temporal variation of UOX expression and clusterize UOX co-expressed genes based on their expression profiles (**S20 Fig**). The resulting cluster comprises mainly feather keratins (**Fig 7E**). Notably, the highest expression occurs at days 13-14, consistent with the increase of gene expression observed between days 10-15 by cap analysis of gene expression (CAGE) available on Chicken-ZENBU platform (Lizio et al. 2017) (**S21 Fig**).

To determine the pattern of UOX expression during embryogenesis at higher spatial resolution, whole-mount in situ hybridization analyses were performed in chicken embryos from day 2 to 15 of development corresponding to Hamburger–Hamilton (HH) stages 16-40 (Hamburger and Hamilton 1992) (**Fig 7D**). At HH stage 16, UOX expression was first detected in the head mesenchyme; at HH stage 23 it was found also in the leg and wing mesenchyme. In the next stages until HH stage 36, UOX mRNAs were further detected in the lung, heart, foregut, and spinal cord. At HH stages 39-40 a punctuated expression of UOX was detected in the feather epithelium. The punctuated expression of UOX during the last stages of embryo development coincides with the increase of gene expression observed by RNA-seq and CAGE analyses.

During this period, primary follicles form allowing differentiation and elongation of the feather filaments (Lucas and Stettenheim 1972). In parallel, the expression of URAH and DNASE2B genes was analyzed, but no staining was observed at HH stage 39 in the feather epithelium (**S22 Fig**).

Both RNA-seq and in situ hybridization data provide evidence that reptilian *Uox* lost expression in the liver and acquired expression in the skin, in particular in bird feather and lizard tail. *Gg*Uox ability to degrade urate was evaluated at slightly acidic pH values that mimic the skin environment. Although the enzyme is still active at lower pH, no increase in the specific activity was observed (**S23 Fig**).

## Discussion

Transition to uricotelism is a main vertebrate adaptation to life on land, accounting for the exquisite ability of birds and reptiles to survive in arid environments. A relevant question about the molecular basis of this physiological condition is the fate of uricase, the enzyme that in other vertebrates catalyzes conversion of uric acid to more soluble compounds. Our study reveals a history of co-option of the uricase gene in uricotelic vertebrates that is marked by an unparalleled increase in cysteine content. Functional divergence of orthologous genes at the sequence level is known to occur through substitutions of a few amino acid residues that are crucial for the protein function (Bartlett et al. 2003; Studer et al. 2013). Here, we show that a shift in the physiological role of uricase in reptiles occurred through a significant increase in the abundance of a particular amino acid (cysteine).

Through a large-scale analysis of the variation in amino acid composition among orthologs, we showed that in vertebrates the cysteine content tends to remain constant during evolution, and that Uox represents a unique case of cysteine enrichment across the two amniote lineages, sauropsids and mammals. Interestingly, the majority of cysteines were introduced in the Uox sequence after sauropsids split from stem amniotes, but before the divergence between lepidosaurs (lizards and snakes) and archelosaurs (turtles, birds, and crocodilians) (**Fig 8**). At this time, the evolutionary transition to uricotelism is thought to have already occurred (see **Fig 1**). Therefore, cysteine enrichment entailing the co-option of *Uox* gene is one of the molecular mechanisms that led to the establishment of uricotelism in the reptile ancestor.

**Fig 8.**
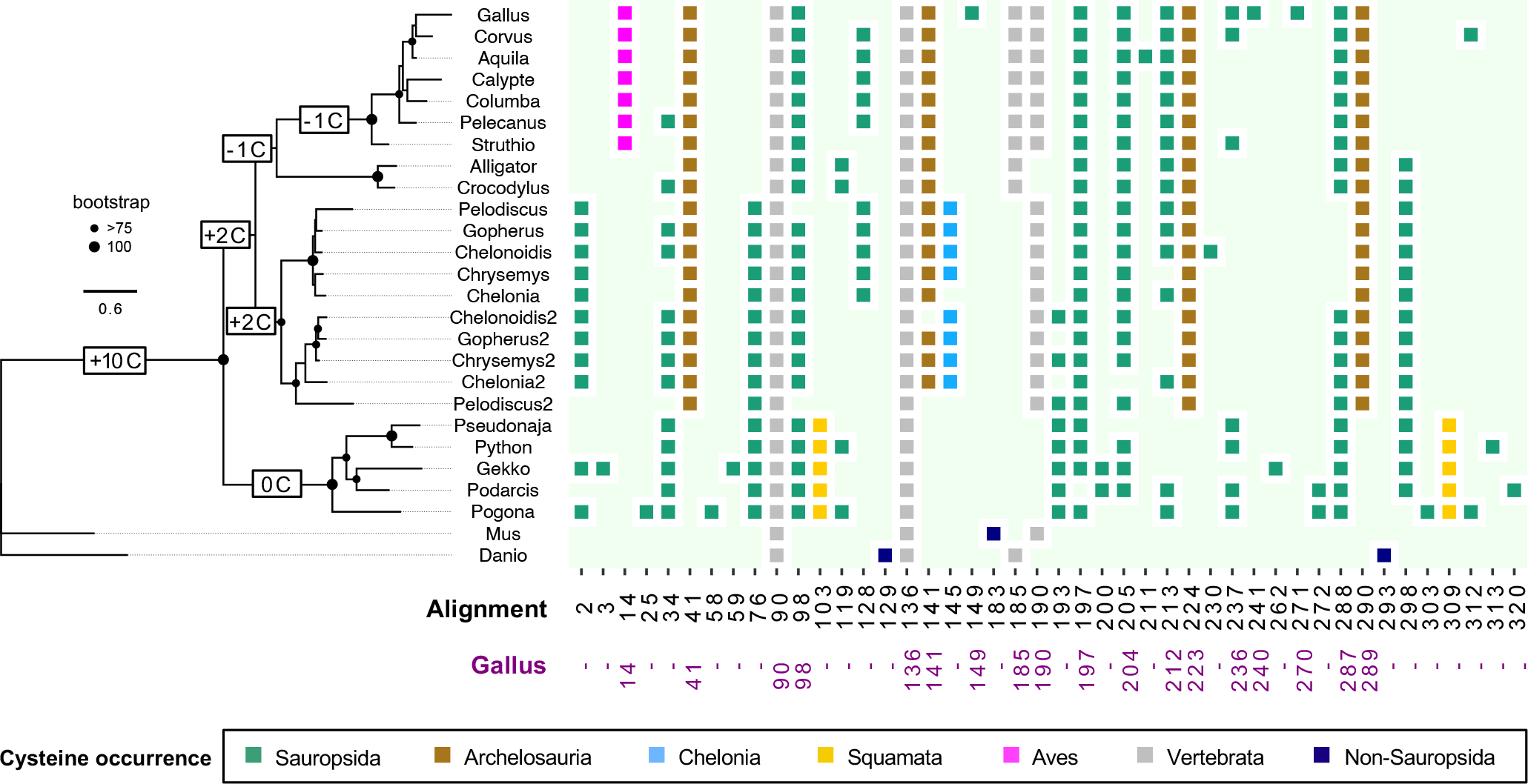
Cysteine enrichment in the Uox sequence at the stem of reptile evolution. Maximum likelihood phylogenetic tree of Uox sequences from vertebrate species and cysteine distribution in each sequence. *Danio rerio* Uox was used as an outgroup to root the tree. Bootstrap values (>75%; 100%) are indicated at internal nodes according to the legend; values below 75% are not indicated. Cysteines (C) enrichment (gain minus loss) in the Uox sequence during reptile evolution according to the bayes model of ancestral sequence reconstruction (see **S24 Fig**) is indicated on branches. Scale bar, substitution/site. Cysteine residues are numbered according to multiple alignment (see **S15 Fig**) and colored according to their presence in the different lineages as indicated in the legend.

Unlike cysteine, three amino acids emerged as highly variable among vertebrate orthologs: isoleucine, asparagine, and lysine. These residues are encoded by AU-rich codons and were found to be enriched in proteins from GC poor coding regions of vertebrate genomes (Huttener et al. 2019). Our findings that I, N, and K are enriched in several reptilian proteins compared to their mammalian and piscine orthologs (see **Figs 3C, 3D** and **S3**) is in agreement with the lower GC content measured in reptilian transcriptomes (Huttener et al. 2019).

Uricases are cofactor-independent oxidases (Fetzner and Steiner 2010; Bui and Steiner 2016) that catalyze the two-electron oxidation of UA to HIU that is typically irreversible. In this study we have shown that, unique amongst the uricases hitherto characterized, *Gg*Uox also reduces HIU back to UA. One appealing hypothesis for the key determinant of this unexpected feature is the rich complement of cysteine residues found in sauropsids’ uricases that might provide the source of reducing equivalents required for UA regeneration. A role for thiol groups in HIU reduction has been already proposed for the ‘conventional’ soybean Uox (Sarma and Tipton 2000). In the presence of DTT or free cysteine, it was found that O_2_ consumption greatly exceeded the concentration of UA in solution and that H_2_O_2_ production was attenuated. This is because DTT-mediated reduction of HIU resulted in UA regeneration, with the natural substrate becoming catalytic, rather than stoichiometric, in the reaction. With *Gg*Uox, the apparent rate-enhancement of UA conversion in the presence of HIUase can be interpreted in terms of the Le Chatelier’s principle with the latter enzyme able to degrade HIU once it leaves the active site. In the absence of HIUase, we propose that the many solvent-exposed cysteines act as reducing agents leading to UA regeneration thus resulting in its apparent slower rate of conversion under steady-state conditions. Our kinetic experiments with the competitive inhibitor AZA suggest, however, that HIU reduction also depends on active site residues. It is tempting to speculate that the nearby C287 as well as C14* and C289 play a role in this process possibly via solvent-mediated electron transfer as the X-ray structure of *Gg*Uox shows that compared to the ‘conventional’ *Af*Uox, its ‘peroxo hole’ (**S25 Fig**) is filled with a network of water molecules that might act as an electron conduit linking these cysteines with HIU. A water-mediated electron transfer involving sulfur containing amino acids has been proposed in model peptide systems (Wang et al. 2009). Whilst an adequate understanding of molecular mechanism underpinning the reversibility in *Gg*Uox catalysis will undoubtedly require additional studies, we propose that a key role for this is to be ascribed to the unique cysteine enrichment in reptilian uricase.

A single amino acid substitution (a tyrosine instead of an aliphatic amino acid) occurred at the reptile Uox active site. In the X-ray structure of *Gg*Uox we observed that this residue (Y230) lines the ‘peroxo hole’ and likely stabilizes PIU contributing to the equilibrium between this intermediate and HIU. To assess the role of this mutation, we compared the activity of wild-type *Gg*Uox with that of a mutated variant having a valine at the active site position. In view of a moderate decrease in the specific activity of the mutant protein, we conclude that this substitution is not the only determinant of the reduced catalytic efficiency of *Gg*Uox.

Gene co-option may occur through changes in expression pattern instead of or in addition to functional modifications of the encoded protein (True and Carroll 2002; Rebeiz et al. 2011; Malatesta et al. 2020). In the case of reptilian *Uox*, both mechanisms are involved. The shift in gene expression from the liver to the skin was mediated by the displacement of *Uox* 5’ exon that resulted in the loss of co-regulated transcription between Uox and the divergently oriented *Dnase2b* gene (see **fig 2B**). Separation of two head-to-head genes is far from common because head-to-head gene pairs were demonstrated to largely maintain this organization enabling coordinated expression and functionality during vertebrate evolution (Li et al. 2006).

The reptilian *Uox-Dnase2b* locus provides an example of a mechanism by which a functional split can occur in head-to-head genes without chromosomal rearrangement. Another modification of the *Uox* locus was observed in turtles with the presence of a duplicated *Uox* copy. The presence of multiple copies of *Uox* is exceptional as this gene is typically a single copy in all genomes. Since Uox is a tetrameric protein, multiple gene copies could result in a heterogeneous composition of the oligomer. This hypothesis is substantiated by the co-expression of the two *Uox* paralogs in turtles’ skin.

The expression of UOX detected in the feather epithelium of chick embryos is high at stages (around HH40; days 13-14) during which the most rapid growth of feather filaments occurs (Meyer and Baumgärtner 1998). At this time also the synthesis of keratins has started as the sheath covering the developing feather is fully keratinized after HH stage 41 (Lucas and Stettenheim 1972). Of note, apart from a protein with similarity to methanethiol oxidase (LOC425662) and two other exceptions, the list of the proteins that are co-expressed with *Gg*Uox in the skin of developing embryos (see **Fig 7D**) is entirely constituted by keratins, mostly of the β-type. Also in the adult, *Gg*Uox gene has been found in the same co-expression cluster with feather (β) keratins and keratin-associated genes (Bush et al. 2018). β-keratins are small proteins of the stratum corneum of birds and reptiles that have a high content of cysteines (4-20%), form stacked β- sheets, and aggregate into very resistant microfibrils. The dense β-keratin matrix forms hard skin appendages, like scales, scutes, turtles’ shell, claws, beaks, and feathers, providing waterproofing and mechanical protection to these structures (Holthaus et al. 2018). At variance with the more widespread α-keratins, the mechanism of aggregation of β-keratins is largely unknown (Gregg and Rogers 1986; Alibardi et al. 2006). Intriguingly, we observed a tendency to aggregate in *Gg*Uox which is dependent on the redox state and mediated by disulfide bond formation (see **Fig 6**).

Since the presence of *bona fide Uox* genes is in contrast with the uricotelism of birds and reptiles, an official gene symbol has not been established for their *Uox* orthologs, which are annotated in the sequence database with a generic ‘LOC’ prefix followed by a numeric Gene ID. To account for the catalytic activity of the encoded protein on uric acid, but the radically different physiological role, tissue specificity, and structural features, we propose to name the *Gallus gallus* gene ‘cysteine-rich urate oxidase’. A request for a new gene name and symbol (CRUOX) has been submitted to the chicken gene nomenclature committee.

## Material and Methods

### Amino acid content variation in vertebrate orthologous genes

A procedure for the analysis of the amino acid content of orthologous proteins from vertebrate species was implemented in R (https://github.com/lab83bio/AA_Comp). Orthogroups were downloaded from the OrthoDB database (v. 10) and parsed with R scripts to create a filtered dataframe with only orthogroups at the Vertebrata level, in single copy, and present in 90% of species. Then, a statistical analysis was performed to calculate *P*-value (t-test) and Log2 fold change from pairwise comparisons between the three different classes of Mammalia, Sauropsida and Actinopterygii. Volcano plots and bar plots were created using R packages “EnhancedVolcano’’ and “ggplot2”, respectively.

### Bioinformatics

Protein sequences were retrieved by homology search at NCBI, aligned using Clustal Omega at the EMBL-EBI webportal (https://www.ebi.ac.uk/Tools/msa/clustalo/) and visualized with ESPript (https://espript.ibcp.fr/ESPript/ESPript/) (Robert and Gouet 2014). The search of nitrogen metabolism gene sequences in complete genomes of vertebrate and invertebrate organisms was conducted with Hidden Markov models (Eddy 1996) for individual gene families. The figure of the gene distribution map was drawn using the R language and the taxizedb (https://docs.ropensci.org/taxizedb/) and ggtree (Yu 2020) libraries. Maximum-Likelihood trees were constructed with PhyML v3.0 (http://www.atgc-montpellier.fr/phyml/) (Guindon et al. 2010) using the automated model selection procedure (Lefort et al. 2017). Nodal support was estimated by bootstrap analysis with 100 replicates. Tree images were generated with FigTree v1.4.3 (https://github.com/rambaut/figtree/). Ancestral sequence reconstruction was performed using four different models implemented in Phangorn (Schliep 2011): maximum parsimony (mpr), mpr with accelerated transformation (mpr_acctran), maximum likelihood (ml), and highest posterior probability (bayes) based on the tree of **Fig 8**.

### Cloning, expression and purification of *Gg*Uox

For construction of *Gg*Uox expression plasmid, the uricase-like coding sequence (XM_015290876.2) of LOC101747367 was synthesized by Genscript (USA Inc.) into pcDNA3.1+/ C-(K)DYK standard vector. The sequence was PCR-amplified with specific primers (Fw: 5’-ATATGCTAGCATGAGCCAAGTGACAATTAAGG-3’ and Rev: 5’-ATATCTCGAGTTATTTTCCCTGGGCAGCTAC-3’) and subcloned into a pET28 expression vector at NheI/XhoI sites, in frame with the N-terminal His-tag and thrombin cleavage site. For construction of *Gg*UoxY230V and *Gg*UoxY230H expression plasmids, the mutated uricase-like coding sequence was synthesized by Genscript (USA Inc.) into a pET28 expression vector. All plasmids were verified by DNA sequencing and transformed into *E. coli* BL21-Codon Plus.

*Gg*Uox (wt and mutants) was expressed in auto-induction medium (LB supplemented with 0.05% glucose and 0.2% lactose) as follows: a 1 L culture was incubated for 10 hours at 30 °C until OD_600_ reached 0.6-0.8; the culture was subsequently incubated overnight at 20 °C. Cells were harvested by centrifugation (20 min, 5000 × g, 4 °C), and resuspended in 40 mL lysis buffer (50 mM Tris-HCl pH 7.6, 300 mM NaCl). After lysis by sonication on ice (15 min of 1 s pulse at 35% intensity / 1 s pause), the soluble fraction was recovered by centrifugation (30 min, 20000 × g, 4 °C). *Gg*Uox was purified using a 5 mL His-trap Ni-NTA column (GE Healthcare) equilibrated in 50 mM Tris-HCl pH 7.6, with 300 mM NaCl, and eluted with an imidazole gradient (10-500 mM) on Äkta FPLC (GE Healthcare). The purified protein was buffer exchanged with PD-10 columns (GE Healthcare) into 100 mM potassium phosphate buffer pH 7.6, containing 150 mM NaCl. The average yield was 60 mg of *Gg*Uox per liter of culture, with > 95% purity as estimated by resolution on 12% SDS-PAGE followed by staining with Coomassie blue. The final protein concentration was estimated by absorbance at 280 nm (ɛ_280_=35,505 M^-1^ cm^-1^; https://web.expasy.org/protparam/). Aliquots were snap-frozen in liquid nitrogen and stored at -80 °C.

*Gg*Uox purification through xanthine affinity chromatography was performed following the protocol previously used for *Dr*Uox purification (Marchetti et al. 2016). *Gg*Uox soluble fraction was applied to xanthine-agarose (X3128, Sigma-Aldrich) column equilibrated with 100 mM potassium phosphate pH 7.6. After washing, the protein was eluted in the same buffer containing 0.5 mM urate. An aliquot of xanthine-agarose resin was taken before elution and loaded on SDS-PAGE to confirm the specific binding of *Gg*Uox.

### Activity assays

Catalytic activity measurements were carried out with a Jasco V-750 spectrophotometer and a quartz cuvette of 1 cm path length. Uricase activity was assayed by monitoring the absorbance at 292 nm or at 250-340 nm in the presence of 100 μM urate in 100 mM potassium phosphate buffer, pH 7.6, at 25 °C. Concentration of urate (flash frozen stocks at a ∼5 mM concentration in aqueous solution with 0.02 N NaOH) was calculated by using the molar extinction coefficient ɛ_292_=12 650 M^-1^ cm^-1^. To avoid the interference of the Uox reaction product, 5-hydroxyisourate (HIU), which absorbs at 292 nm, *Danio rerio* (*Dr*) HIU hydrolase (HIUase or UraH; 0.25 μM) (Zanotti et al. 2006) or *Phaeodactylum tricornutum* allantoin synthase (Alls; 1.25 μM) (Oh et al. 2018) was added to the reaction mixture. HIUase converts HIU to 2-oxo-4-hydroxy-4-carboxy-5-ureidoimidazoline (OHCU), which has negligible absorbance at 292 nm; Alls converts HIU to allantoin, which does not absorb at 292 nm.

For the measurement of uricase activity in reducing conditions, *Gg*Uox was pre-incubated with 1 mM β-mercaptoethanol (βME). The uricase activity of *Dr*Uox (Marchetti et al. 2016) was measured in the same reaction conditions for comparison. The kinetic parameters of *Gg*Uox were measured at different concentrations (2-140 μM) of urate, in the presence of HIUase. Data of single wavelength (292 nm) kinetics were fitted either to the Michaelis-Menten or the Schell-Mendoza equation using the R packages “drc” and “lamW”, respectively.

The reverse reaction of HIU to urate was performed as follows: HIU was synthetized enzymatically from urate (500 μM) in the presence of an excess of *Dr*Uox (20 μM); the reaction was carried out in 100 mM potassium phosphate buffer, pH 7.6, at room temperature for 5 minutes, and monitored by UV spectra (250-350 nm). When the synthesis was completed, as determined by the shift of the urate peak from 292 to 301 nm, the reaction mixture was filtered through a centrifugal membrane filter (cut-off 10000 Da MW) to remove *Dr*Uox. *Gg*Uox (30 μM) pre-incubated with βME (300 μM) was added to HIU (60 μM) to start the reverse reaction. UV spectra were recorded from 250 to 350 nm. Spontaneous, non-enzymatic degradation of HIU was measured by adding either 100 mM potassium phosphate buffer or *Dr*Uox (30 μM) pre-incubated with βME (300 μM) to the filtered reaction mixture. Both the urate oxidation to HIU and the reverse reaction were assayed in the presence of the urate oxidase inhibitor 8-azaxanthine (AZA; 11460, Sigma-Aldrich), which was added to the reaction mixture at a final concentration of 50 μM.

### Stopped flow kinetics

Stopped flow experiments were carried out using an Applied Photophysics SX18 apparatus, with an instrumental dead time of 1.5 ms, equipped with a thermostatted bath set at 25 °C. *Gg*Uox or *Dr*Uox solutions at 70 μM concentration in 100 mM potassium phosphate buffer, pH 7.6, were mixed 1:1 with 50 μM urate in a single turnover condition and spectral changes were followed in the UV-visible range through a photodiode array. The time interval between spectra acquisitions was logarithmically distributed to have a higher acquisition rate (one spectrum every 1 ms) in the early phase, avoiding oversampling at longer time. To improve signal to noise ratio at least 3 kinetic traces were averaged.

Three-step (irreversible and reversible) models generation and data fitting to the models were carried out exploiting the Simbiology tool (MATLAB). All other data processing was carried out using MATLAB R2020b.

### Hydrogen peroxide quantification

Hydrogen peroxide produced in the urate oxidation reaction catalyzed by *Gg*Uox or *Dr*Uox was quantified by Ferrous Oxidation Xylenol Orange (FOX) assay (Wolff 1994). *Gg*Uox (30 μM) or *Dr*Uox (30 μM) was incubated with urate (25 μM) in 100 mM potassium phosphate buffer, pH 7.6, in the absence or presence of HIUase (0.25 μM). At time intervals (0, 2, 5, 10, 15, 30, and 60 min after urate oxidase addition) 20 μL-aliquots were taken and added to 200 μL FOX1 reagent (100 μM xylenol orange, 250 μM ammonium ferrous sulfate, 100 mM sorbitol, 25 mM H_2_SO_4_) in a 96-well plate and incubated for a minimum of 40 min at RT. The absorbance was read at 595 nm using an iMark microplate absorbance reader (Bio-rad). Each condition was conducted in triplicate. Concentrations of hydrogen peroxide were calculated from the absorbance at 595 nm using a regression equation and a correlation coefficient obtained experimentally from a standard curve. The standard curve (absorbance at 595 nm versus hydrogen peroxide concentration) was developed under the same conditions except that enzyme was not added and that dilutions of a hydrogen peroxide solution (8.8 M) were used as standards (final concentrations in the 200 μl assay: 10, 25, 50, 100 μM).

### X-ray crystallography

Although the cloning strategy employed allows for thrombin-mediated cleavage of the engineered N-terminal His affinity tag, all crystallographic work was carried out with an uncleaved version of the protein. Protein purification was performed using a combination of Ni-affinity and Superdex 75 size-exclusion chromatographic (SEC) steps. *Gg*Uox eluted from SEC in 50 mM Tris-HCl pH 8.0, 10 mM NaCl, 1 mM TCEP-HCl buffer was concentrated to 22 mg/mL and mixed with a 20-fold molar excess of the 8-azaxanthine (AZA) inhibitor. Sitting-drop vapor diffusion crystallization experiments were set up at 18 °C using a 1:1 protein:reservoir ratio dispensed with the Mosquito liquid handler (TTP Labtech). Single crystals of the *Gg*Uox-AZA complex were obtained after two-three days in the alternative orthorhombic space groups *C*222_1_ or *P*2_1_2_1_2_1_ using 8% PGA-LM, 0.3 M KBr, 0.1 M Tris-HCl pH 7.8 or 0.8 M K/Na tartrate tetrahydrate, 0.1 M HEPES pH 7.5 as crystallization reservoirs, respectively. For cryoprotection, crystals were transferred to their reservoir solution enriched with 20% ethylene glycol for a few minutes and then rapidly quenched and stored in liquid nitrogen. To probe oxidative conditions, crystals were transferred into cryoprotective solutions supplemented with 100 mM H_2_O_2_ for 15 minutes prior to cryocooling in liquid nitrogen. Complete datasets for reduced and oxidized *Gg*Uox-AZA complexes in both available space groups were measured at beamlines I24 and I04 of Diamond Light Source (Didcot, UK). Structure solution starting from the *Aspergillus flavus* Uox model (PDB code 7A0L) (McGregor et al. 2021) was accomplished using the molecular replacement method with the *Phaser* package (McCoy et al. 2007) as implemented in the CCP4 suite (Winn et al. 2011). The programs *COOT* (Emsley and Cowtan 2004) and *Refmac*5 (Steiner et al. 2003; Vagin et al. 2004; Murshudov et al. 2011) were used throughout for model rebuilding and refinement, respectively. Data collection and refinement statistics are available in **S2 Table**. Atomic coordinates and structure factors have been deposited with the Protein Data Bank with entry codes 8OFK, 8OH8, 8OIH, 8OIW.

### Dynamic light scattering

Dynamic light scattering (DLS) experiments were carried out with a Malvern Zetasizer NANO ZSP (Malvern Panalytical, Malvern, UK), at 25 °C, using a plastic cuvette. To measure the hydrodynamic diameter distribution of Uox treated with H_2_O_2_, a solution containing 7 μM *Gg*Uox or *Dr*Uox, pre-treated with 0 or 100 μM βME, was incubated with 100 μM H_2_O_2_ in 100 mM potassium phosphate buffer at pH 7.6. DLS spectra were recorded at different time points from 0 to 60 min. To measure the hydrodynamic diameter distribution of Uox during urate oxidation, a solution containing 21 μM *Gg*Uox or *Dr*Uox, pre-treated with 0 or 300 μM βME, was incubated with 2.1 mM urate in 100 mM potassium phosphate buffer at pH 7.6. DLS spectra were recorded at different time points (0-60 min) after diluting the reaction to a final protein concentration of 7 μM. The data were analyzed with the Malvern Zetasizer software v8.1 using standard settings.

### Atomic force microscopy

Atomic force microscopy (AFM) images (512x512 pixels with a scan size of 2 µm) were collected in air with a Nanoscope IIIA microscope (Digital Instruments, Santa Barbara, CA, USA) operating in tapping mode and equipped with the E scanner. Commercial silicon cantilevers (MikroMasch, Tallinn, Estonia) with a nominal tip radius of 5 nm were used. In a first set of experiments, 21 μM *Gg*Uox or *Dr*Uox pre-incubated with 0 or 300 μM βME was mixed with 2.1 mM urate in 100 mM potassium phosphate buffer, pH 7.6, and incubated at RT for 60 min. In a second set, 7 μM *Gg*Uox or *Dr*Uox pre-incubated with 0 or 100 μM βME was mixed with 100 μM H_2_O_2_ in 100 mM potassium phosphate buffer, pH 7.6, and incubated at RT for 60 min. All samples were diluted in deposition buffer (4 mM HEPES pH 7.4, 10 mM NaCl, 2 mM MgCl_2_) to a final protein concentration of 100 nM, and 20 µl were deposited onto freshly cleaved mica that had been previously incubated with 50 µL MilliQ water for 5 min, and with 50 µl deposition buffer for 3 min. After 5 min incubation, the excess of sample on the mica surface was removed by rinsing with 50 µl MilliQ water and drying under a nitrogen flux.

### RNA-seq analysis

Paired-end RNA-seq reads of liver or skin samples from different reptiles were retrieved by keyword searches and downloaded from the European Nucleotide Archive (ENA) database (Tables S3 and S4). Gene expression was quantified as transcripts per million (TPM) with Kallisto (v.0.44.0) (Bray et al. 2016) using indexes built on the transcriptomes of *Gallus gallus* (GCF_016699485.2), *Anolis carolinensis* (GCF_000090745.1), *Serinus canaria* (GCF_022539315.1), *Pelodiscus sinensis* (GCF_000230535.1), and *Chelonia mydas* (GCF_015237465.2). Principal component analysis was conducted with the variance stabilizing transformation (vst) and PCAplot functions or the DESeq2 R package (Love et al. 2014). Plots for expression levels of individual genes were obtained with IsoformSwitchAnalyzeR (Vitting-Seerup and Sandelin 2019). Cluster analysis of gene expression profiles was conducted with the hclust and cutree (k=100) functions of the R package.

### Embryo collection and in situ hybridization

Fertile chicken eggs (HyLine, Iowa; not a commercially available source) were incubated in a humidified incubator at 37.5 °C for 0.5 to 5 days. Embryos were collected into chilled chick saline (123 mM NaCl), removed from the vitelline membrane and cleaned of yolk.

Extra-embryonic membranes and large body cavities (brain vesicles, atria, allantois, eye) were opened to minimize trapping of the in situ reagents. Embryos were fixed overnight at 4 °C in freshly prepared 4% paraformaldehyde in PBS, washed twice briefly in in PBS plus 0.1% Triton X-100 then dehydrated through a graded MEOH series and stored at -20 °C overnight in 100% MEOH. cDNA templates for generating all antisense RNA probes were obtained by reverse transcriptase-polymerase chain reaction using pooled RNA from embryos between HH stages 4 and 30. Primer sequences were designed using the mRNA sequence in the NCBI database.

The sequence 5′-AATTAACCCTCACTAAAGG-3, corresponding to the T3 DNA polymerase binding site, was added to the 5′ end of each reverse primer. The following primers were used: LOC101747367 (1089t): Forward, 5’-ATAAACCTCTTGGCTTCCTG-3’, Reverse, 5’-AATTAACCCTCACTAAAGGATATGGTGCAGGACGTTATC-3’. Embryo processing, antisense RNA probe preparation and whole-mount ISHs were performed as described (Antin et al. 2010). A detailed protocol is available for download at http://geisha.arizona.edu.

## Acknowledgments

We thank Davide Cavazzini and Marialaura Marchetti for assistance, Francesco Trigiante and Luciano Macaluso for their help in the initial phase of the work, and Lorenzo Alibardi for valuable discussions. Scientists and staff at Diamond Light Source (Didcot, UK) are gratefully acknowledged for their support. This work benefited from the equipment and framework of the COMP-HUB and COMP-R initiatives, funded by the “Departments of Excellence” program of the Italian Ministry for University and Research (MIUR, 2018-2022 and MUR, 2023-2027), and from the High Performance Computing facility of the University of Parma, Italy. G.M. is a recipient of postdoctoral fellowships from COMP-HUB. This work was supported by the Italian Ministry for Education, University and Research PRIN grant 2017483NH8 to R.P. and by the United Kingdom Biotechnology and Biological Sciences Research Council (BBSRC) grant BB/P000169/1 awarded to R.A.S.

## Data availability

The datasets and computer code used in this study are available in GitHub at the address: https://github.com/lab83bio/AA_Comp.

## Supporting information

**S1 Fig. Evidence for a complete Uox coding sequence in *Anolis carolinensis*.** (A) Alignment of the translated CDS with database Uox sequences of *Pogona vitticeps* (XP_020650701.1), *Gekko japonicus* (XP_015274881.1) and the N-truncated *A. carolinensis* sequence (XP_016850379.1). The amino acid sequence corresponding to the first exon of *G. japonicus Uox* (**Fig 2A**) is indicated by a green bar. (B) Consensus mRNA obtained from the multiple alignment of *A. carolinensis* sequences identified through homology search (tblast) in dbEST. The putative CDS is in bold.

**S2 Fig. Variation in amino acid composition of vertebrate orthologous proteins.** (A) Volcano plot depicting the cysteine content variation of orthologous proteins between groups of vertebrates: mammals vs. fishes. Horizontal dashed line indicates p-value of 1e^-16^; vertical dashed lines indicate log_2_ fold change of −1 and 1. (B) Bar plot illustrating the number of orthogroups in which the content of the corresponding amino acid is significantly increased (light orange) or decreased (green) according to p-value < 1e^-16^ and log_2_ fold change > ±1 in mammals vs. fishes.

**S3 Fig. Variation in I, N, K amino acid composition of vertebrate orthologous proteins.** (A) Volcano plot depicting the “INK” content variation of orthologous proteins between groups of vertebrates: sauropsids vs. mammals (upper panels), sauropsids vs. fishes (middle panels), and mammals vs. fishes (lower panels). Horizontal dashed line indicates p-value of 1e^-16^; vertical dashed lines indicate log_2_ fold change of −1 and 1.

**S4 Fig. Purification and characterization of recombinant *Gg*Uox.** (A) SDS-PAGE of the purification of *Gg*Uox by Ni^2+^-affinity chromatography. M: marker; I: post-induction total cell fraction; S: soluble cell fraction; FT: flow-through; E1-E3: elution fractions. (B) Size-exclusion chromatography (Superose 6 Increase 10/300 column) profile of purified *Gg*Uox (light orange) and of protein markers (light blue). (C) Measurement of hydrogen peroxide generated in the urate oxidation reaction catalysed by *Gg*Uox or *Dr*Uox. Reactions were carried out under single turnover conditions with 100 mM potassium phosphate (pH 7.6), 30 μM enzyme, 25 μM urate. The amount of hydrogen peroxide was quantified by Ferrous Oxidation Xylenol Orange (FOX) assay.

**S5 Fig. Time-resolved UV-Vis spectra showing urate oxidation by *Dr*Uox.** (A-B) Reaction contained 100 mM potassium phosphate (pH 7.6), 100 μM urate, 30 μM *Dr*Uox, and (A) 0 or (B) 300 μM β-mercaptoethanol (βME). Spectra were acquired every 1 min at 25 °C.

**S6 Fig. *Gg*Uox-catalyzed or spontaneous HIU degradation.** (A-C) Time-resolved UV-Vis spectra showing HIU degradation. Reaction contained 100 mM potassium phosphate (pH 7.6), 60 μM HIU, and (A) 30 μM *Dr*Uox, 300 μM β-mercaptoethanol (βME), or (C) *Gg*Uox (30 μM), 300 μM βME, and 8-azaxanthine (AZA; 50 μM). Spectra were acquired at 25 °C. (D) Time-dependent change of absorbance at 315 nm during HIU degradation by *Gg*Uox (pink), *Gg*Uox with AZA (bordeaux), and *Dr*Uox (light blue).

**S7 Fig. Time-resolved UV-Vis spectra of urate oxidation by *Gg*Uox in the presence of 8-azaxanthine (AZA).** Reaction contained 100 mM potassium phosphate (pH 7.6), 100 μM urate, 10 μM *Gg*Uox, 50 μM AZA and 1 μM Alls. Spectra were acquired every 1 min at 25 °C.

**S8 Fig. Stopped flow analysis of *Gg*Uox and *Dr*Uox reactions.** Stopped flow kinetics at selected wavelengths of *Gg*Uox (left) or *Dr*Uox (right). Reactions were carried out under single turnover conditions at 25 °C with 100 mM potassium phosphate (pH 7.6), 35 μM enzyme, 25 μM UA. To improve signal to noise ratio 3 mixing kinetics were averaged. (B) Main spectral components obtained from the SVD analysis on the two time-resolved spectral series following the reaction between UA and *Gg*Uox (light blue: SVD component I; yellow: SVD component II) or *Dr*Uox (blue: SVD component I; orange: SVD component II) . Spectral components are reported as the product of the spectral component (u) multiplied by its corresponding relative weight (s). (C) Difference spectrum (green closed circles) between 5-peroxyisourate (PIU; the second intermediate of Uox reaction) and HIU (experimentally determined spectra were extracted from Kahn and Tipton 1998), and SVD component I spectrum (blue line) from *Dr*Uox SVD analysis. Data are normalized for UA concentration.

**S9 Fig. Kinetics parameters determination of *Gg*Uox and *Dr*Uox activity upon model fitting.** (A) Three-steps irreversible model for UA to HIU conversion. Intermediate species were already attributed to urate dianion (UA^2-^) and 5-peroxyisourate (PIU) (Kahn and Tipton 1998) (Kahn and Tipton 1998). (B) Kinetic constants obtained from stopped flow data fitting to SimBiology models. *Gg*Uox data were fitted to the reversible three step model shown in **Fig 4J**; *Dr*Uox data were fitted to the irreversible three step model shown in (A).

**S10 Fig. *Gg*Uox active sites in the tetramer.** Stick representation of the active sites (A) between chain B and chain A, (B) between chain C and chain D, and (C) between chain D and chain C with the four protomers highlighted by different colors. 2*mF*_o_-*DF*_c_ electron density map for AZA, solvent molecules in its proximity, and C287 is shown in blue at the +1.0s level. In (C) the electron density for the solvent above AZA is also shown as chicken-wire representation at the +0.5s level in grey. Selected hydrogen bonds are highlighted in black as broken lines. In general, the solvent structure is quite conserved and best explained by a mixture of dioxygen and water molecules. Water molecules w1, w2, w4, w5 above AZA appear well defined whilst water molecule w3 is rather mobile and in one active site (B) is not visible. In another active site (C) w3 occupies two alternative positions with dioxygen and a solvent molecule (W) sharing the same position with partial occupancy.

**S11 Fig. DLS analysis of the size of *Gg*Uox in solution.** (A) Samples containing 100 mM potassium phosphate (pH 7.6) and 20 μM *Gg*Uox were incubated at RT and analysed at different time points after the addition of 100 μM urate (UA). (B) Samples containing 100 mM potassium phosphate (pH 7.6) and 20 μM *Gg*Uox were incubated at RT and analysed at different time points after the addition of 100 μM urate (UA). Samples containing 100 mM potassium phosphate (pH 7.6) and 7 μM *Gg*Uox were incubated at RT and analysed at different time points after the addition of 100 μM H_2_O_2_. After 16 min, 5 mM β-mercaptoethanol was added; samples measured at 20, 30, and 60 minutes, containing β-mercaptoethanol are indicated with *.

**S12 Fig. AFM analysis of the size of *Gg*Uox and *Dr*Uox in solution.** Volume distribution of (A) native *Gg*Uox, (B) native *Dr*Uox, (C) *Gg*Uox pre-incubated with β-mercaptoethanol (βME), and (D) *Dr*Uox pre-incubated βME. (E) Calibration curve used to infer the molecular weights of *Gg*Uox and *Dr*Uox. Data points correspond to: Carbonic anhydrase (30 kDa), Conalbumin (75 kDa), Alcohol dehydrogenase (140 kDa) and Catalase (240 kDa). The red square indicates the molecular weight of *Gg*Uox (156 kDa) based on the highest peak of the volume distribution in (A); the blue square indicates the molecular weight of *Dr*Uox (146 kDa) based on the highest peak of the volume distribution in (B). (F-G) Volume distribution of *Gg*Uox after incubation at RT with (F) urate (UA) or (G) hydrogen peroxide (H_2_O_2_). The highest peak in (F) corresponds to 10000 nm^3^; the highest peak in (G) corresponds to 2000 nm^3^.S13. Oxidized cysteines in the *Gg*Uox X-ray structure. Example of cysteine oxidation to its sulfinic form (CSD) following H_2_O_2_ treatment. 2*mF*_o_-*DF*_c_ electron density map is shown in blue at the +1.0s level.

**S13 Fig. Oxidized cysteines in the *Gg*Uox X-ray structure.** Example of cysteine oxidation to its sulfinic form (CSD) following H_2_O_2_ treatment. 2*mF*_o_-*DF*_c_ electron density map is shown in blue at the +1.0s level.

**S14 Fig. *Uox* gene duplication in chelonian reptiles.** (A) Chronogram of vertebrate phylogeny derived from TimeTree. Conserved synteny between *Uox* and *Dnase2b* genes in the different species and tandem duplication of *Uox* in chelonian reptiles are displayed at the terminal nodes. The 5’exon of reptilian *Uox* is represented by a green square before the 3’end of *Dnase2b* gene. The letter above the arrow corresponds to the amino acid at position 230 in the sequence of *Gg*Uox. (B) Maximum likelihood unrooted phylogenetic tree of Uox sequences from animal species. Genus names and silhouettes of one or more taxa for each taxonomic group (legend) are shown. Paralogous proteins in chelonian reptiles (Uox2) are highlighted in dark green. Scale bar, substitution/site. (C) Specific activity of *Gg*Uox wild-type (wt) and mutants Y230H, Y230V. Reactions were carried out in 100 mM potassium phosphate (pH 7.6), at 25 °C, with the addition of 0.25 µM HIUase. Error bars represent the standard deviation of three measurements.

**S15 Fig. Multiple alignment of vertebrate Uox proteins that was used to infer the phylogenetic tree presented in** Fig 8. Shown is the portion of the alignment extending from the first to the last amino acid of *Gallus gallus* (*Gg*) Uox sequence. Residue numbering at the top refers to *Gg*Uox; numbering at the bottom refers to the alignment. Cysteine residues are marked by triangles coloured as indicated in the legend, and are the same shown in Fig 8.

**S16 Fig. Recombinant expression of *Gg*Uox mutants.** SDS-PAGE of the purification of *Gg*Uox mutants (A) Y230H and (B) Y230V by Ni^2+^-affinity chromatography. M: marker; S: soluble cell fraction; P: insoluble cell fraction (pellet); FT: flow-through; E1-E2: elution fractions.

**S17 Fig. Principal component analysis (PCA) of the RNA-seq datasets used for the quantification of *Uox* expression.** Data belong to different tissues of *Gallus gallus* (upper panel), *Serinus canaria* (middle panel), and *Anolis carolinensis* (lower panel). PCA analysis was conducted with the variance stabilizing transformation (vst) and PCAplot functions of the DESeq2 R package based on transcript abundance value determined by the Kallisto software. Sequence Read Archive (SRA) accessions of the shown datasets are reported in **S3 Table**.

**S18 Fig. Expression of reptilian *Dnase2b* and *UraH*.** Expression levels (TPM: Transcripts Per kilobase Million) of *Dnase2b* (upper panels) and *Urah* (lower panels) in the liver and tegumental tissues (skin and feather follicle) of *Gallus gallus, Serinus canaria,* and *Anolis carolinensis* as derived from RNA-seq data analysis.

**S19 Fig. Expression of the two *Uox* paralogs, *Dnase2b*, and *Urah* in chelonians.** Expression levels (TPM: Transcripts Per kilobase Million) in the kidney and skin of *Chelonia mydas* and in the liver of *Pelodiscus sinensis*, of *Uox* (uricase.1 and uricase.2), *Dnase2b*, and *Urah*, as derived from RNA-seq data analysis.

**S20 Fig. Expression profiles of *Gg*Uox co-expressed genes in skin embryo development.** Gene co-expression cluster was obtained by hierarchical clustering with the hclust function of the R package based on euclidean distances of scaled transcript abundance values. Transcript abundance was determined with Kallisto based on RNA-seq data of the PRJNA397795 Bioproject (**S3 Table**). Gene accession numbers and descriptions for individual traces (gray lines) are reported in **Fig 7D**. The median value of the cluster is indicated by a brown line.

**S21 Fig. UOX transcription start site (TSS) peaks and expression levels during embryo development.** A strong single peak (arrow) is mapped to the 5’ end of *G. gallus* UOX (LOC101747367) through Cap Analysis of Gene Expression (CAGE) [PMC5600399] visualized by Chicken-ZENBU. Developmental stage-resolved analysis (bar graph) shows late stage-specific expression of UOX (Hamburger and Hamilton stage 41 [HH41], day 15).

**S22 Fig. Expression of *Gallus gallus* URAH and DNASE2B during embryogenesis.** In situ hybridization analysis of URAH (upper panels) and DNASE2B (lower panels) expression in chick embryos at HH developmental stages 36 (left panels) and 39 (bottom panels).

**S23 Fig. Uricase activity of *Gg*Uox at acidic pH.** Reactions were carried out in 100 mM potassium phosphate, pH 6.5 or 5.5, at 25 °C, with the addition of 1 µM HIUase. The activities are expressed in %, where 100% refers to the activity of *Gg*Uox measured at pH 7.6. Error bars represent the standard deviation of three measurements.

**S24 Fig. Cysteine enrichment in reptilian uricase.** (A) Net cysteine variation (gain minus loss) along selected nodes of the vertebrate uricase tree according to four different models implemented in Phangorn: maximum parsimony (mpr), mpr with accelerated transformation (mpr_acctran), maximum likelihood (ml), and highest posterior probability (bayes). (B) Absolute (top) and relative (bottom) amino acid enrichment in the uricase sequence of the stem reptile. Data are mean and standard deviation of the four different models in (A).

**S25 Fig. *Gg*Uox and *Aspergillus flavus* (*Af*) Uox active sites.** Superposition of the active sites of *Gg*Uox (wheat) and *Af*Uox (light green) in the AZA (grey and black carbon atoms for *Af*UOX and *Gg*Uox, respectively) bound state. Residues in the different orthologs are highlighted using the same color scheme. Solvent molecules in *Gg*Uox (w_1_-w_5_, w_N_, and O_2_) and *Af*Uox (W and w_N_) are shown in red and light green, respectively. Asterisks indicate residues belonging to different protomers. Although several residues are conserved between *Gg*Uox and *Af*Uox including the AZA-bound w_N_, amino acid differences are observed at important topological positions. The V227→Y230 replacement forces an alternative conformation for *Gg*Uox N257 side chain that is H-bonded to w_1_. In *Af*Uox this residue is involved in the coordination of the only water molecule (W) that occupies the ‘peroxo hole’ and is also stabilised by the side chain of T57*. Additionally, H256→F259 and I288→C287 replacements increase the volume of the ‘peroxo hole’ that accommodates under normoxic conditions a more complex solvent structure constituted by a mixture of O_2_ and water molecules. Coordinates for *Af*Uox were taken from the main conformation of the high-resolution crystal structure jointly refined using neutron and X-ray data (PDB code 7A0L) **(Bui et al. 2014; McGregor et al. 2021)**.

S1 Table. Orthogroups in which the cysteine content is significantly different between groups of vertebrates: Sauropsida (S), Mammalia (M), and Actinopterygii (A).

**S2 Table. Data collection and refinement statistics of X-ray crystallography.**

**S3 Table. Sequence read archive IDs of the RNA-seq data analyzed for gene expression in *G. gallus, S. canaria, A. carolinensis*.**

S4 Table. Sequence read archive IDs of the RNA-seq data analyzed for gene expression in Chelonians.

## Notes

https://github.com/lab83bio/AA_Comp

### Competing Interest Statement

The authors have declared no competing interest.

## References

Alibardi L, Dalla Valle L, Toffolo V, Toni M. 2006. Scale keratin in lizard epidermis reveals amino acid regions homologous with avian and mammalian epidermal proteins. Anat. Rec. A. Discov. Mol. Cell. Evol. Biol. 288A:734–752.

Ames BN, Cathcart R, Schwiers E, Hochstein P. 1981. Uric acid provides an antioxidant defense in humans against oxidant- and radical-caused aging and cancer: a hypothesis. Proc. Natl. Acad. Sci. 78:6858–6862.

Antin PB, Pier M, Sesepasara T, Yatskievych TA, Darnell DK. 2010. Embryonic expression of the chicken Krüppel-like (KLF) transcription factor gene family. Dev. Dyn. Off. Publ. Am. Assoc. Anat. 239:1879–1887.

Bartlett GJ, Borkakoti N, Thornton JM. 2003. Catalysing new reactions during evolution: economy of residues and mechanism. J. Mol. Biol. 331:829–860.

Bray NL, Pimentel H, Melsted P, Pachter L. 2016. Near-optimal probabilistic RNA-seq quantification. Nat. Biotechnol. 34:525–527.

Bui S, Steiner RA. 2016. New insight into cofactor-free oxygenation from combined experimental and computational approaches. Curr. Opin. Struct. Biol. 41:109–118.

Bui S, von Stetten D, Jambrina PG, Prangé T, Colloc’h N, de Sanctis D, Royant A, Rosta E, Steiner RA. 2014. Direct Evidence for a Peroxide Intermediate and a Reactive Enzyme-Substrate-Dioxygen Configuration in a Cofactor-free Oxidase. Angew. Chem. Int. Ed. 53:13710–13714.

Bush SJ, Freem L, MacCallum AJ, O’Dell J, Wu C, Afrasiabi C, Psifidi A, Stevens MP, Smith J, Summers KM, et al. 2018. Combination of novel and public RNA-seq datasets to generate an mRNA expression atlas for the domestic chicken. BMC Genomics 19:594.

Campbell JW, Vorhaben JE, Smith DD. 1987. Uricoteley: Its nature and origin during the evolution of tetrapod vertebrates. J. Exp. Zool. 243:349–363.

Chiu Y-C, Hsu T-S, Huang C-Y, Hsu C-H. 2021. Structural and biochemical insights into a hyperthermostable urate oxidase from Thermobispora bispora for hyperuricemia and gout therapy. Int. J. Biol. Macromol. 188:914–923.

Colloc’h N, El Hajji M, Bachet B, L’Hermite G, Schiltz M, Prangé T, Castro B, Mornon J-P. 1997. Crystal Structure of the protein drug urate oxidase-inhibitor complex at 2.05 Å resolution. Nat. Struct. Biol. 4:947–952.

Dembech E, Malatesta M, De Rito C, Mori G, Cavazzini D, Secchi A, Morandin F, Percudani R. 2023. Identification of hidden associations among eukaryotic genes through statistical analysis of coevolutionary transitions. Proc. Natl. Acad. Sci. 120:e2218329120.

Dessauer HC. 1970. Blood chemistry of reptiles: physiological and evolutionary aspects. In: The Biology of the Reptilia. Vol. 3. Academic Press, NY, USA. p. 1–72.

Eddy SR. 1996. Hidden Markov models. Curr. Opin. Struct. Biol. 6:361–365.

Emsley P, Cowtan K. 2004. Coot: model-building tools for molecular graphics. Acta Crystallogr. D Biol. Crystallogr. 60:2126–2132.

Fetzner S, Steiner RA. 2010. Cofactor-independent oxidases and oxygenases. Appl. Microbiol. Biotechnol. 86:791–804.

Gregg K, Rogers GE. 1986. Feather Keratin: Composition, Structure and Biogenesis. In: Bereiter-Hahn J, Matoltsy AG, Richards KS, editors. Biology of the Integument: 2 Vertebrates. Berlin, Heidelberg: Springer. p. 666–694. Available from: https://doi.org/10.1007/978-3-662-00989-5_33

Guindon S, Dufayard J-F, Lefort V, Anisimova M, Hordijk W, Gascuel O. 2010. New algorithms and methods to estimate maximum-likelihood phylogenies: assessing the performance of PhyML 3.0. Syst. Biol. 59:307–321.

Hamburger V, Hamilton HL. 1992. A series of normal stages in the development of the chick embryo. Dev. Dyn. 195:231–272.

Hayashi S, Fujiwara S, Noguchi T. 2000. Evolution of urate-degrading enzymes in animal peroxisomes. Cell Biochem. Biophys. 32 Spring:123–129.

Heitaroh I, Itaru Y, Eiichi G, Kyoji M, Mitsutaka N, Keiko S. 1973. Potent competitive uricase inhibitors—2,8-diazahypoxanthine and related compounds. Biochem. Pharmacol. 22:2237–2245.

Hibi T, Kume A, Kawamura A, Itoh T, Fukada H, Nishiya Y. 2016. Hyperstabilization of Tetrameric Bacillus sp. TB-90 Urate Oxidase by Introducing Disulfide Bonds through Structural Plasticity. Biochemistry 55:724–732.

Holthaus KB, Eckhart L, Dalla Valle L, Alibardi L. 2018. Review: Evolution and diversification of corneous beta-proteins, the characteristic epidermal proteins of reptiles and birds. J. Exp. Zoolog. B Mol. Dev. Evol. 330:438–453.

Huttener R, Thorrez L, in’t Veld T, Granvik M, Snoeck L, Van Lommel L, Schuit F. 2019. GC content of vertebrate exome landscapes reveal areas of accelerated protein evolution. BMC Evol. Biol. 19:144.

Kahn K, Tipton PA. 1998. Spectroscopic Characterization of Intermediates in the Urate Oxidase Reaction. Biochemistry 37:11651–11659.

Keebaugh AC, Thomas JW. 2010. The Evolutionary Fate of the Genes Encoding the Purine Catabolic Enzymes in Hominoids, Birds, and Reptiles. Mol. Biol. Evol. 27:1359–1369.

Kratzer JT, Lanaspa MA, Murphy MN, Cicerchi C, Graves CL, Tipton PA, Ortlund EA, Johnson RJ, Gaucher EA. 2014. Evolutionary history and metabolic insights of ancient mammalian uricases. Proc. Natl. Acad. Sci. 111:3763–3768.

Krissinel E, Henrick K. 2004. Secondary-structure matching (SSM), a new tool for fast protein structure alignment in three dimensions. Acta Crystallogr. D Biol. Crystallogr. 60:2256–2268.

Lefort V, Longueville J-E, Gascuel O. 2017. SMS: Smart Model Selection in PhyML. Mol. Biol.Evol. 34:2422–2424.

Li Y-Y, Yu H, Guo Z-M, Guo T-Q, Tu K, Li Y-X. 2006. Systematic Analysis of Head-to-Head Gene Organization: Evolutionary Conservation and Potential Biological Relevance. PLoS Comput. Biol. 2:e74.

Li Z, Hoshino Y, Tran L, Gaucher EA. 2022. Phylogenetic Articulation of Uric Acid Evolution in Mammals and How It Informs a Therapeutic Uricase.Battistuzzi FU, editor. Mol. Biol. Evol. 39:msab312.

Lizio M, Deviatiiarov R, Nagai H, Galan L, Arner E, Itoh M, Lassmann T, Kasukawa T, Hasegawa A, Ros MA, et al. 2017. Systematic analysis of transcription start sites in avian development. PLOS Biol. 15:e2002887.

Love MI, Huber W, Anders S. 2014. Moderated estimation of fold change and dispersion for RNA-seq data with DESeq2. Genome Biol. 15:550.

Lucas AM, Stettenheim PR. 1972. Avian Anatomy Integument. Avian Anatomy Project, Poultry Research Branch, Animal Science Research Division, Agricultural Research Service, U.S. Department of Agriculture

Malatesta M, Mori G, Acquotti D, Campanini B, Peracchi A, Antin PB, Percudani R. 2020. Birth of a pathway for sulfur metabolism in early amniote evolution. *Nat*. Ecol. Evol. 4:1239–1246.

Marchetti M, Liuzzi A, Fermi B, Corsini R, Folli C, Speranzini V, Gandolfi F, Bettati S, Ronda L, Cendron L, et al. 2016. Catalysis and Structure of Zebrafish Urate Oxidase Provide Insights into the Origin of Hyperuricemia in Hominoids. Sci. Rep. 6:38302.

McCoy AJ, Grosse-Kunstleve RW, Adams PD, Winn MD, Storoni LC, Read RJ. 2007. Phaser crystallographic software. J. Appl. Crystallogr. 40:658–674.

McGregor L, Foldes T, Bui S, Moulin M, Coquelle N, Blakeley MP, Rosta E, Steiner RA. 2021. Joint neutron/X-ray crystal structure of a mechanistically relevant complex of perdeuterated urate oxidase and simulations provide insight into the hydration step of catalysis. IUCrJ 8:46–59.

McLennan DA. 2008. The Concept of Co-option: Why Evolution Often Looks Miraculous. Evol.Educ. Outreach 1:247–258.

Meyer W, Baumgärtner G. 1998. Embryonal feather growth in the chicken. J. Anat. 193:611–616.

Mori G, Delfino D, Pibiri P, Rivetti C, Percudani R. 2022. Origin and significance of the human DNase repertoire. Sci. Rep. 12:10364.

Murshudov GN, Skubák P, Lebedev AA, Pannu NS, Steiner RA, Nicholls RA, Winn MD, Long F, Vagin AA. 2011. REFMAC5 for the refinement of macromolecular crystal structures. Acta Crystallogr. D Biol. Crystallogr. 67:355–367.

Nishikimi M, Fukuyama R, Minoshima S, Shimizu N, Yagi K. 1994. Cloning and chromosomal mapping of the human nonfunctional gene for L-gulono-gamma-lactone oxidase, the enzyme for L-ascorbic acid biosynthesis missing in man. J. Biol. Chem. 269:13685–13688.

Nishimura H, Yoshida K, Yokota Y, Matsushima A, Inada Y. 1982. Physicochemical Properties and States of Sulfhydryl Groups of Uricase from Candida utilis. J. Biochem. (Tokyo) 91:41–48.

Noguchi T, Takada Y, Fujiwara S. 1979. Degradation of uric acid to urea and glyoxylate in peroxisomes. J. Biol. Chem. 254:5272–5275.

Oda M, Satta Y, Takenaka O, Takahata N. 2002. Loss of Urate Oxidase Activity in Hominoids and its Evolutionary Implications. Mol. Biol. Evol. 19:640–653.

Oh J, Liuzzi A, Ronda L, Marchetti M, Corsini R, Folli C, Bettati S, Rhee S, Percudani R. 2018.Diatom Allantoin Synthase Provides Structural Insights into Natural Fusion Protein Therapeutics. ACS Chem. Biol. 13:2237–2246.

Peterson DW, Hamilton WH, Lilyblade AL. 1971. Hereditary Susceptibility to Dietary Induction of Gout in Selected Lines of Chickens. J. Nutr. 101:347–354.

Ramazzina I, Folli C, Secchi A, Berni R, Percudani R. 2006. Completing the uric acid degradation pathway through phylogenetic comparison of whole genomes. Nat. Chem. Biol. 2:144–148.

Rebeiz M, Jikomes N, Kassner VA, Carroll SB. 2011. Evolutionary origin of a novel gene expression pattern through co-option of the latent activities of existing regulatory sequences. Proc. Natl. Acad. Sci. 108:10036–10043.

Robert X, Gouet P. 2014. Deciphering key features in protein structures with the new ENDscript server. Nucleic Acids Res. 42:W320–4.

Rothschild BM, Schultze H-P, Pellegrini R. 2013. Osseous and Other Hard Tissue Pathologies in Turtles and Abnormalities of Mineral Deposition. In: Brinkman DB, Holroyd PA, Gardner JD, editors. Morphology and Evolution of Turtles. Vertebrate Paleobiology and Paleoanthropology. Dordrecht: Springer Netherlands. p. 501–534. Available from: http://link.springer.com/10.1007/978-94-007-4309-0_27

Rothschild BM, Tanke D, Carpenter K. 1997. Tyrannosaurs suffered from gout. Nature 387:357–357.

Salway JG. 2018. The Krebs Uric Acid Cycle: A Forgotten Krebs Cycle. Trends Biochem. Sci. 43:847–849.

Sarma AD, Tipton PA. 2000. Evidence for Urate Hydroperoxide as an Intermediate in the Urate Oxidase Reaction. J. Am. Chem. Soc. 122:11252–11253.

Schliep KP. 2011. phangorn: phylogenetic analysis in R. Bioinformatics 27:592–593.

Sharma V, Hiller M. 2020. Losses of human disease-associated genes in placental mammals. NAR Genomics Bioinforma. 2:lqz012.

Simic MG, Jovanovic SV. 1989. Antioxidation mechanisms of uric acid. J. Am. Chem. Soc. 111:5778–5782.

Singer MA. 2003. Do mammals, birds, reptiles and fish have similar nitrogen conserving systems? Comp. Biochem. Physiol. B Biochem. Mol. Biol. 134:543–558.

Steiner RA, Lebedev AA, Murshudov GN. 2003. Fisher’s information in maximum-likelihood macromolecular crystallographic refinement. Acta Crystallogr. D Biol. Crystallogr. 59:2114–2124.

Studer RA, Dessailly BH, Orengo CA. 2013. Residue mutations and their impact on protein structure and function: detecting beneficial and pathogenic changes. Biochem. J. 449:581–594.

True JR, Carroll SB. 2002. Gene Co-Option in Physiological and Morphological Evolution. Annu. Rev. Cell Dev. Biol. 18:53–80.

Vagin AA, Steiner RA, Lebedev AA, Potterton L, McNicholas S, Long F, Murshudov GN. 2004. REFMAC5 dictionary: organization of prior chemical knowledge and guidelines for its use. Acta Crystallogr. D Biol. Crystallogr. 60:2184–2195.

Vitting-Seerup K, Sandelin A. 2019. IsoformSwitchAnalyzeR: analysis of changes in genome-wide patterns of alternative splicing and its functional consequences.Berger B, editor. Bioinformatics 35:4469–4471.

Wang M, Gao J, Müller P, Giese B. 2009. Electron Transfer in Peptides with Cysteine and Methionine as Relay Amino Acids. Angew. Chem. Int. Ed. 48:4232–4234.

Winn MD, Ballard CC, Cowtan KD, Dodson EJ, Emsley P, Evans PR, Keegan RM, Krissinel EB, Leslie AGW, McCoy A, et al. 2011. Overview of the CCP4 suite and current developments. Acta Crystallogr. D Biol. Crystallogr. 67:235–242.

Wolff SP. 1994. [18] Ferrous ion oxidation in presence of ferric ion indicator xylenol orange for measurement of hydroperoxides. In: Methods in Enzymology. Vol. 233. Oxygen Radicals in Biological Systems Part C. Academic Press. p. 182–189. Available from: https://www.sciencedirect.com/science/article/pii/S0076687994330212

Wright PA. 1995. Nitrogen excretion: three end products, many physiological roles. J. Exp. Biol. 198:273–281.

Wu XW, Lee CC, Muzny DM, Caskey CT. 1989. Urate oxidase: primary structure and evolutionary implications. Proc. Natl. Acad. Sci. 86:9412–9416.

Yu G. 2020. Using ggtree to Visualize Data on Tree-Like Structures. Curr. Protoc. Bioinforma. 69:e96.

Zanotti G, Cendron L, Ramazzina I, Folli C, Percudani R, Berni R. 2006. Structure of zebra fish HIUase: insights into evolution of an enzyme to a hormone transporter. J. Mol. Biol. 363:1–9.

