## Supplemental figures for "Cysteine enrichment mediates co-option of uricase in reptilian skin and transition to uricotelism"

A

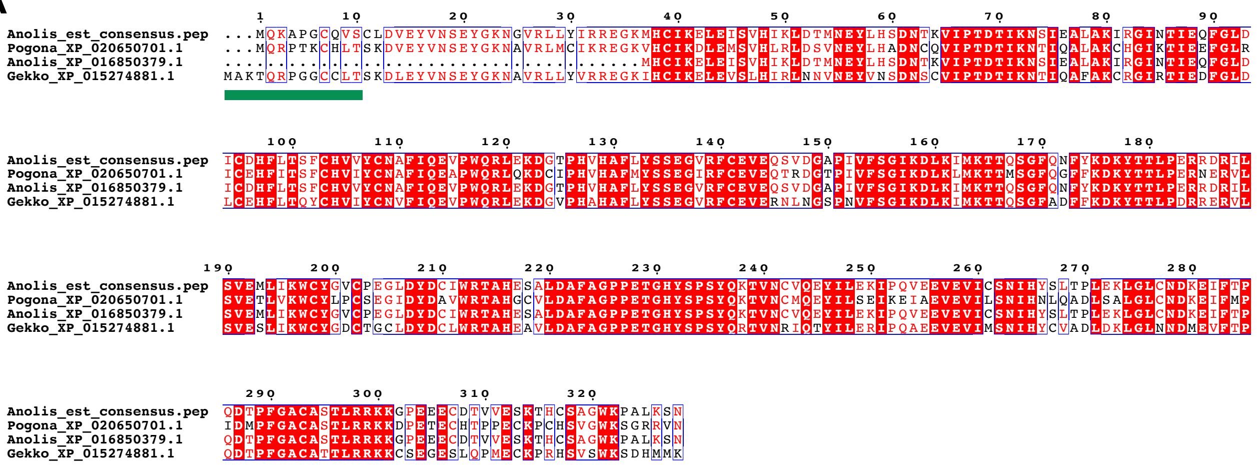


B

>Anolis_est_consensus.fa

TCCAAGCTGAAGCACTGTGAGTCGCTATTACTTTCCAGTGCCATCTCCAAGAGACGAAA**ATGCAGAAAGCACCGGGGTGCCAGGTTTCTTGCCTGGATGTGGAATACGT**
**GAATTCTGAATATGGAAAAAATGGAGTGCGACTCCTGTATATTAGAAGAGAAGGCAAGATGCATTGCATCAAAGAACTGGAAATCTCTGTACATATAAAACTGGACACT**
**ATGAATGAATACCTGCATTCTGATAACACCAAAGTTATCCCCACTGACACCATAAAGAATTCAATTGAAGCTCTAGCCAAAATTCGTGGGATCAACACGATAGAACAGT**
**TTGGCCTTGACATCTGTGATCACTTCCTCACCTCATTTTGCCACGTTGTGTATTGCAATGCCTTCATCCAAGAGGTGCCATGGCAGCGCCTAGAAAAGGATGGTACCCC**
**ACATGTACATGCTTTTCTGTATAGCTCTGAAGGGGTTCGATTTTGTGAAGTAGAACAGTCTGTGGATGGTGCACCAATTGTTTTCTCTGGCATCAAAGATCTGAAAATT**
**ATGAAAACAACCCAATCTGGATTTCAAAACTTCTACAAGGATAAGTACACCACACTTCCAGAAAGAAGAGATAGAATTCTCTCAGTGGAAATGTTGATAAAGTGGTGCT**
**ATGGTGTATGCCCAGAAGGCCTTGACTATGACTGTATATGGAGAACTGCACATGAAAGTGCCCTTGATGCCTTTGCTGGACCACCTGAAACAGGCCATTACTCCCCTTC**
**TTACCAGAAGACTGTCAACTGTGTTCAGGAATACATCCTGGAAAAAATCCCACAGGTGGAAGAAGTGGAAGTGATCTGTTCCAACATCCATTATTCACTGACACCTCTA**
**GAAAAACTGGGGTTGTGCAATGACAAGGAGATCTTTACACCTCAGGATACACCCTTTGGGGCTTGTGCATCAACGCTGCGTAGGAAGAAGGGCCCAGAAGAAGAGTGTG**
**ACACTGTCGTAGAGAGCAAAACCCACTGCTCTGCTGGATGGAAGCCTGCCCTGAAGTCGAAT**TAAGCAGTCAACGACATTATTACAATTATCTCATATCAGCATGTATT
ATTTACTCCACGCTTTGTCCCCATAATGCTCAACGTGTAGATATGAGTGATGTTTTCAAAAGTAGCCTAAAAGAAACCCTATGCTAACATCATAGTTCATTTTCTTTTG
TTTCTTTTCAAAATACAAGTAAGAACTGTTCAACGCAGCTCACTAGTACACATTTGAAGCTGGACATATGTAGAATCAAATCATTGAAAGTAGTGGGATTTAAGTTGGT
TAAATGTGACTTCTTTCAGTTCTATTGATTCCAGTGATCTTTTTACATCAAGGCTATATCTGGATCCACCTAAATAGTTTAGCAGGCTTCAAATATGAATATGTTGCTA
CATGGGGGGTTTTCACATCTGAAGCCTCCTGGGTTTTTTCCCCCTTGTAAGTGAATCCTACTAAAATATTAAGAAGTTCTACAAAAAAAAAAAAAAAA

**S1 Fig. Evidence for a complete Uox coding sequence in *Anolis carolinensis*.** (A) Alignment of the translated CDS with database Uox sequences of *Pogona vitticeps* (XP_020650701.1), *Gekko japonicus* (XP_015274881.1) and the N-truncated *A. carolinensis* sequence (XP_016850379.1). The amino acid sequence corresponding to the first exon of *G. japonicus Uox* (**fig. 2A**) is indicated by a green bar. (B) Consensus mRNA obtained from the multiple alignment of *A. carolinensis* sequences identified through homology search (tblast) in dbEST. The putative CDS is in bold.

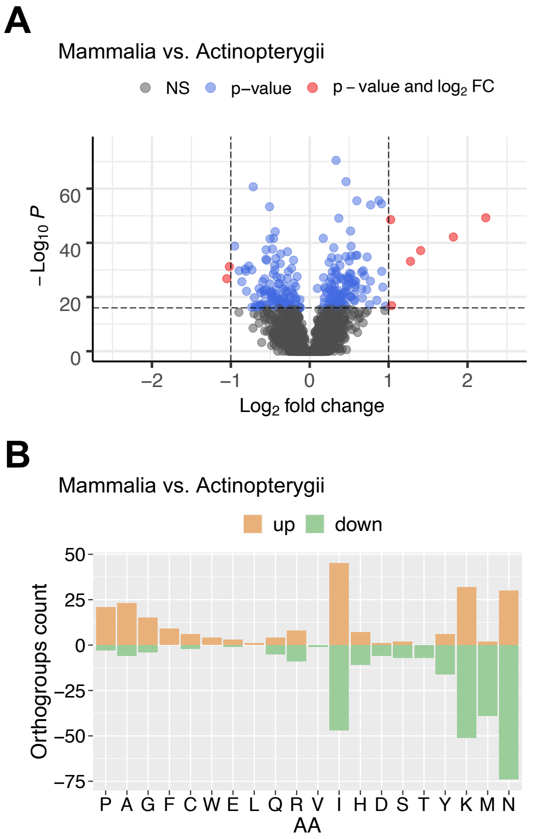


**S2 Fig. Variation in amino acid composition of vertebrate orthologous proteins.** (A) Volcano plot depicting the cysteine content variation of orthologous proteins between groups of vertebrates: mammals vs. fishes. Horizontal dashed line indicates p-value of $1e^{-16}$; vertical dashed lines indicate $\log_2$ fold change of −1 and 1. (B) Bar plot illustrating the number of orthogroups in which the content of the corresponding amino acid is significantly increased (light orange) or decreased (green) according to p-value < $1e^{-16}$ and $\log_2$ fold change > ±1 in mammals vs. fishes.

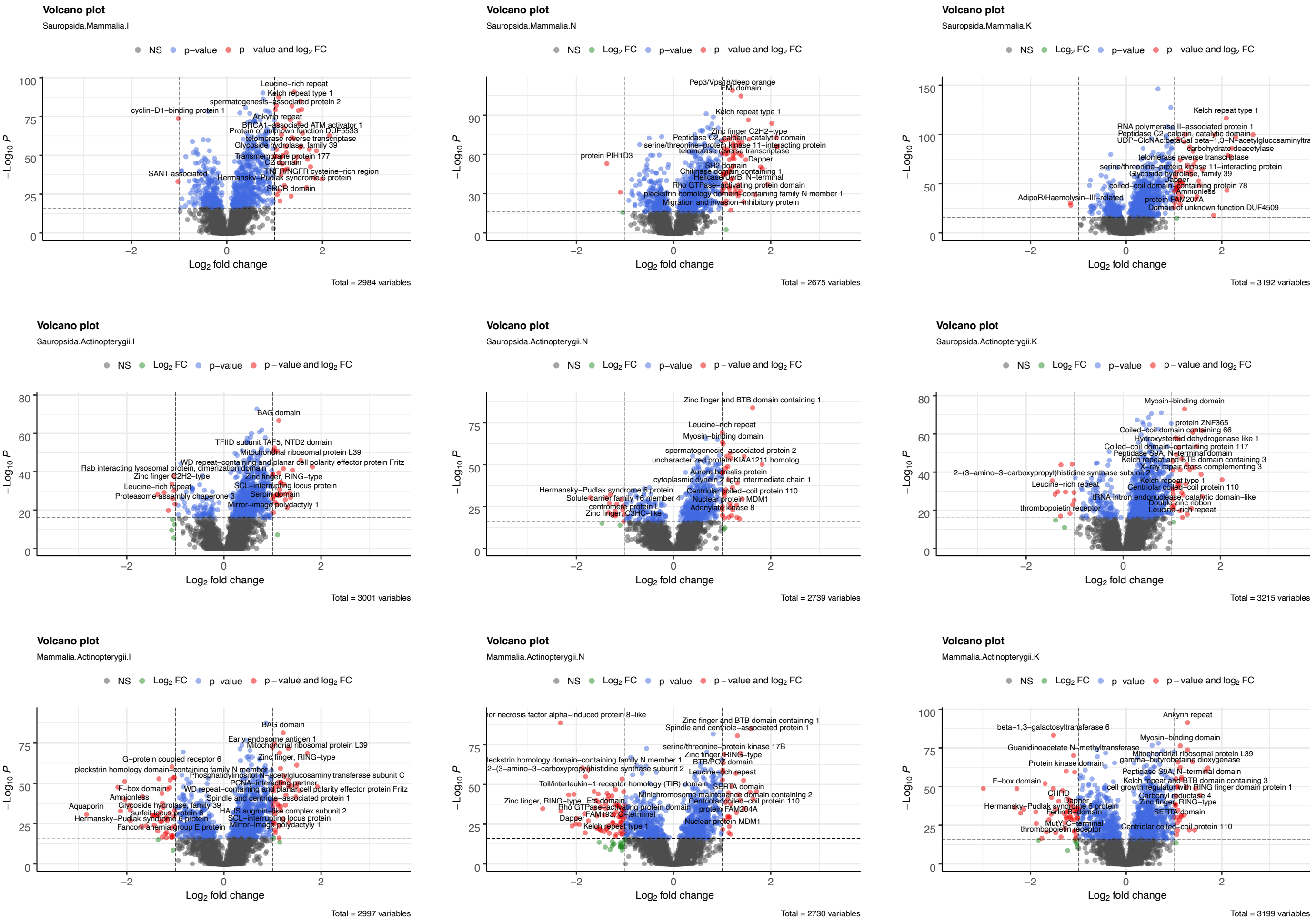


**S3 Fig. Variation in I, N, K amino acid composition of vertebrate orthologous proteins.** (A) Volcano plot depicting the "INK" content variation of orthologous proteins between groups of vertebrates: sauropsids vs. mammals (upper panels), sauropsids vs. fishes (middle panels), and mammals vs. fishes (lower panels). Horizontal dashed line indicates p-value of $1e^{-16}$; vertical dashed lines indicate $\log_2$ fold change of −1 and 1.

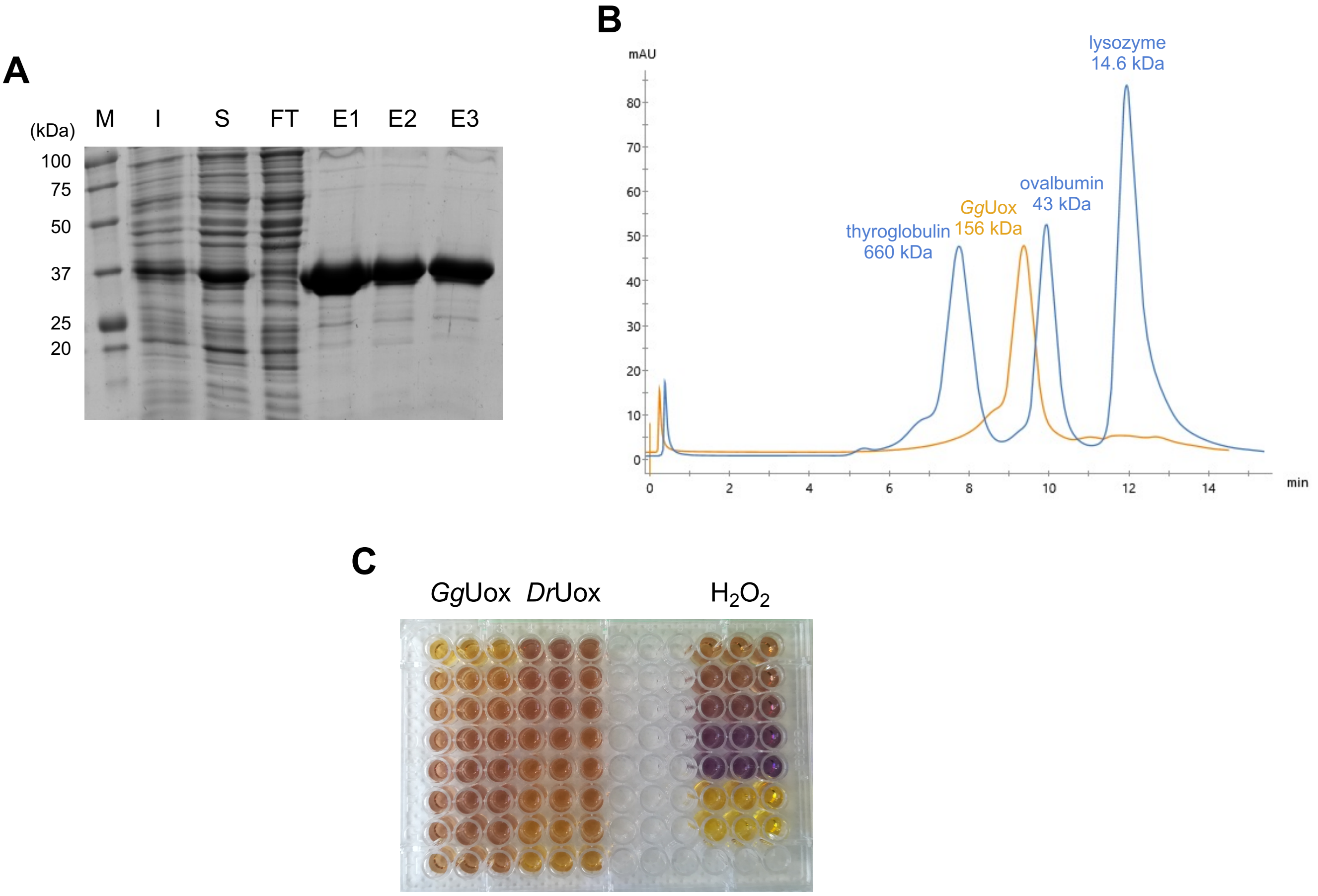


**S4 Fig. Purification and characterization of recombinant *Gg*Uox.** (A) SDS-PAGE of the purification of *Gg*Uox by $Ni^{2+}$-affinity chromatography. M: marker; I: post-induction total cell fraction; S: soluble cell fraction; FT: flow-through; E1-E3: elution fractions. (B) Size-exclusion chromatography (Superose 6 Increase 10/300 column) profile of purified *Gg*Uox (light orange) and of protein markers (light blue). (C) Measurement of hydrogen peroxide generated in the urate oxidation reaction catalysed by *Gg*Uox or *Dr*Uox. Reactions were carried out under single turnover conditions with 100 mM potassium phosphate (pH 7.6), 30 µM enzyme, 25 µM urate. The amount of hydrogen peroxide was quantified by Ferrous Oxidation Xylenol Orange (FOX) assay.

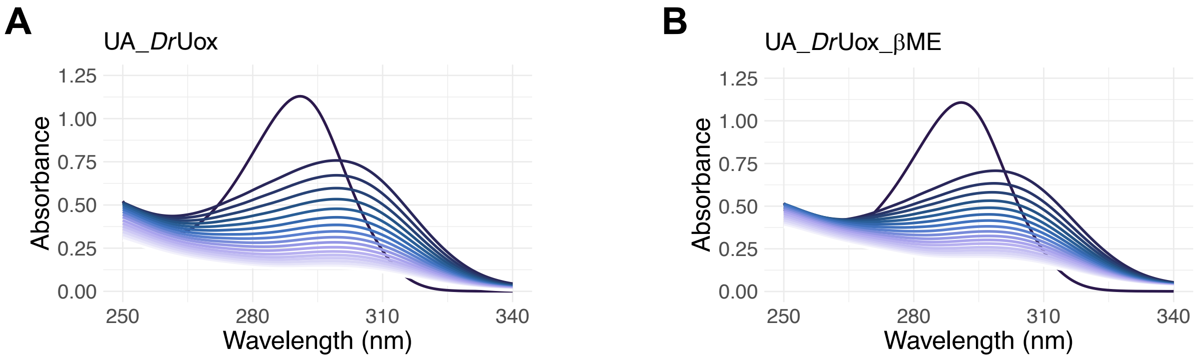


**S5 Fig. Time-resolved UV-Vis spectra showing urate oxidation by *Dr*Uox.** (A-B) Reaction contained 100 mM potassium phosphate (pH 7.6), 100 μM urate, 30 μM *Dr*Uox, and (A) 0 or (B) 300 μM β-mercaptoethanol (βME). Spectra were acquired every 1 min at 25 °C.

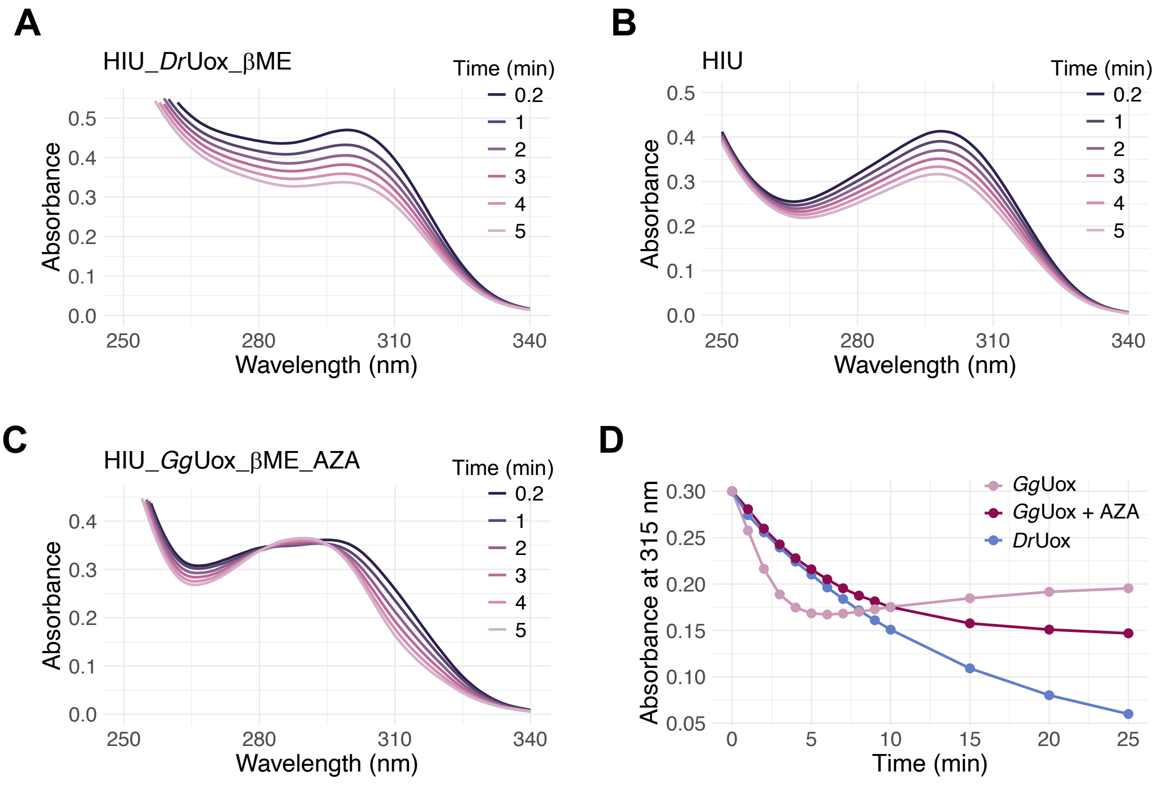


**S6 Fig. *Gg*Uox-catalyzed or spontaneous HIU degradation.** (A-C) Time-resolved UV-Vis spectra showing HIU degradation. Reaction contained 100 mM potassium phosphate (pH 7.6), 60 μM HIU, and (A) 30 μM *Dr*Uox, 300 μM β-mercaptoethanol (βME), or (C) *Gg*Uox (30 μM), 300 μM βME, and 8-azaxanthine (AZA; 50 μM). Spectra were acquired at 25 °C. (D) Time-dependent change of absorbance at 315 nm during HIU degradation by *Gg*Uox (pink), *Gg*Uox with AZA (bordeaux), and *Dr*Uox (light blue).

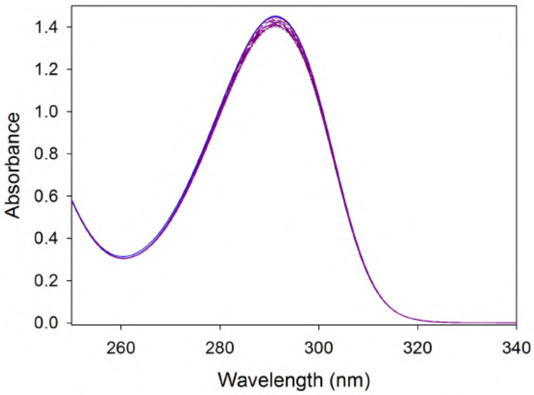


**S7 Fig. Time-resolved UV-Vis spectra of urate oxidation by *Gg*Uox in the presence of 8-azaxanthine (AZA).** Reaction contained 100 mM potassium phosphate (pH 7.6), 100 μM urate, 10 μM *Gg*Uox, 50 μM AZA and 1 μM Alls. Spectra were acquired every 1 min at 25 °C.

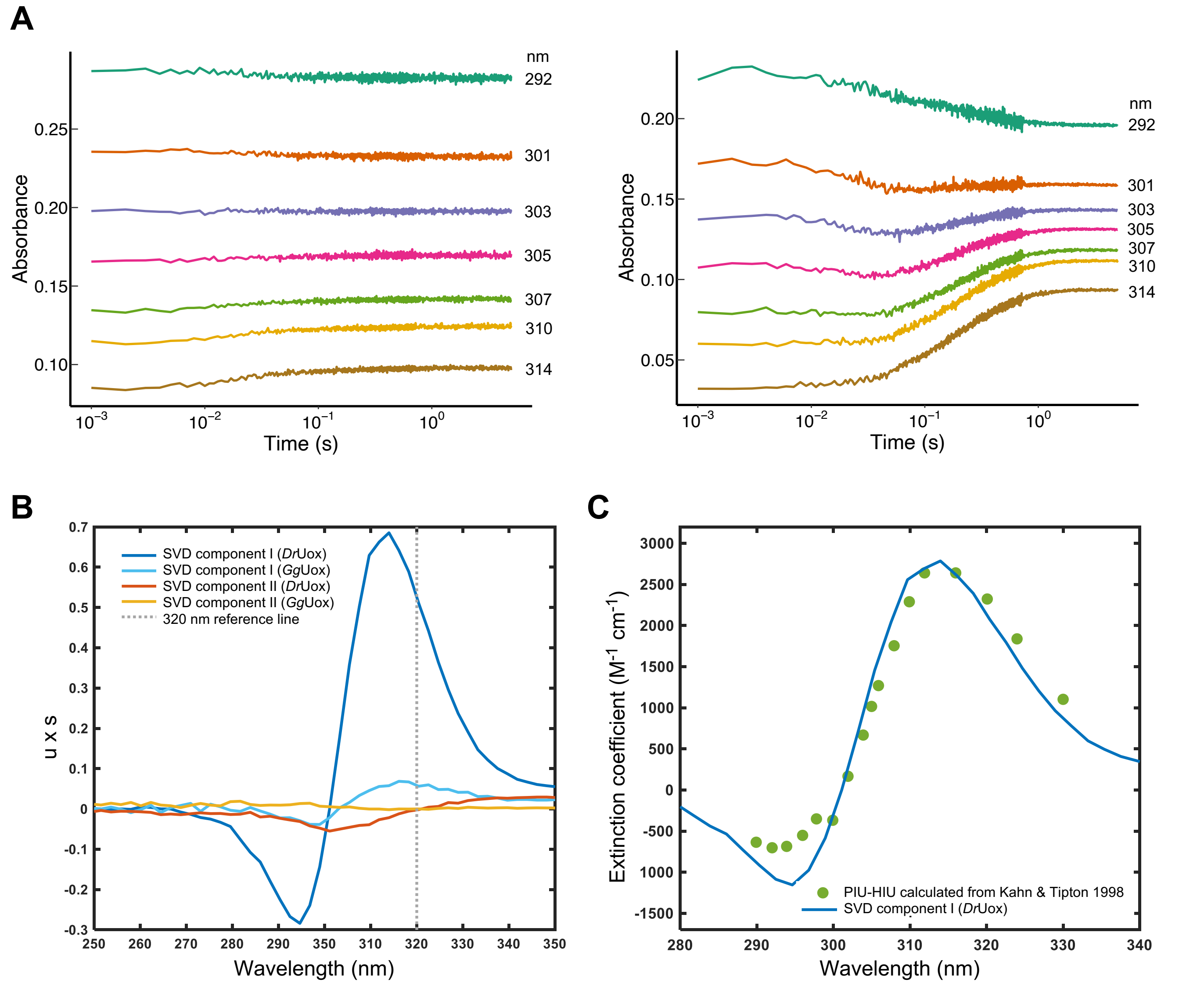


**S8 Fig. Stopped flow analysis of *Gg*Uox and *Dr*Uox reactions.** (A) Stopped flow kinetics at selected wavelengths of *Gg*Uox (left) or *Dr*Uox (right). Reactions were carried out under single turnover conditions at 25 °C with 100 mM potassium phosphate (pH 7.6), 35 µM enzyme, 25 µM UA. To improve signal to noise ratio 3 mixing kinetics were averaged. (B) Main spectral components obtained from the SVD analysis on the two time-resolved spectral series following the reaction between UA and *Gg*Uox (light blue: SVD component I; yellow: SVD component II) or *Dr*Uox (blue: SVD component I; orange: SVD component II) . Spectral components are reported as the product of the spectral component (u) multiplied by its corresponding relative weight (s). (C) Difference spectrum (green closed circles) between 5-peroxyisourate (PIU; the second intermediate of Uox reaction) and HIU (experimentally determined spectra were extracted from Kahn and Tipton 1998), and SVD component I spectrum (blue line) from *Dr*Uox SVD analysis. Data are normalized for UA concentration.

**A**

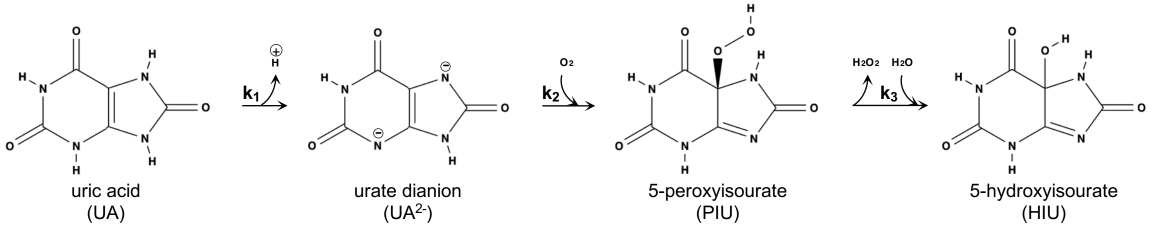


**B**

| | $k_1$ ($s^{-1}$) | $k_{-1}$($s^{-1}$) | $k_2$($s^{-1}$) | $k_{-2}$($s^{-1}$) | $k_3$($s^{-1}$) | $k_{-3}$($s^{-1}$) |
|---|---|---|---|---|---|---|
| ***Gg*Uox** | 68384±160110 | 0.45±24.83 | 29.77±45.77 | 0.11±59.20 | 2768±85984 | 9732±32073 |
| ***Dr*Uox** | 87643±246 | - | 20242±43 | - | 4.11±0.02 | - |

**S9 Fig. Kinetics parameters determination of *Gg*Uox and *Dr*Uox activity upon model fitting.** (A) Three-steps irreversible model for UA to HIU conversion. Intermediate species were already attributed to urate dianion ($UA^{2-}$) and 5-peroxyisourate (PIU) (Kahn and Tipton 1998). (B) Kinetic constants obtained from stopped flow data fitting to SimBiology models. *Gg*Uox data were fitted to the reversible three step model shown in Fig 4J; *Dr*Uox data were fitted to the irreversible three step model shown in (A).

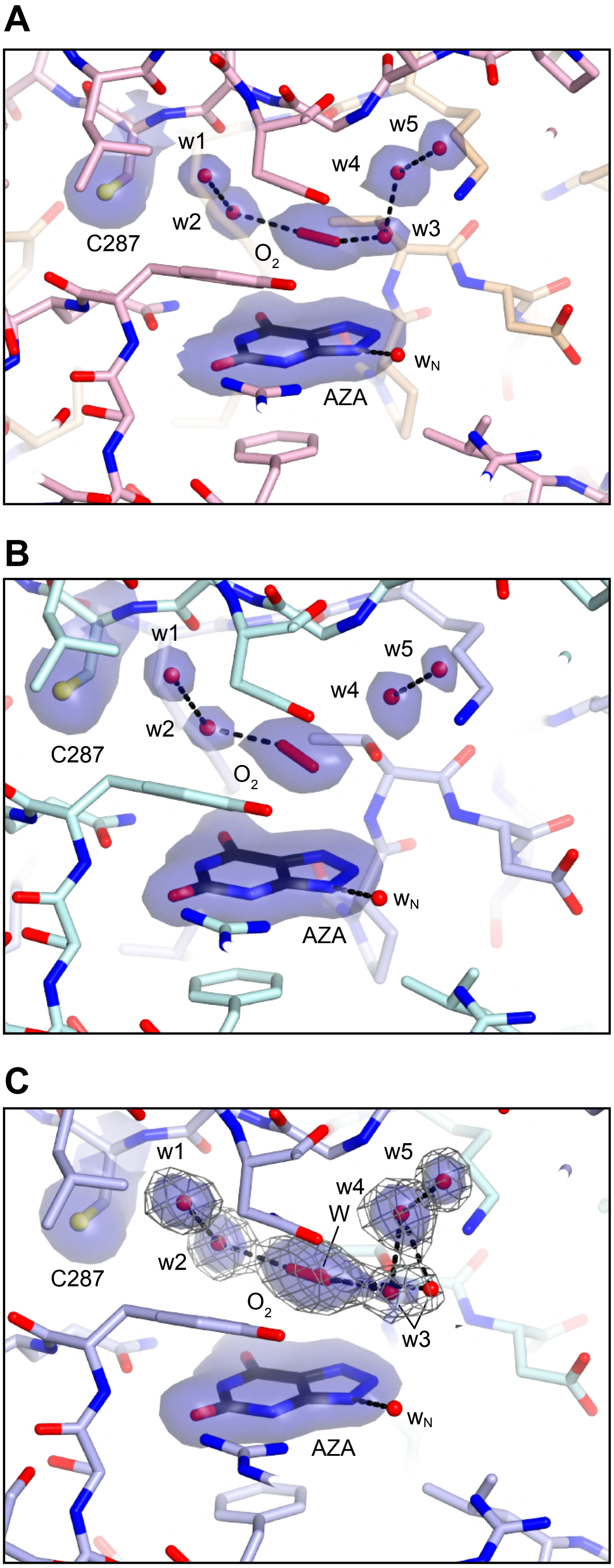


**S10 Fig. *Gg*Uox active sites in the tetramer.** Stick representation of the active sites (A) between chain B and chain A, (B) between chain C and chain D, and (C) between chain D and chain C with the four protomers highlighted by different colors. $2mF_o$-$DF_c$ electron density map for AZA, solvent molecules in its proximity, and C287 is shown in blue at the $+1.0\sigma$ level. In (C) the electron density for the solvent above AZA is also shown as chicken-wire representation at the $+0.5\sigma$ level in grey. Selected hydrogen bonds are highlighted in black as broken lines. In general, the solvent structure is quite conserved and best explained by a mixture of dioxygen and water molecules. Water molecules w1, w2, w4, w5 above AZA appear well defined whilst water molecule w3 is rather mobile and in one active site (B) is not visible. In another active site (C) w3 occupies two alternative positions with dioxygen and a solvent molecule (W) sharing the same position with partial occupancy.

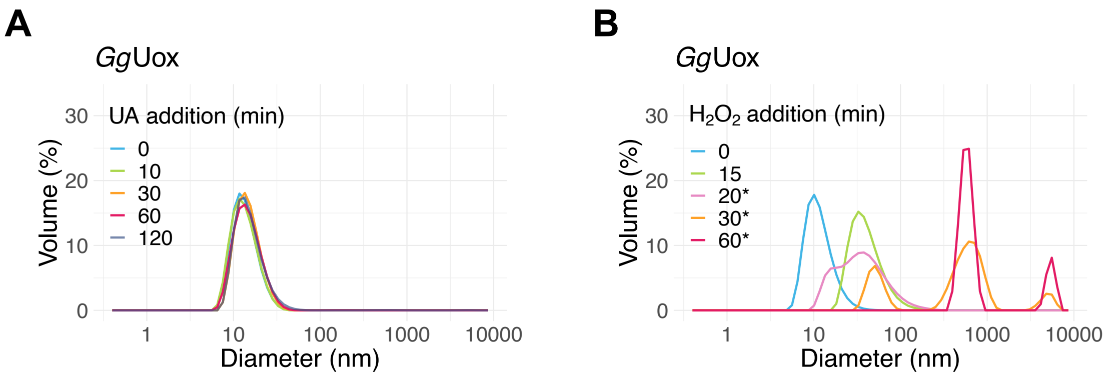


**S11 Fig. DLS analysis of the size of *Gg*Uox in solution.** (A) Samples containing 100 mM potassium phosphate (pH 7.6) and 20 µM *Gg*Uox were incubated at RT and analysed at different time points after the addition of 100 µM urate (UA). (B) Samples containing 100 mM potassium phosphate (pH 7.6) and 20 µM *Gg*Uox were incubated at RT and analysed at different time points after the addition of 100 µM urate (UA). Samples containing 100 mM potassium phosphate (pH 7.6) and 7 µM *Gg*Uox were incubated at RT and analysed at different time points after the addition of 100 µM $H_2O_2$. After 16 min, 5 mM β-mercaptoethanol was added; samples measured at 20, 30, and 60 minutes, containing β-mercaptoethanol are indicated with *.

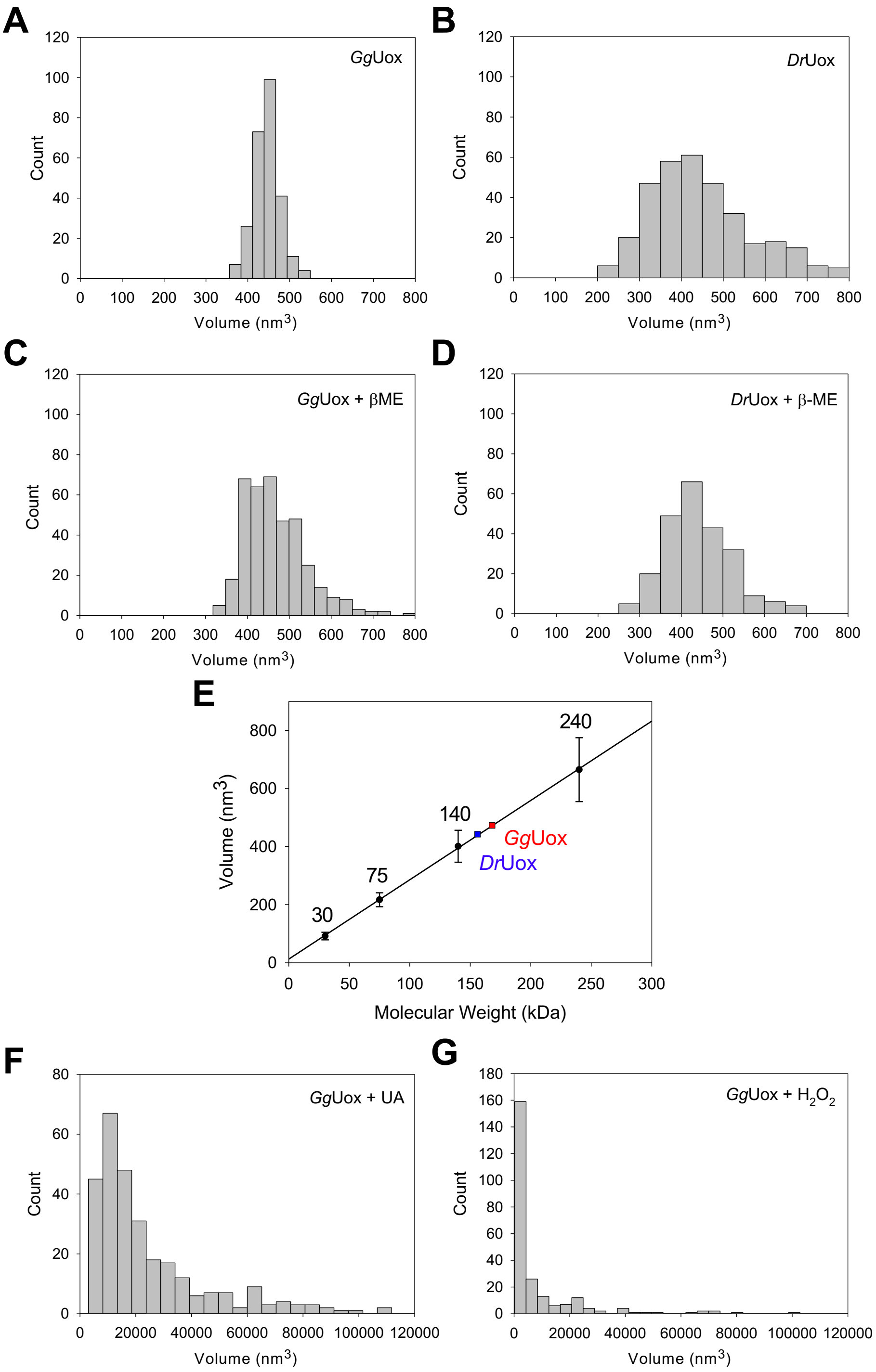


**S12 Fig. AFM analysis of the size of *Gg*Uox and *Dr*Uox in solution.** Volume distribution of (A) native *Gg*Uox, (B) native *Dr*Uox, (C) *Gg*Uox pre-incubated with β-mercaptoethanol (βME), and (D) *Dr*Uox pre-incubated βME. (E) Calibration curve used to infer the molecular weights of *Gg*Uox and *Dr*Uox. Data points correspond to: Carbonic anhydrase (30 kDa), Conalbumin (75 kDa), Alcohol dehydrogenase (140 kDa) and Catalase (240 kDa). The red square indicates the molecular weight of *Gg*Uox (156 kDa) based on the highest peak of the volume distribution in (A); the blue square indicates the molecular weight of *Dr*Uox (146 kDa) based on the highest peak of the volume distribution in (B). (F-G) Volume distribution of *Gg*Uox after incubation at RT with (F) urate (UA) or (G) hydrogen peroxide ($H_2O_2$). The highest peak in (F) corresponds to 10000 $nm^3$; the highest peak in (G) corresponds to 2000 $nm^3$.

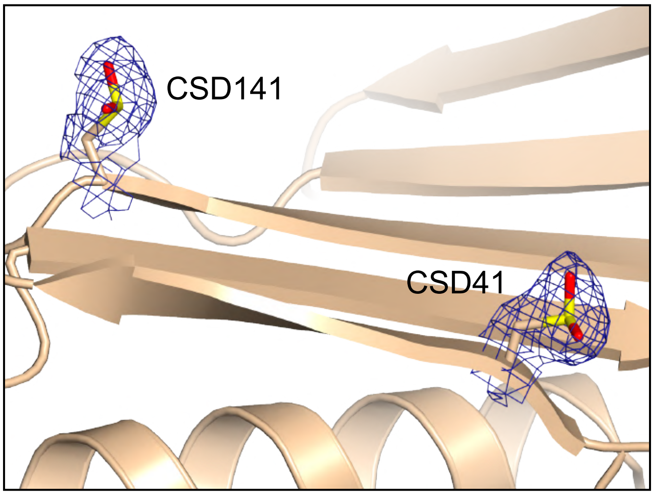


**S13 Fig. Oxidized cysteines in the *Gg*Uox X-ray structure.** Example of cysteine oxidation to its sulfinic form (CSD) following $H_2O_2$ treatment. $2mF_o$-$DF_c$ electron density map is shown in blue at the +1.0σ level.

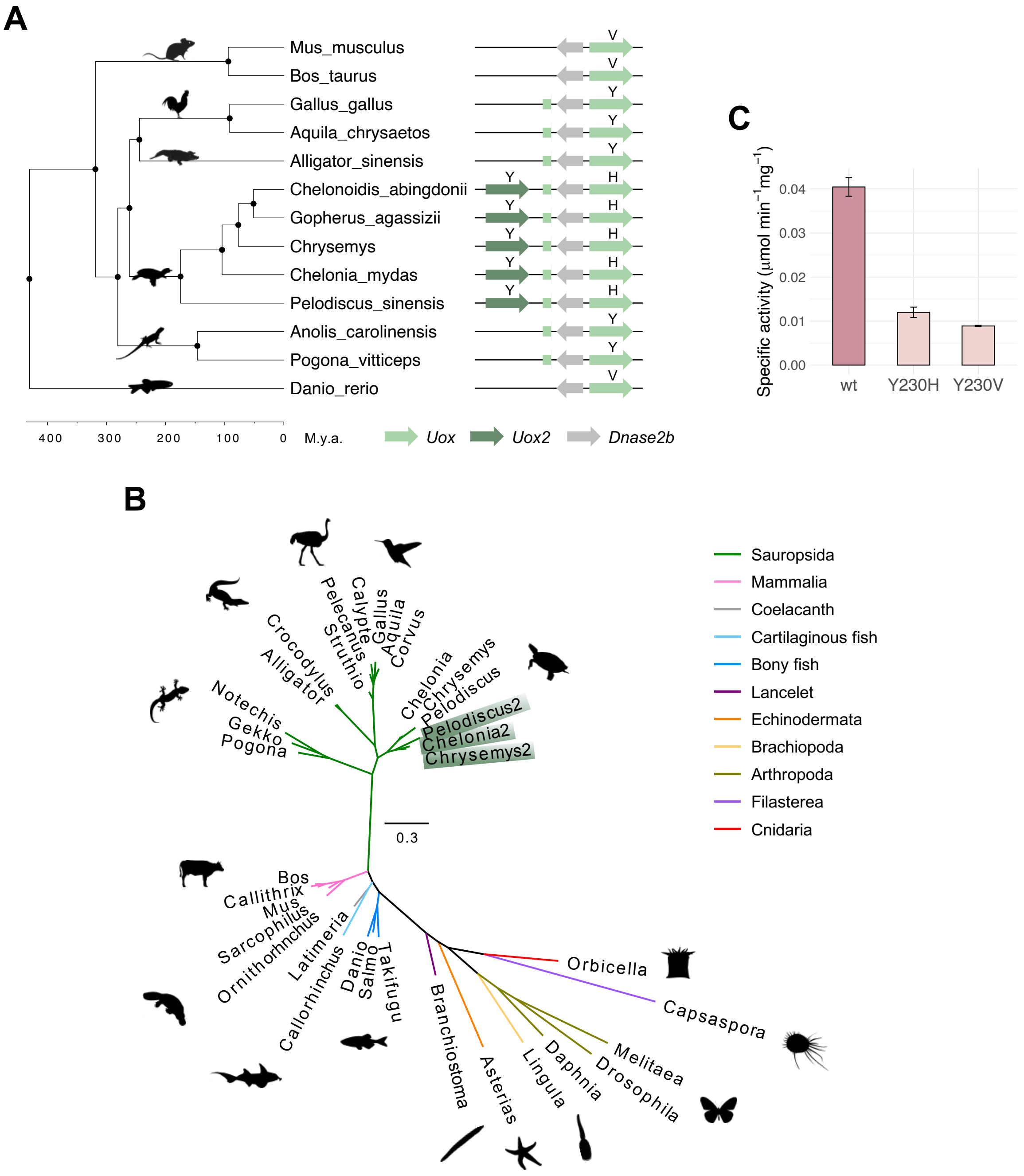


**S14 Fig. *Uox* gene duplication in chelonian reptiles.** (A) Chronogram of vertebrate phylogeny derived from TimeTree. Conserved synteny between *Uox* and *Dnase2b* genes in the different species and tandem duplication of *Uox* in chelonian reptiles are displayed at the terminal nodes. The 5'exon of reptilian *Uox* is represented by a green square before the 3'end of *Dnase2b* gene. The letter above the arrow corresponds to the amino acid at position 230 in the sequence of *Gg*Uox. (B) Maximum likelihood unrooted phylogenetic tree of Uox sequences from animal species. Genus names and silhouettes of one or more taxa for each taxonomic group (legend) are shown. Paralogous proteins in chelonian reptiles (Uox2) are highlighted in dark green. Scale bar, substitution/site. (C) Specific activity of *Gg*Uox wild-type (wt) and mutants Y230H, Y230V. Reactions were carried out in 100 mM potassium phosphate (pH 7.6), at 25 °C, with the addition of 0.25 µM HIUase. Error bars represent the standard deviation of three measurements.

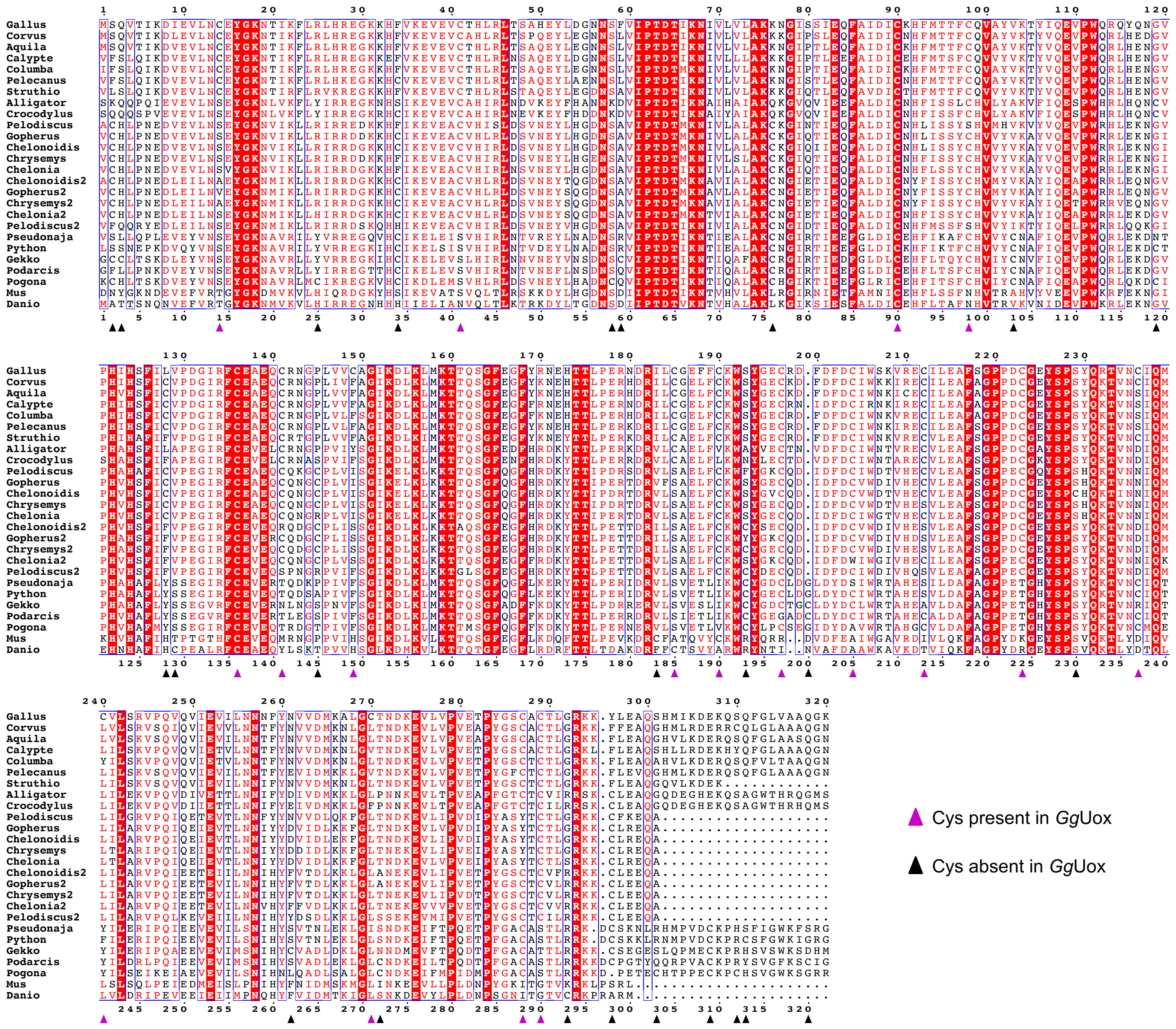


**S15 Fig. Multiple alignment of vertebrate Uox proteins that was used to infer the phylogenetic tree presented in Fig 8.** Shown is the portion of the alignment extending from the first to the last amino acid of *Gallus gallus* (*Gg*) Uox sequence. Residue numbering at the top refers to *Gg*Uox; numbering at the bottom refers to the alignment. Cysteine residues are marked by triangles coloured as indicated in the legend, and are the same shown in Fig 8.

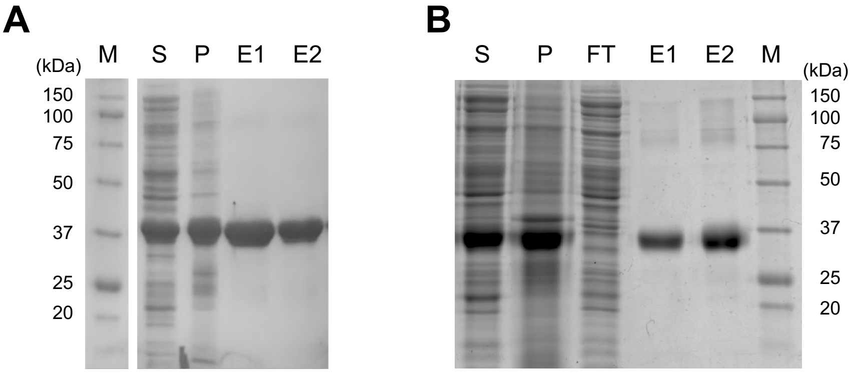


**S16 Fig. Recombinant expression of *Gg*Uox mutants.** SDS-PAGE of the purification of *Gg*Uox mutants (A) Y230H and (B) Y230V by $Ni^{2+}$-affinity chromatography. M: marker; S: soluble cell fraction; P: insoluble cell fraction (pellet); FT: flow-through; E1-E2: elution fractions.

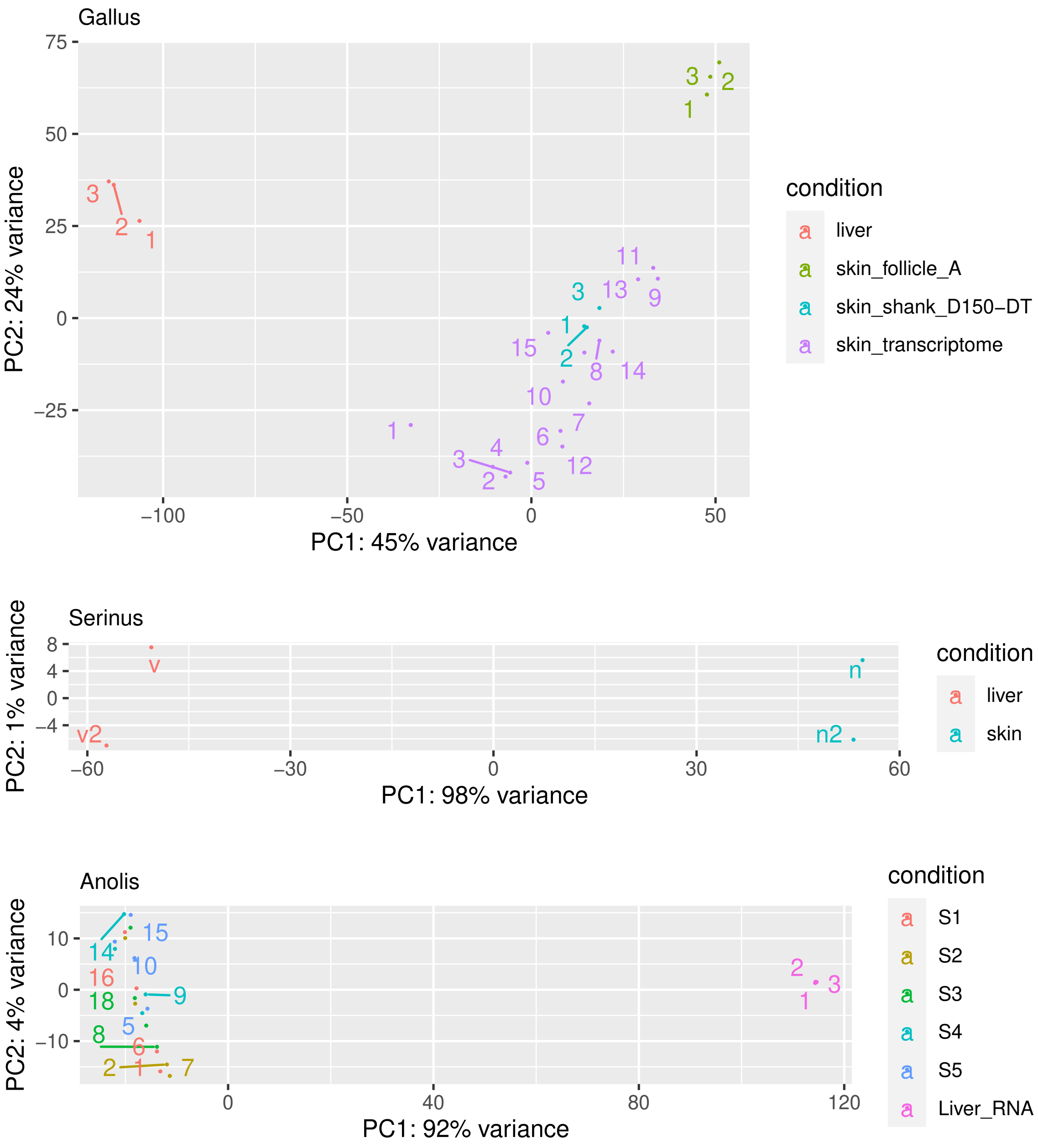


**S17 Fig. Principal component analysis (PCA) of the RNA-seq datasets used for the quantification of *Uox* expression.** Data belong to different tissues of *Gallus gallus* (upper panel), *Serinus canaria* (middle panel), and *Anolis carolinensis* (lower panel). PCA analysis was conducted with the variance stabilizing transformation (vst) and PCAplot functions of the DESeq2 R package based on transcript abundance value determined by the Kallisto software. Sequence Read Archive (SRA) accessions of the shown datasets are reported in S3 Table.

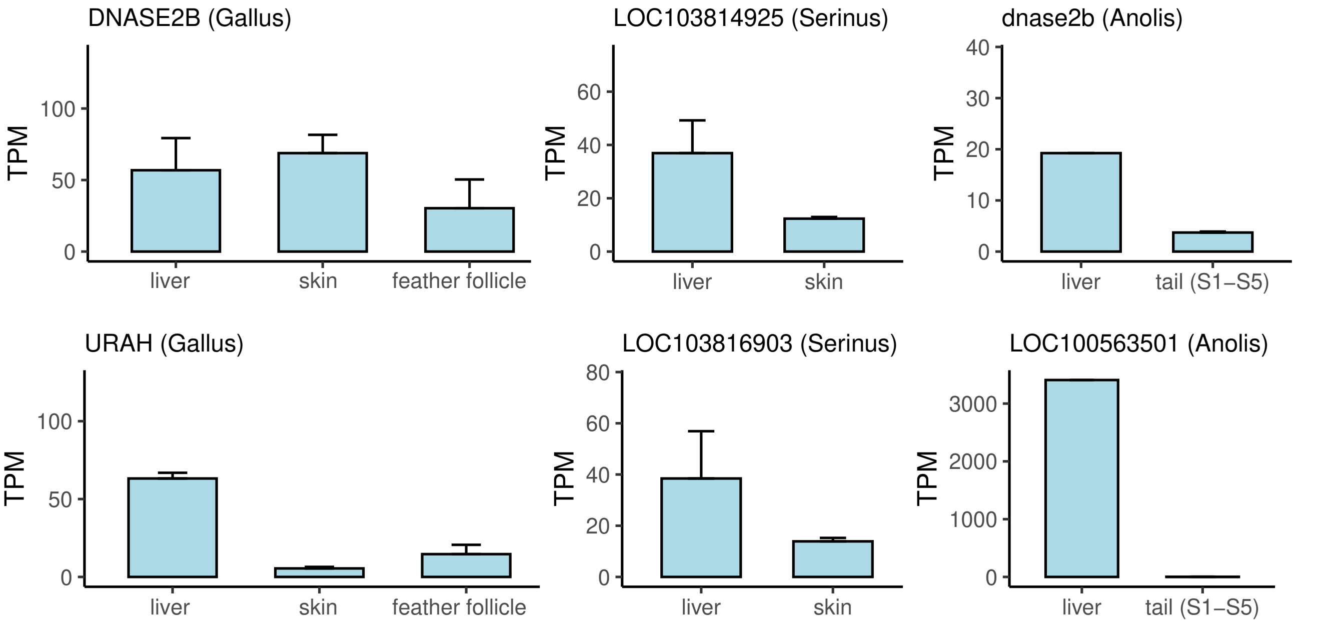


**S18 Fig. Expression of reptilian *Dnase2b* and *UraH*.** Expression levels (TPM: Transcripts Per kilobase Million) of *Dnase2b* (upper panels) and *Urah* (lower panels) in the liver and tegumental tissues (skin and feather follicle) of *Gallus gallus, Serinus canaria,* and *Anolis carolinensis* as derived from RNA-seq data analysis.

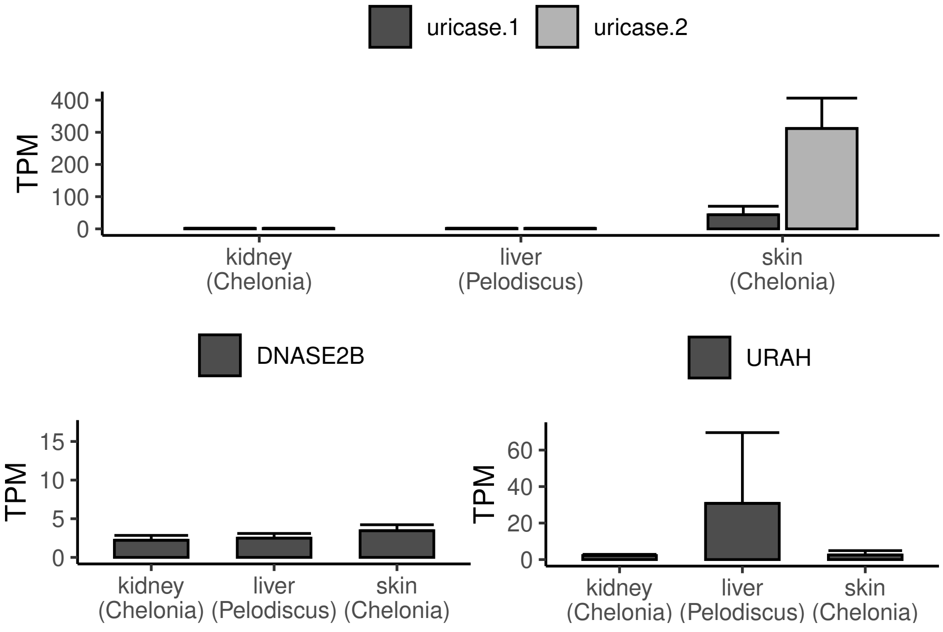


**S19 Fig. Expression of the two *Uox* paralogs, *Dnase2b*, and *Urah* in chelonians.** Expression levels (TPM: Transcripts Per kilobase Million) in the kidney and skin of *Chelonia mydas* and in the liver of *Pelodiscus sinensis*, of *Uox* (uricase.1 and uricase.2), *Dnase2b*, and *Urah*, as derived from RNA-seq data analysis.

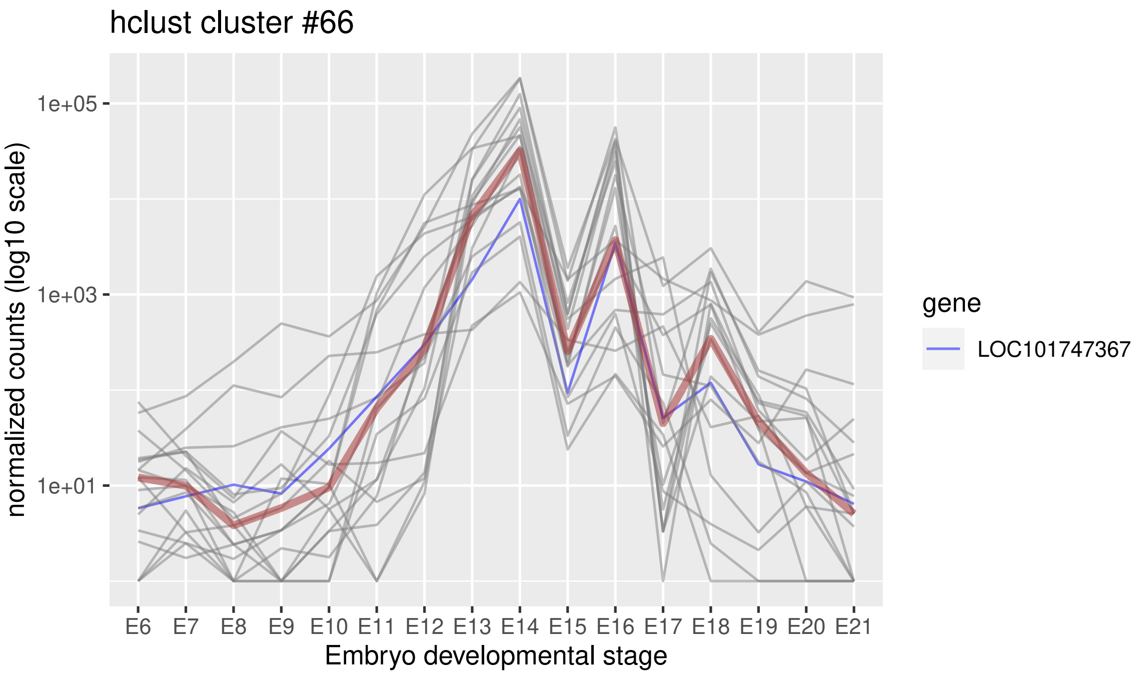


**S20 Fig. Expression profiles of *Gg*Uox co-expressed genes in skin embryo development.** Gene co-expression cluster was obtained by hierarchical clustering with the hclust function of the R package based on euclidean distances of scaled transcript abundance values. Transcript abundance was determined with Kallisto based on RNA-seq data of the PRJNA397795 Bioproject (S3 Table). Gene accession numbers and descriptions for individual traces (gray lines) are reported in Fig 7D. The median value of the cluster is indicated by a brown line.

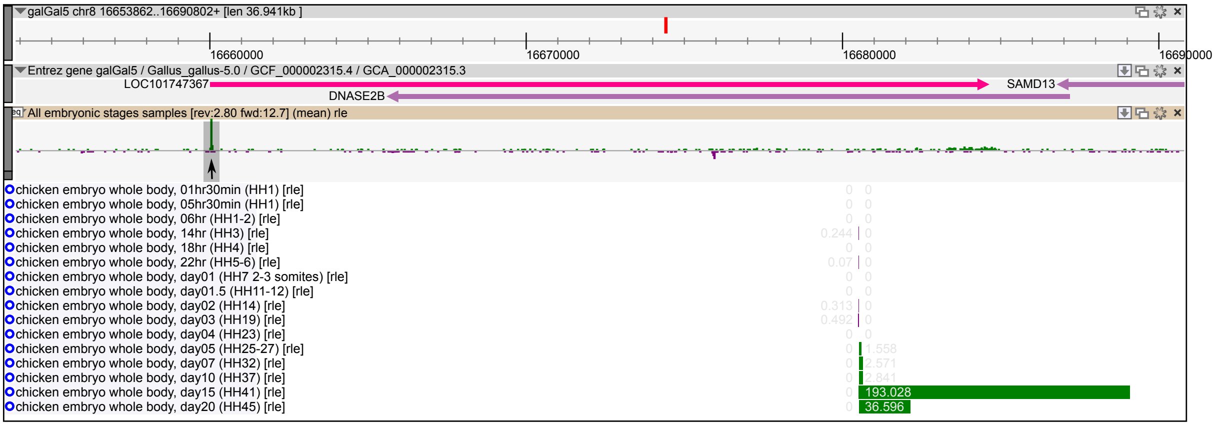


**S21 Fig. UOX transcription start site (TSS) peaks and expression levels during embryo development.** A strong single peak (arrow) is mapped to the 5’ end of *G. gallus* UOX (LOC101747367) through Cap Analysis of Gene Expression (CAGE) [PMC5600399] visualized by Chicken-ZENBU. Developmental stage-resolved analysis (bar graph) shows late stage-specific expression of UOX (Hamburger and Hamilton stage 41 [HH41], day 15).

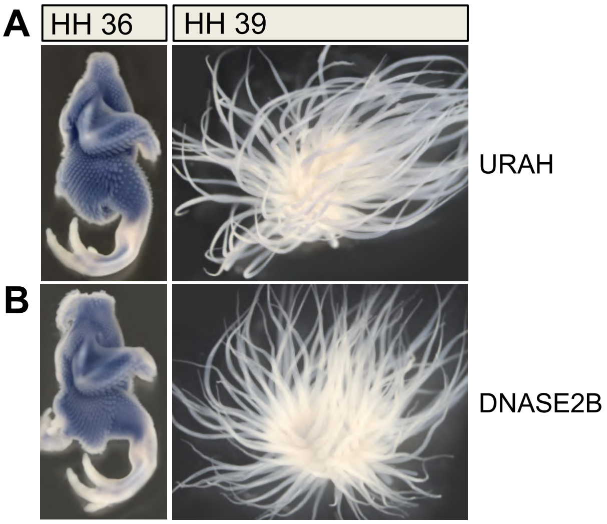


**S22 Fig. Expression of *Gallus gallus* URAH and DNASE2B during embryogenesis.** In situ hybridization analysis of URAH (upper panels) and DNASE2B (lower panels) expression in chick embryos at HH developmental stages 36 (left panels) and 39 (bottom panels).

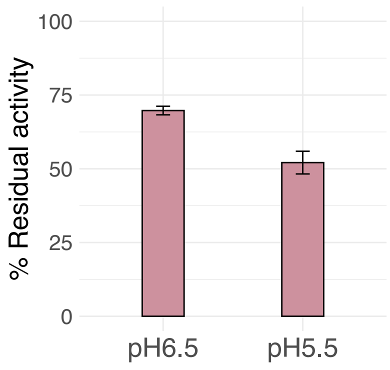


**S23 Fig. Uricase activity of *Gg*Uox at acidic pH.** Reactions were carried out in 100 mM potassium phosphate, pH 6.5 or 5.5, at 25 °C, with the addition of 1 μM HIUase. The activities are expressed in %, where 100% refers to the activity of *Gg*Uox measured at pH 7.6. Error bars represent the standard deviation of three measurements.

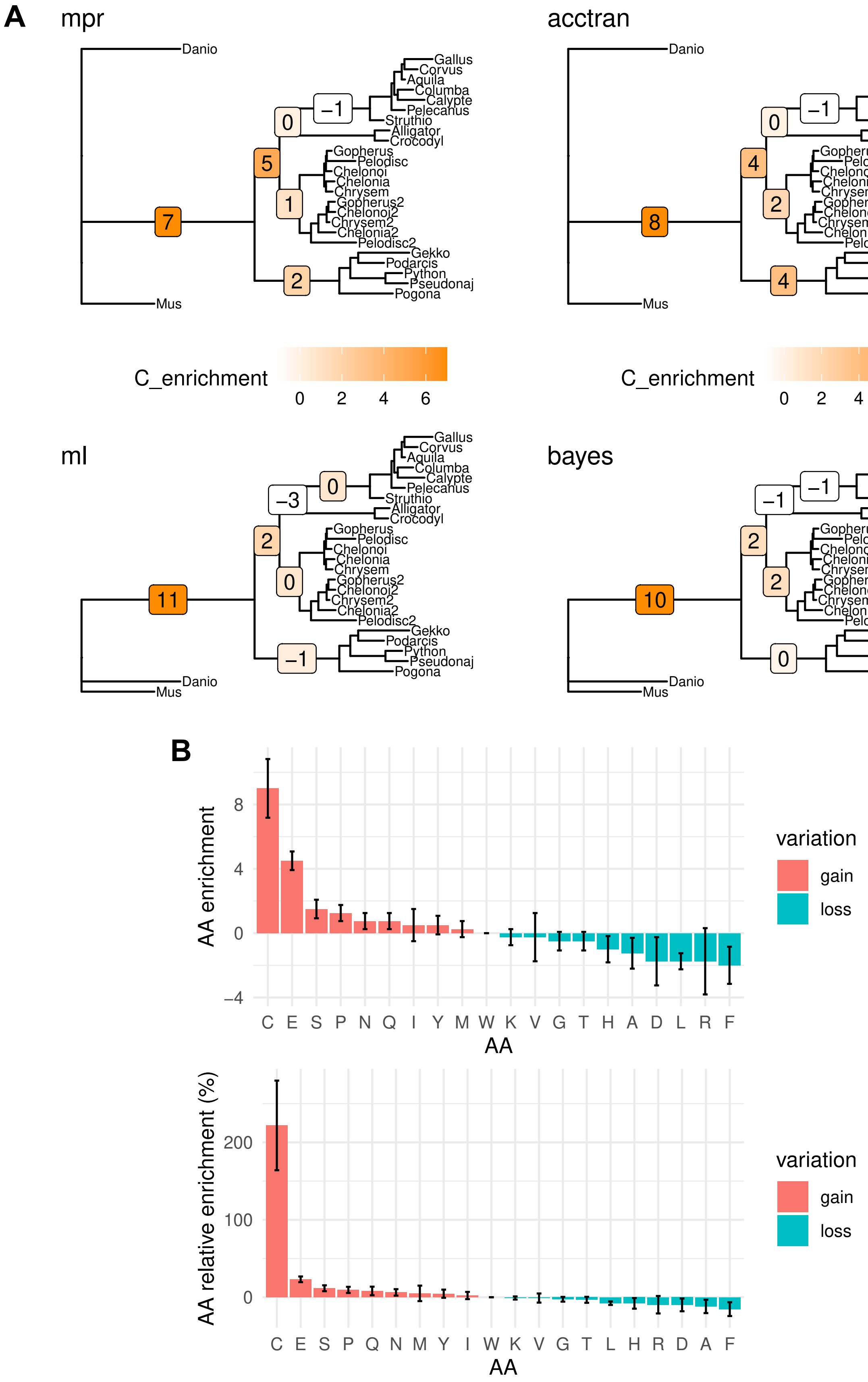


**S24 Fig. Cysteine enrichment in reptilian uricase.** (A) Net cysteine variation (gain minus loss) along selected nodes of the vertebrate uricase tree according to four different models implemented in Phangorn: maximum parsimony (mpr), mpr with accelerated transformation (mpr_acctran), maximum likelihood (ml), and highest posterior probability (bayes). (B) Absolute (top) and relative (bottom) amino acid enrichment in the uricase sequence of the stem reptile. Data are mean and standard deviation of the four different models in (A).

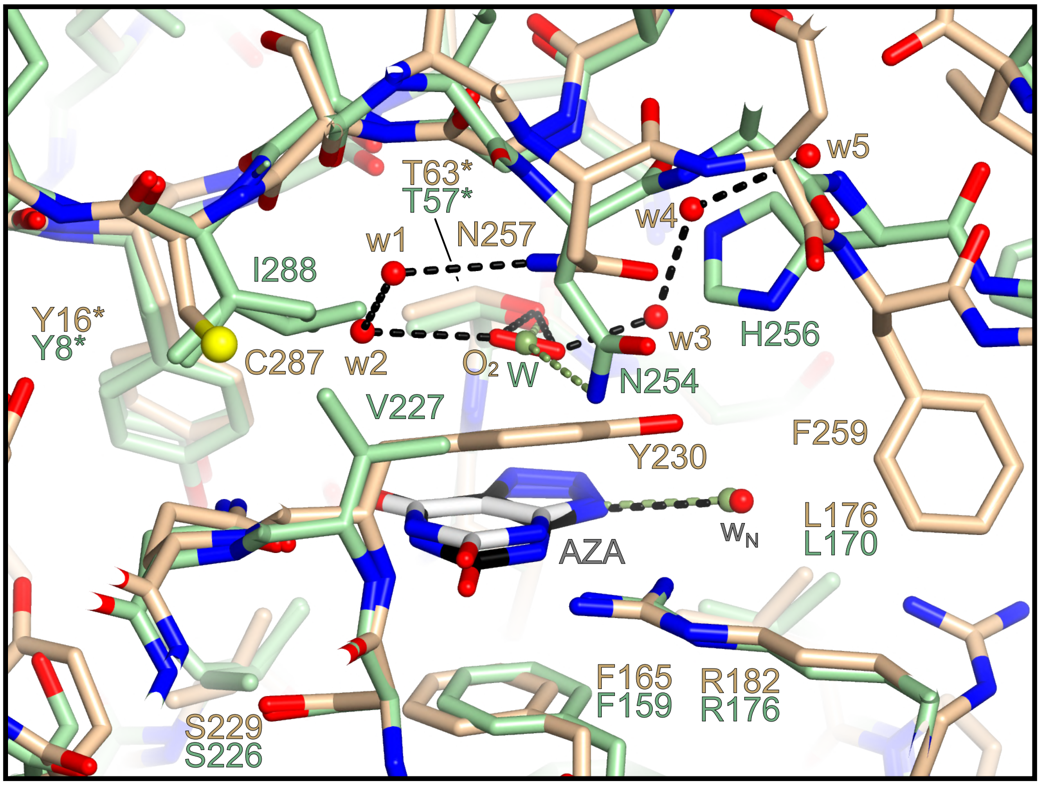


**S25 Fig. *Gg*Uox and *Aspergillus flavus* (*Af*) Uox active sites.** Superposition of the active sites of *Gg*Uox (wheat) and *Af*Uox (light green) in the AZA (grey and black carbon atoms for *Af*UOX and *Gg*Uox, respectively) bound state. Residues in the different orthologs are highlighted using the same color scheme. Solvent molecules in *Gg*Uox ($w_1$-$w_5$, $w_N$, and $O_2$) and *Af*Uox (W and $w_N$) are shown in red and light green, respectively. Asterisks indicate residues belonging to different protomers. Although several residues are conserved between *Gg*Uox and *Af*Uox including the AZA-bound $w_N$, amino acid differences are observed at important topological positions. The V227→Y230 replacement forces an alternative conformation for *Gg*Uox N257 side chain that is H-bonded to $w_1$. In *Af*Uox this residue is involved in the coordination of the only water molecule (W) that occupies the 'peroxo hole' and is also stabilised by the side chain of T57*. Additionally, H256→F259 and I288→C287 replacements increase the volume of the 'peroxo hole' that accommodates under normoxic conditions a more complex solvent structure constituted by a mixture of $O_2$ and water molecules. Coordinates for *Af*Uox were taken from the main conformation of the high-resolution crystal structure jointly refined using neutron and X-ray data (PDB code 7A0L) (McGregor et al., 2021).

**S1 Table. Orthogroups in which the cysteine content is significantly different between groups of vertebrates: Sauropsida (S), Mammalia (M), and Actinopterygii (A).**

| Orthogroup ID | Orthogroup name | Comparison | p-value | fold-change |
|---|---|---|---|---|
| 305356at7742 | Uricase | S-M | 8.154863e-81 | 1.75831596 |
| 266521at7742 | C2 domain | S-M | 1.021959e-37 | -1.3919219 |
| 315274at7742 | coiled-coil domain-containing protein 77 | S-M | 1.206626e-54 | -1.3604567 |
| 239759at7742 | protein POF1B | S-A | 1.066391e-32 | 2.7169599 |
| 305356at7742 | Uricase | S-A | 2.202899e-66 | 1.71220659 |
| 17781at7742 | Thyroid hormone receptor interactor 11 | S-A | 9.082236e-34 | 1.3541841 |
| 170533at7742 | Highly divergent homeobox | S-A | 1.378068e-25 | 1.2181134 |
| 164002at7742 | RNA recognition motif domain | S-A | 1.053009e-29 | 1.0971664 |
| 419694at7742 | Protein of unknown function DUF4693 | S-A | 1.055696e-12 | 1.0861566 |
| 237715at7742 | serine/threonine-protein kinase 17B | S-A | 4.789620e-52 | 1.0731682 |
| 238395at7742 | inward rectifier potassium channel 13 | S-A | 1.194897e-58 | -1.3048945 |
| 266521at7742 | C2 domain | S-A | 2.112682e-17 | -1.1529012 |
| 239759at7742 | protein POF1B | M-A | 5.680612e-50 | 2,2318501 |
| 315274at7742 | coiled-coil domain-containing protein 77 | M-A | 6.925240e-43 | 1,8207163 |
| 17781at7742 | Thyroid hormone receptor interactor 11 | M-A | 7.455289e-38 | 1,4061053 |
| 253823at7742 | Protein moonraker | M-A | 6.733202e-34 | 1,2771096 |
| 325663at7742 | Methyl-CpG DNA binding | M-A | 1.526045e-17 | 1,0379213 |
| 258449at7742 | mitochondria-eating protein | M-A | 2.734615e-49 | 1,0258442 |
| 237266at7742 | Charged multivesicular body protein 7 | M-A | 1.700895e-27 | -1.0513878 |
| 45090at7742 | Limbin | M-A | 6.312410e-32 | -1.0186463 |

### S2 Table. Data collection and refinement statistics of X-ray crystallography.

| Data collection | | | | |
|---|---|---|---|---|
| Data set | GgUOX-AZA reducing conditions (TCEP-HCl) | GgUOX-AZA oxidizing conditions ($H_2O_2$) | GgUOX-AZA reducing conditions (TCEP-HCl) | GgUOX-AZA oxidizing conditions ($H_2O_2$) |
| Beam Line | I24 (DLS) | I04 (DLS) | I24 (DLS) | I04 (DLS) |
| Wavelength (Å) | 0.97950 | 0.97948 | 0.97950 | 0.97948 |
| Temperature (K) | 100 | 100 | 100 | 100 |
| Resolution range (Å) Highest resolution bin (Å) | 81.502-1.71 (1.74-1.71) | 77.06-1.86 (1.89-1.86) | 66.88-2.12 (2.16-2.12) | 78.89-1.89 (1.92-1.89) |
| Space group | ***C*222₁** → $C222_1$ | $C222_1$ | $P2_12_12_1$ | $P2_12_12_1$ |
| Cell dimensions (Å) | 107.19, 125.47, 238.58 | 106.51, 125.99, 240.68 | 102.22, 124.40, 125.77 | 101.77, 124.86, 128.27 |
| Unique reflections | 169907 (8369) | 135367 (6674) | 91505 (4515) | 131041 (6365) |
| Multiplicity | 4.4 (4.0) | 13.8 (14.2) | 8.0 (8.2) | 13.5 (11.4) |
| Completeness, (%) | 98.9 (98.2) | 100.0 (100.0) | 100.0 (100.0) | 100.0 (98.1) |
| $R_{merge}$, (%) | 8.2 (141.8) | 11.6 (403.8) | 18.6 (253.9) | 17.0 (405.1) |
| $R_{pim}$ (I), (%) | 4.2 (76.8) | 3.3 (110.8) | 7.0 (93.4) | 4.8 (123.2) |
| ⟨I/σ(I)⟩ | 8.7 (0.8) | 12.1 (0.5) | 8.1 (0.9) | 7.6 (0.3) |
| CC(1/2) | 0.998 (0.345) | 0.999 (0.300) | 0.997 (0.363) | 0.998 (0.253) |
| **Refinement** | | | | |
| PDB code | 8OFK | 8OIH | 8OH8 | 8OIW |
| $R_{factor}$ (%)/$R_{free}$(%) | 18.1/20.1 | 17.2/20.6 | 18.5/21.4 | 18.9/21.4 |
| # non-H atoms | 11102 | 10860 | 10403 | 10503 |
| rms bond lengths (Å) | 0.008 | 0.008 | 0.008 | 0.008 |
| rms bond angles (°) | 1.362 | 1.483 | 1.451 | 1.414 |

**Table S3.** Sequence read archive IDs of the RNA-seq data analyzed for gene expression in *G. gallus, S. canaria, A. carolinensis.*

| Organism | Tissue | Study accession | Sample/Library | Run accession |
|---|---|---|---|---|
| *Gallus gallus* | Liver | PRJNA865899 | FPE01_liver_11 | SRR20821387_1.fastq.gz, SRR20821387_2.fastq.gz |
| | | | FPE01_liver_2 | SRR20821388_1.fastq.gz, SRR20821388_2.fastq.gz |
| | | | FPE01_liver_1 | SRR20821389_1.fastq.gz, SRR20821389_2.fastq.gz |
| | Skin | PRJNA837029 | D150-DT3 | SRR19161674_1.fastq.gz SRR19161674_2.fastq.gz |
| | | | D150-DT2 | SRR19161675_1.fastq.gz SRR19161675_2.fastq.gz |
| | | | D150-DT1 | SRR19161676_1.fastq.gz SRR19161676_2.fastq.gz |
| | Feather follicle | PRJNA494889 | A1 | SRR7973868_1.fastq.gz SRR7973868_2.fastq.gz |
| | | | A2 | SRR7973869_1.fastq.gz SRR7973869_2.fastq.gz |
| | | | A3 | SRR7973870_1.fastq.gz SRR7973870_2.fastq.gz |
| | Embryonic Skin (days 6-21) | PRJNA397795 | E6 | SRR5922808_1.fastq.gz, SRR5922808_2.fastq.gz |
| | | | E7 | SRR5922809_1.fastq.gz, SRR5922809_2.fastq.gz |
| | | | E8 | SRR5922810_1.fastq.gz, SRR5922810_2.fastq.gz |
| | | | E9 | SRR5922811_1.fastq.gz, SRR5922811_2.fastq.gz |
| | | | E10 | SRR5922812_1.fastq.gz, SRR5922812_2.fastq.gz |
| | | | E11 | SRR5922813_1.fastq.gz, SRR5922813_2.fastq.gz |
| | | | E12 | SRR5922814_1.fastq.gz, SRR5922814_2.fastq.gz |
| | | | E13 | SRR5922815_1.fastq.gz, SRR5922815_2.fastq.gz |
| | | | E14 | SRR5922816_1.fastq.gz, SRR5922816_2.fastq.gz |
| | | | E15 | SRR5922817_1.fastq.gz, SRR5922817_2.fastq.gz |
| | | | E20 | SRR5922818_1.fastq.gz, SRR5922818_2.fastq.gz |
| | | | E21 | SRR5922819_1.fastq.gz, SRR5922819_2.fastq.gz |
| | | | E16 | SRR5922820_1.fastq.gz, SRR5922820_2.fastq.gz |
| | | | E17 | SRR5922821_1.fastq.gz, SRR5922821_2.fastq.gz |
| | | | E18 | SRR5922822_1.fastq.gz, SRR5922822_2.fastq.gz |
| | | | E19 | SRR5922823_1.fastq.gz, SRR5922823_2.fastq.gz |
| *Serinus canaria* | Liver | PRJNA300534 | RED_LIV | SRR2915364_1.fastq.gz, SRR2915364_2.fastq.gz |
| | | | YEL_LIV | SRR2915372_1.fastq.gz, SRR2915372_2.fastq.gz |
| | Regenerating skin | | RED_SKIN | SRR2915352_1.fastq.gz, SRR2915352_2.fastq.gz |
| | | | YEL_SKIN | SRR2915371_1.fastq.gz, SRR2915371_2.fastq.gz |
| *Anolis carolinensis* | Liver | PRJNA78917 | B48.Liver_RNA | SRR391651_1.fastq.gz, SRR391651_2.fastq.gz |
| | | | B48.Liver_RNA | SRR391653_1.fastq.gz, SRR391653_2.fastq.gz |
| | | | B48.Liver_RNA | SRR391656_1.fastq.gz, SRR391656_2.fastq.gz |
| | Regenerating tail (section 1-5) | PRJNA253971 | S1-D015 | SRR1502164_1.fastq.gz SRR1502164_2.fastq.gz |
| | | | S2-D015 | SRR1502165_1.fastq.gz SRR1502165_2.fastq.gz |
| | | | S3-D015 | SRR1502166_1.fastq.gz SRR1502166_2.fastq.gz |
| | | | S4-D015 | SRR1502167_1.fastq.gz SRR1502167_2.fastq.gz |
| | | | S5-D015 | SRR1502168_1.fastq.gz SRR1502168_2.fastq.gz |
| | | | S1-D026 | SRR1502169_1.fastq.gz SRR1502169_2.fastq.gz |
| | | | S2-D026 | SRR1502170_1.fastq.gz SRR1502170_2.fastq.gz |
| | | | S3-D026 | SRR1502171_1.fastq.gz SRR1502171_2.fastq.gz |
| | | | S4-D026 | SRR1502172_1.fastq.gz SRR1502172_2.fastq.gz |
| | | | S5-D026 | SRR1502173_1.fastq.gz SRR1502173_2.fastq.gz |
| | | | S1-D047 | SRR1502174_1.fastq.gz SRR1502174_2.fastq.gz |

continues…

| Organism | Tissue | Study accession | Sample/Library | Run accession |
|---|---|---|---|---|
| *Anolis carolinensis* | Regenerating tail (section 1-5) | PRJNA253971 | S2-D047 | SRR1502175_1.fastq.gz SRR1502175_2.fastq.gz |
| | | | S3-D047 | SRR1502176_1.fastq.gz SRR1502176_2.fastq.gz |
| | | | S4-D047 | SRR1502177_1.fastq.gz SRR1502177_2.fastq.gz |
| | | | S5-D047 | SRR1502178_1.fastq.gz SRR1502178_2.fastq.gz |
| | | | S1-D061 | SRR1502179_1.fastq.gz SRR1502179_2.fastq.gz |
| | | | S2-D061 | SRR1502180_1.fastq.gz SRR1502180_2.fastq.gz |
| | | | S3-D061 | SRR1502181_1.fastq.gz SRR1502181_2.fastq.gz |
| | | | S4-D061 | SRR1502182_1.fastq.gz SRR1502182_2.fastq.gz |
| | | | S5-D061 | SRR1502183_1.fastq.gz SRR1502183_2.fastq.gz |

**Table S4.** Sequence read archive IDs of the RNA-seq data analyzed for gene expression in Chelonians.

| Organism | Tissue | Study accession | Sample/Library | Run accession |
|---|---|---|---|---|
| *Chelonia mydas* | Kidney | PRJNA449022 | 27K4h | SRR10020354_1.fastq.gz SRR10020354_2.fastq.gz |
| | | | chKINB | SRR10020361_1.fastq.gz SRR10020361_2.fastq.gz |
| | | | chKINA | SRR10020362_1.fastq.gz SRR10020362_2.fastq.gz |
| | | | yuLKHdna | SRR12153482_1.fastq.gz SRR12153482_2.fastq.gz |
| | Skin | PRJNA672698 | FP02-01 | SRR12916941_1.fastq.gz SRR12916941_2.fastq.gz |
| | | | FP12-01 | SRR12916943_1.fastq.gz SRR12916943_2.fastq.gz |
| | | | FP11-01 | SRR12916945_1.fastq.gz SRR12916945_2.fastq.gz |
| | | | FP10-01 | SRR12916947_1.fastq.gz SRR12916947_2.fastq.gz |
| | | | FP09-01 | SRR12916949_1.fastq.gz SRR12916949_2.fastq.gz |
| | | | FP08-01 | SRR12916952_1.fastq.gz SRR12916952_2.fastq.gz |
| | | | FP07-01 | SRR12916954_1.fastq.gz SRR12916954_2.fastq.gz |
| | | | FP03-01 | SRR12916981_1.fastq.gz SRR12916981_2.fastq.gz |
| | | | FP01-01 | SRR12916984_1.fastq.gz SRR12916984_2.fastq.gz |
| *Pelodiscus sinensis* | Liver | PRJNA762000 | GSM5570612 | SRR15829334_1.fastq.gz SRR15829334_2.fastq.gz |
| | | | GSM5570613 | SRR15829335_1.fastq.gz SRR15829335_2.fastq.gz |
| | | | GSM5570614 | SRR15829336_1.fastq.gz SRR15829336_2.fastq.gz |
| | | | GSM5570615 | SRR15829337_1.fastq.gz SRR15829337_2.fastq.gz |
| | | | GSM5570616 | SRR15829338_1.fastq.gz SRR15829338_2.fastq.gz |
| | | | GSM5570622 | SRR15829344_1.fastq.gz SRR15829344_2.fastq.gz |
